# Complexity and ultrastructure of infectious extracellular vesicles from cells infected by non-enveloped virus

**DOI:** 10.1101/618306

**Authors:** Jie E. Yang, Evan D. Rossignol, Deborah Chang, Joseph Zaia, Isaac Forrester, Holly Saulsbery, Daniela Nicastro, William T. Jackson, Esther Bullitt

## Abstract

Enteroviruses support cell-to-cell viral transmission prior to their canonical lytic spread of virus. Poliovirus (PV), a prototype for human pathogenic positive-sense RNA enteroviruses, and picornaviruses in general, transport multiple virions *en bloc* via infectious extracellular vesicles secreted from host cells. Using biochemical and biophysical methods we identify multiple components in these secreted vesicles, including PV virions; positive and negative-sense viral RNA; essential viral replication proteins; ribosomal and regulatory cellular RNAs; and numerous host cell proteins, such as regulators of cellular metabolism and structural remodeling. Using cryo-electron tomography, we visualize the near-native three-dimensional architecture of secreted infectious extracellular vesicles containing both virions and a unique mat-like structure. Based on our biochemical data (western blot, RNA-Seq, and mass spectrometry), these mat-like structures are expected to be comprised of unencapsidated RNA and proteins. Our data show that, prior to cell lysis, non-enveloped viruses are secreted within infectious vesicles that also transport viral and host RNAs and proteins.

**Importance:** The family of picornaviridae is comprised of small positive-sense RNA viruses, many of which are significant human pathogens. Picornaviruses exploit secreted extracellular vesicles for cell-to-cell viral transmission without cell lysis, and poliovirus serves as a model system for picornaviruses that are not protected by a surrounding membrane (non-enveloped viruses). The structure and contents of these vesicles secreted by virus-infected cells are described here. In addition to mature virions, these vesicles carry negative-sense, ‘template’ viral RNA and essential replication proteins, as well as cellular resources from the host. Their complex contents may comprise an enhanced virulence factor for propagation of infection, and understanding their structure and function is helping elucidate the mechanism by which extracellular vesicles contribute to the spread of non-enveloped virus infection.

## Introduction

Enteroviruses are responsible for many widespread human diseases, including poliomyelitis (poliovirus; PV), hand-foot-and-mouth disease (Coxsackievirus; CV), and a recent respiratory infection outbreak in the United States (enterovirus D68). PV has been intensively studied for over 60 years (Racaniello, 2006) as a model system for studying infection by non-enveloped, positive-sense (+) single-stranded RNA viruses. In the typical life cycle, PV enters host cells, releases its (+) viral RNA (vRNA) from the capsid, and hijacks the host cell machinery to initiate viral protein translation. In combination, viral (e.g. viral proteins 2BC, 2C, 3AB, 3A) and host cell proteins induce intracellular membrane production and remodeling for construction of viral replication complexes, where template negative-sense (-) vRNA is generated (Bienz et al., 1992; Rossignol et al., 2015). These replication “factories” are essential for the massive generation of virions that then exit the cell to infect new hosts.

As important as replication, virion exit is a critical aspect for the spread of viral infection. Enveloped viruses are surrounded by a viral membrane, providing an elegant mechanism to enter and exit host cells through membrane fusion of the viral and host cell membranes, and budding from the host cell plasma membrane, respectively (Welsch et al., 2007). In contrast, the mechanism by which non-enveloped (naked) virions cross the cell membrane barrier is less well understood. The predominant exit strategy for non-enveloped viruses was thought to be host cell lysis, but there is now mounting evidence that non-enveloped virus exit is not as different from that of enveloped viruses as was previously thought (Feng and Lemon, 2014). Enteroviruses can exit cells non-lytically through “unconventional secretion” of vesicles (e.g. (Feng et al., 2013; Bird et al., 2014)). This alternative pathway provides cell-to-cell transmission of infection through *en bloc* virion transportation. Examples of non-enveloped viruses mediating non-lytic viral spread through secreted vesicles include hepatitis A virus, Coxsackievirus and PV (Feng et al., 2013; Bird et al., 2014; Robinson et al., 2014; Chen et al., 2015). These studies on non-lytic spread have demonstrated that vesicles isolated from the media of infected cells are sufficient to infect new cells. However, the structure, content and any additional roles of these extracellular vesicles in the spread of infection have not yet been well-characterized.

Two-dimensional transmission electron microscopy (TEM) of negatively stained samples has provided insights into morphological features of exosomes released from uninfected (e.g. (Jung and Mun, 2018)) and from virus-infected cells (Feng et al., 2013; Robinson et al., 2014). The three-dimensional (3-D) structural features of native extracellular vesicles from virus-infected cells have not been defined, nor has there been a comparison between vesicles from infected and uninfected cells. Plunge-freezing allows near-native preservation of biological samples, which can then be imaged by cryo-electron microscopy (cryo-EM) for two-dimensional data, and by cryo-electron tomography (cryo-ET) to reveal 3-D reconstruction of pleiomorphic structures at nanometer to near-atomic resolution (e.g. ((Nicastro et al., 2000; Wan et al., 2017)).

Biochemically and structurally, we analyzed secreted vesicles from PV-infected cells for proteomic and RNA content, probed for cellular pathways and viral markers, and visualized their 3-D ultrastructure by cryo-ET. We describe the complex composition of extracellular vesicles released from infected cells, including considerable differences from counterpart vesicles released from uninfected cells. We show the presence of cellular markers from multiple cellular pathways, including the endosomal and autophagic pathways. Rather than the secreted vesicles containing only virions to be transported *en bloc*, PV-induced extracellular vesicles act as infectious units that deliver unencapsidated (+) vRNA and (-) vRNA, essential viral replication proteins, host proteins, and host RNAs to new cells. In summary, PV infection propagated by secreted vesicles delivers viruses and a diverse set of both viral and cellular RNA and proteins.

## Results

Extracellular vesicles serve multiple roles for cells, including cell signaling and transport of functional proteins and coding or non-coding RNAs (Gould et al., 2003; Kalra et al., 2016; Valadi et al., 2007). Vesicles with a diameter of 100-1000 nm secreted from PV-infected cells were shown to carry PV virions (Chen et al., 2015). Therefore, we analyzed extracellular vesicles isolated by size using the established fractionation method of differential centrifugation for microvesicles (100-1000 nm) (Lane et al., 2017) and for exosomes (40-100 nm) (Li et al., 2017). Phospatidylserine-containing microvesicles and CD9-positive exosomes were purified for additional analysis, as diagrammed in Figure 1A (see also Materials & Methods).

**Fig. 1.**
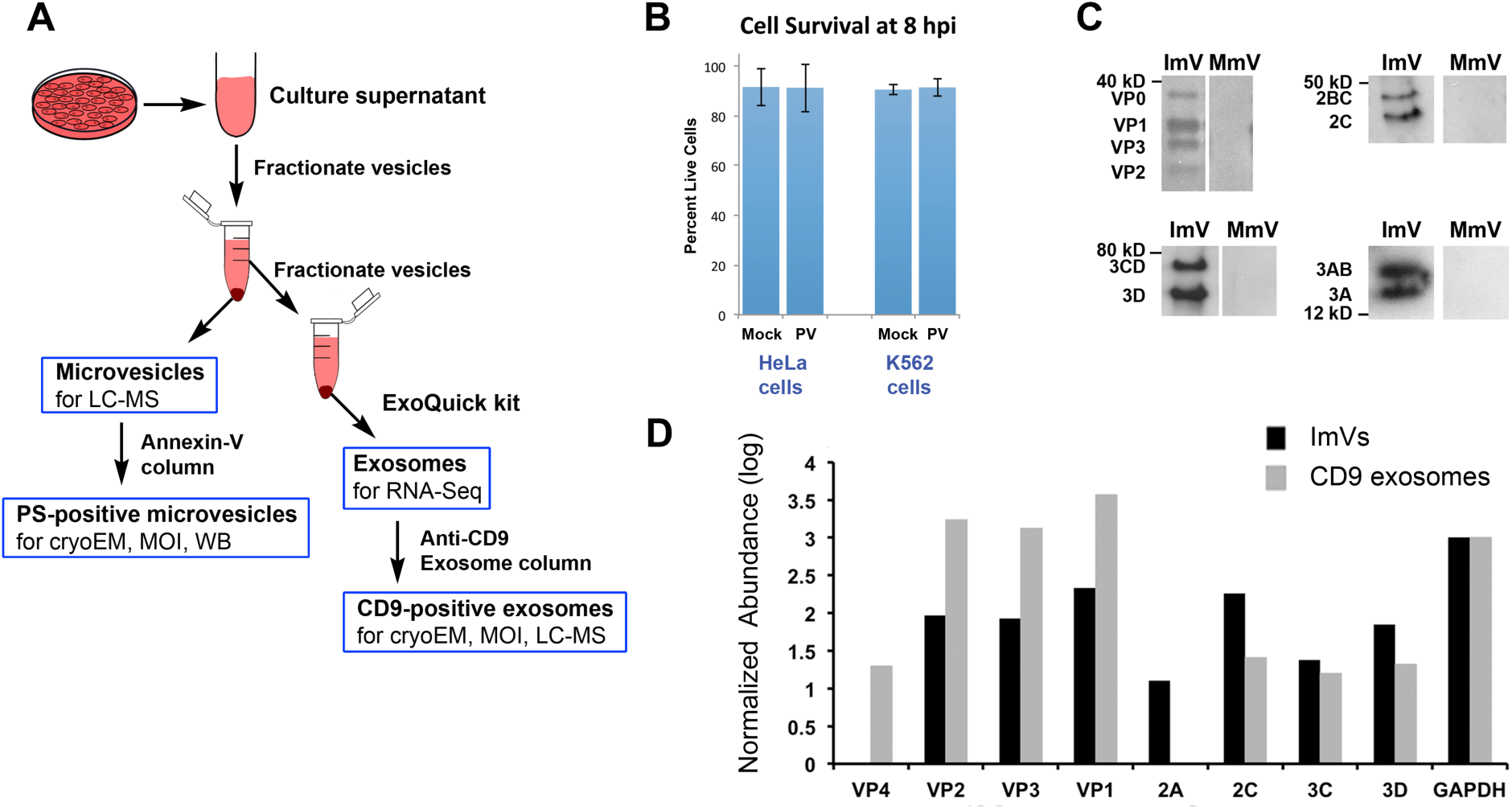
Sample preparation and the presence of PV proteins in secreted vesicles. (A) Schematic of collection and purification of microvesicles and exosomes. Annexin-V-coated magnetic beads were used to purify microvesicles with PS in their outer membrane. (B) Cell viability at the time of vesicle collection. (C) Mass spectrometric analysis of poliovirus protein abundance of infectious microvesicles and infectious CD9-positive exosomes collected at 8 hpi. The protein abundance was normalized to the host GAPDH level of PV-infected cells, multiplied by 1000 and presented as log_10_(normalized abundance). (D) Viral structural and non-structural proteins identified in PS-containing infectious microvesicles (ImVs) detected via western blot, 8 hpi.

To follow PV-infection of HeLa cells and determine efficiency of viral infection, control (mock-infected) and PV-infected cells were harvested at 1, 3, 5, and 8 hours post infection (hpi), and cytoplasmic lysates were probed for viral proteins. The non-structural viral protein 3CD was produced by PV-infected HeLa cells as early as 1 hpi, and its levels stayed constant from 3 to 8 hpi; production of the viral polymerase 3Dpol started by 3 hpi and reached a high level by 5 hpi (western blot data, not shown), consistent with the literature (e.g. (Castrillo et al., 1987)).

### Capsid proteins and essential viral replication proteins were present within extracellular vesicles secreted from PV-infected cells

To characterize the vesicles and their contents, we collected and size-fractionated secreted vesicles from infected and mock-infected cells (Fig. 1A). Because the release of PV-containing infectious microvesicles peaks at approximately 8 hpi (Chen et al., 2015), at which time infected cells have not yet begun to lyse (Bird et al., 2014; Chen et al., 2015)), vesicles were collected at 8 hpi. To rule out the presence of significant contaminants from intracellular contents in this study, cell viability greater than 90% was confirmed for both mock- and virus-infected cells (Fig. 1B). Samples included all vesicles secreted after the cells were washed and non-FBS-containing media was added at 4 hpi, the time of peak replication. Liquid chromatography-mass spectrometry (LC-MS) of 100-1000 nm diameter vesicles (called here ‘microvesicles’) secreted from infected and mock-infected cells resulted in the detection of the non-capsid, essential viral replication 2A-, 2C-, 3C- and 3D-containing proteins, in addition to viral capsid proteins VP1-VP3 (Fig. 1C). Consistent with the LC-MS data from size-fractionated microvesicles, western blot analyses (Fig. 1D) further confirmed the presence of non-capsid viral proteins in microvesicles that contained the membrane phospholipid phosphatidylserine (PS), a sub-population of microvesicles that had been shown to be infectious (Chen et al., 2015). Specifically, the PV proteins 2BC, 2C, 3A, 3AB, 3CD, and 3D were identified in PS-containing microvesicles secreted from PV-infected cells but not those secreted from mock-infected cells (Fig. 1D).

### Host cell proteins identified by mass spectrometry analysis

Our data showed that less than 10% of the LC-MS identified protein peptides in microvesicles from PV-infected cells were viral proteins. Studying the components in more detail, we observed that the host cell protein components in microvesicles from mock-infected cells exhibited a much smaller diversity of proteins than infectious microvesicles from PV-infected cells (Table S1A-B). We identified a total of 168 host cell protein matches in infectious microvesicles, with 65 proteins (38%) identified in both independent runs. There were three technical replicates per run and a threshold cutoff of 0.999 probability. Of these proteins, 11 were identified in both ImV repeats and in the MmV sample (17% of the proteins found in both ImV expts), out of a total of 12 identified in either ImV and the MmV samples (7% of the proteins identified in at least one ImV experiment were present in the mock-infected vesicles).

We then categorized functionally-related proteins using Ingenuity Pathway Analysis (IPA; Qiagen) software (Krämer et al., 2014). The significant pathways (-log p > 3), as calculated by the p-value of overlap (see Materials and Methods), are shown in Table 1. Seven enzymes from the glycolytic pathway were identified in infectious microvesicles (PGK1, ENO1 {ENOA}, GPI {G6PI}, TPI {TPIS}, PKM {KPYM}, ALDOA, GAPDH); names in braces are synonyms), as were 12 proteins from the RhoGDI (Rho GDP dissociation inhibitor) pathway (ITGB1 {CD29}, GNAI3, GNAS, MYL6, CFL1, ACTB, EZR, RHOA, RAC1, CD44, GNG12, MSN). Proteins identified in infectious microvesicles are also involved in both caveolar-mediated and clathrin-mediated endocytic virus entry pathways (ITGB1, B2M, FYN, CD55 {DAF}, HLA-A, FLNA, ACTB, RAC1, TFRC {CD71}, FOLR1), which include the enterovirus 70 protein receptor (CD55, also known as the CV A21 host-entry receptor DAF) and the echovirus receptor Integrin beta 1 proteins for caveolar-mediated endocytosis (Newcombe et al., 2004; Karnauchow et al., 1996; Jokinen et al., 2010), and the host receptor TfR1 protein that is utilized by the enveloped arenavirus for entry through clathrin-coated endocytosis (Flanagan et al., 2008). Another pathway identified with high probability (-log p = 8.25) was actin cytoskeleton reorganization, including RAC proteins (Rho family GTPases), the tyrosine-protein kinase FYN, and additional actin cytoskeleton-regulating proteins (ITGB1, PFN1, CFL1, MYL6, FLNA, RHOA, EZR, RAC1, TMSB10/TMSB4X, IQGAP1, GNG12, MSN).

**Table 1.**
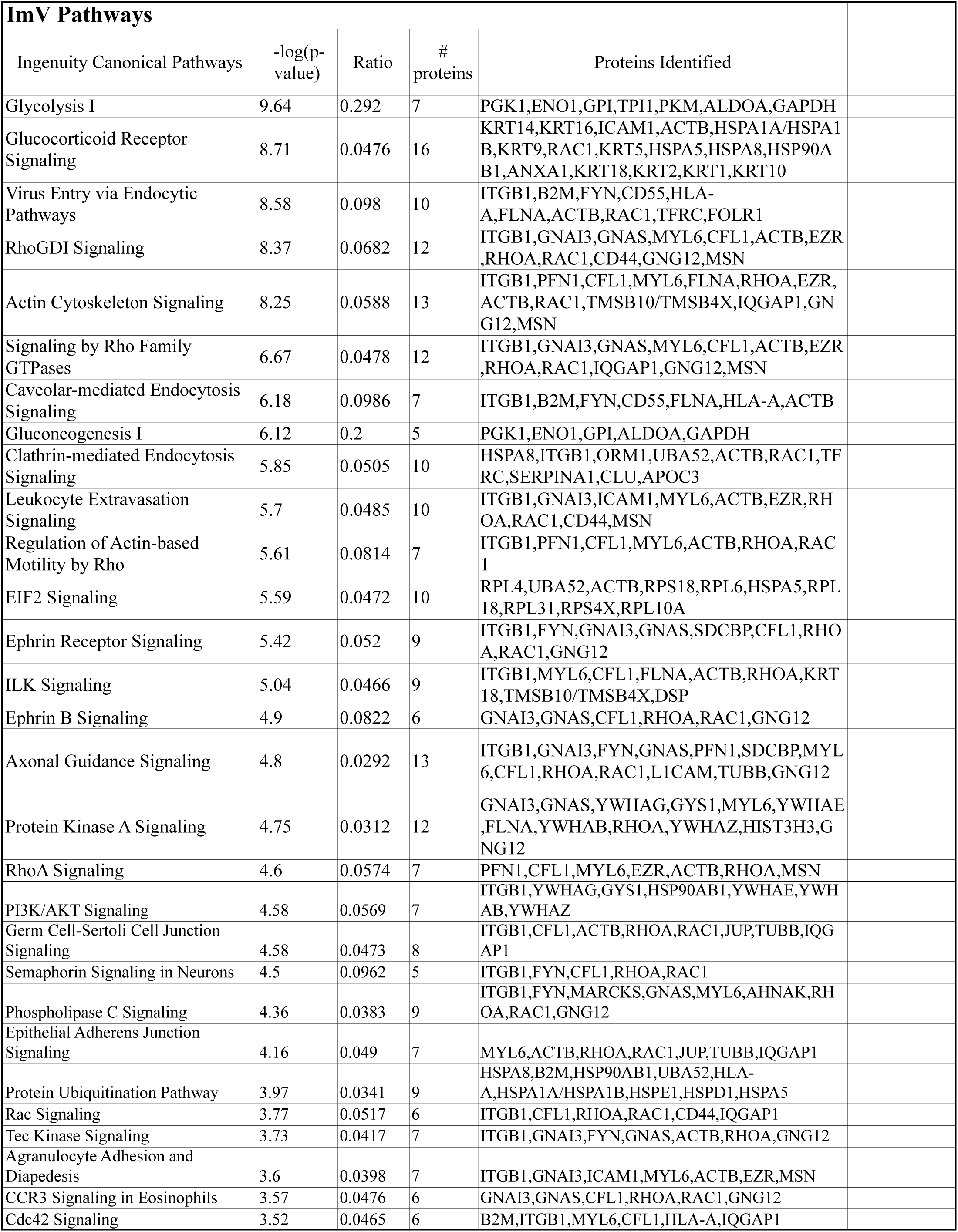
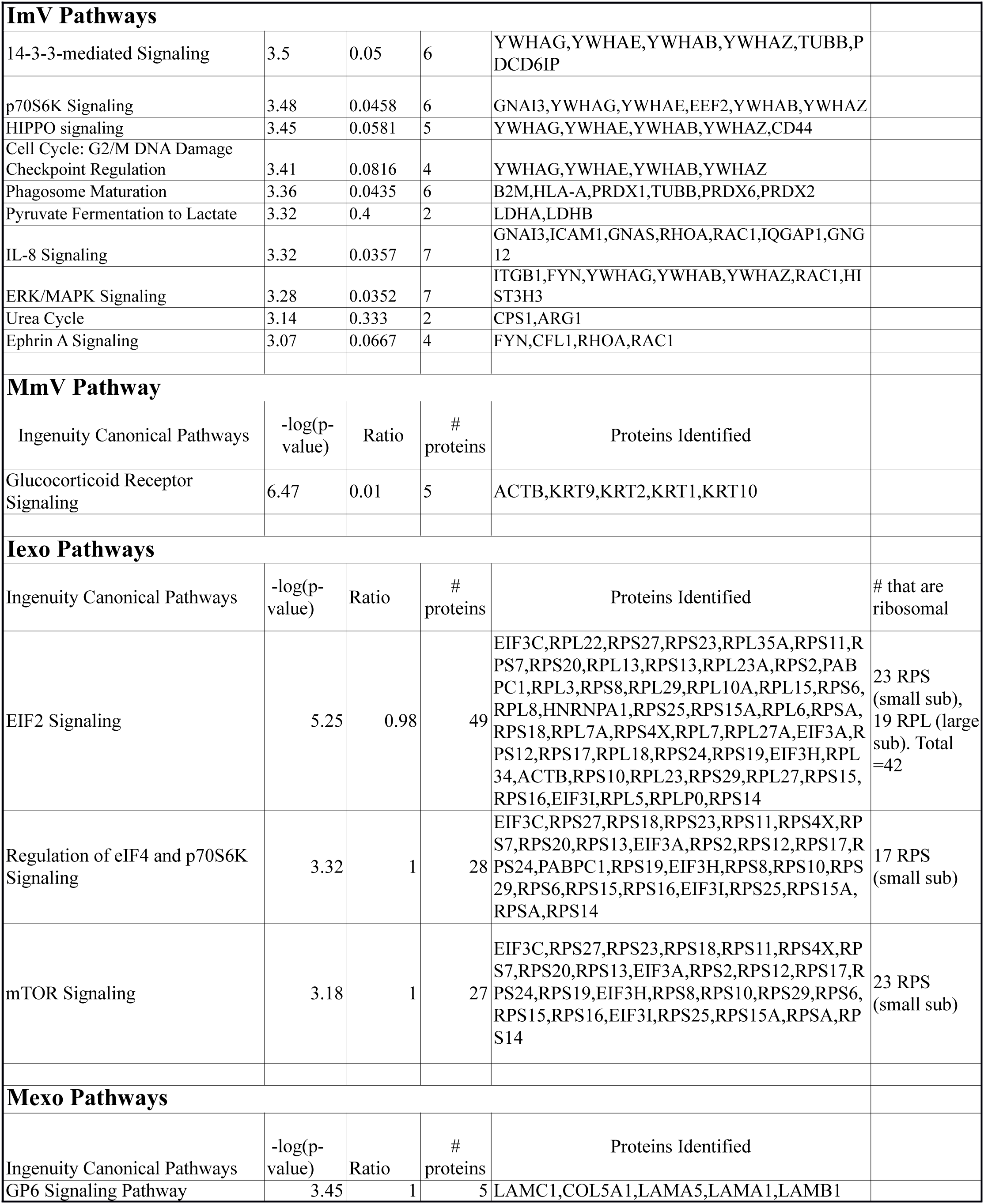
Pathways identified by the significant presence of functionally-related proteins in secreted vesicles, based on LC/MS data (-log p >3).

Although exosomes are typically smaller sized vesicles, some overlap between microvesicles and exosomes can be expected from a differential centrifugation separation (Kanada et al., 2015; Raposo and Stoorvogel, 2013). Therefore, it is not surprising that components of the exosomal pathway were also identified in our size-fractionated infectious microvesicles. These components include PDCD6IP (programmed cell death 6 interacting protein, also known as ALIX), which participates in ESCRT-III recruitment (Schmidt and Teis, 2012), syndecan binding protein (SDCBP), which is involved in the biogenesis and cargo loading of exosomes (Friand et al., 2015), and several classical exosomal markers such as the ESCRT-III associated factor, increased sodium tolerance 1 (IST1) (Allison et al., 2013) and CD9 (Andreu and Yáñez-Mó, 2014) (Table S1A).

The detection of these exosomal components led us to question whether PV infection might also exploit smaller exosome-like vesicles to transport virions and viral proteins from cell to cell, as has been shown for exosome-like virion-containing vesicles from hepatitis A virus-infected cells (Feng et al., 2013). To investigate their role in PV spread, exosomes shed from mock- and PV-infected cells were collected, using established size-based fractionation of 40-100 nm diameter vesicles (see Materials and Methods). For experiments where it was essential to avoid contamination of the exosome fraction with similarly sized free virions (28 nm diameter) and microvesicles, i.e. for LC-MS, cryo-ET, and functional infectivity characterization (plaque assays), the exosome fraction was further purified using antibodies against the exosomal marker CD9 (Fig. 1A). LC-MS data from CD9-positive exosomes secreted from PV-infected cells showed that 9% of detected protein peptide varieties were derived from PV proteins, both capsid (VP) and non-capsid viral proteins (2C-, 3C-, and 3D-containing proteins) (Fig. 1C), similar to the results from infectious microvesicles. The absence of 2A-containing peptides is noted as likely a technical absence, since the elution time of 2A corresponded with a high abundance of background peptides that could prevent detection of 2A (data not shown). In the infectious microvesicle sample, where 2A was detected, the 2A peptide had a high enough relative abundance compared to background to be detected by tandem MS. As seen in Table 1, exosomes from infected-cells contained numerous host proteins associated with pathways for 1) eIF2 signaling/translation initiation/protein translation that include RNA-binding and ribosomal proteins, ribonucleoproteins, and tRNA synthetase complex components, 2) eIF4 and p70S6K signaling, and 3) mTOR signaling. Conversely, CD9-positive exosomes from mock-infected cells only met the stringent pathway criterion for protein matches in the GP6 signaling pathway (Table 1, and see Table S1 for a complete list of protein matches above a stringent cut-off).

As exosome-sized vesicles from PV-infected cells were not previously known to be involved in non-lytic PV transmission, we compared the protein contents of microvesicles and exosomes. As shown in Table S1, we identified 36 host cell protein matches present in all infectious vesicle samples, including proteins involved in negative regulation of apoptosis (e.g. YWHAZ). Exosomes from PV-infected cells matched to 100 proteins that were not identified in microvesicles from infected cells (Table S1). Although three proteins from the glycolysis pathway (PKM, ALDOA, and GAPDH) were identified in infectious exosomes, and one (GAPDH) was also identified in exosomes from mock-infected cells, in neither case did this pathway meet the criteria for significance. The majority (55%) of the proteins only identified in exosomes were ribosomal proteins; these ribosomal proteins comprised 34% of all proteins identified in infectious exosome samples.

### Template and genomic RNA were present in secreted infectious microvesicles

The initiation of PV replication for packaging (+) vRNA within virions requires the presence of template (-) vRNA. Therefore, we examined the content of PS-positive infectious microvesicles with regard to the presence of vRNA by RT-PCR (Materials & Methods). To confirm the absence of DNA contamination from the total RNA extraction/purification process, SuperScript III Reverse Transcriptase was omitted from the “master mix” for reverse transcription, and no amplification occurred in total RNA from PV-infected cells, when positive-(genomic) /negative-sense (anti-genomic) viral RNA was probed (data not shown). To test for nonspecific binding introduced by primer dimers and secondary structures of the primers, RT-qPCR was performed on RNase/DNase-free water including primers either against positive-/negative-sense RNA. No amplicons were detected above the threshold that was set at a fixed intensity (0.475) for all experiments (data not shown). In addition to these system controls, we analyzed melt curves to test whether the dye qPCR assay (SYBER) produced single, specific products. The appearance of single peaks indicated one melting event, corresponding to the (positive or negative) target amplicon (data not shown). We found significant decreases (*p* ≤ 0.001) in the cycle threshold, C_t_ (number of PCR cycles for the signal to exceed background) for PV (+) vRNA (viral genomic RNA that can also be used for both translation and as a template for (-) RNA synthesis) and (-) vRNA (viral template RNA) within infectious microvesicles in comparison to non-specific amplification from mock-infected microvesicles (Fig. 2A).

**Fig. 2.**
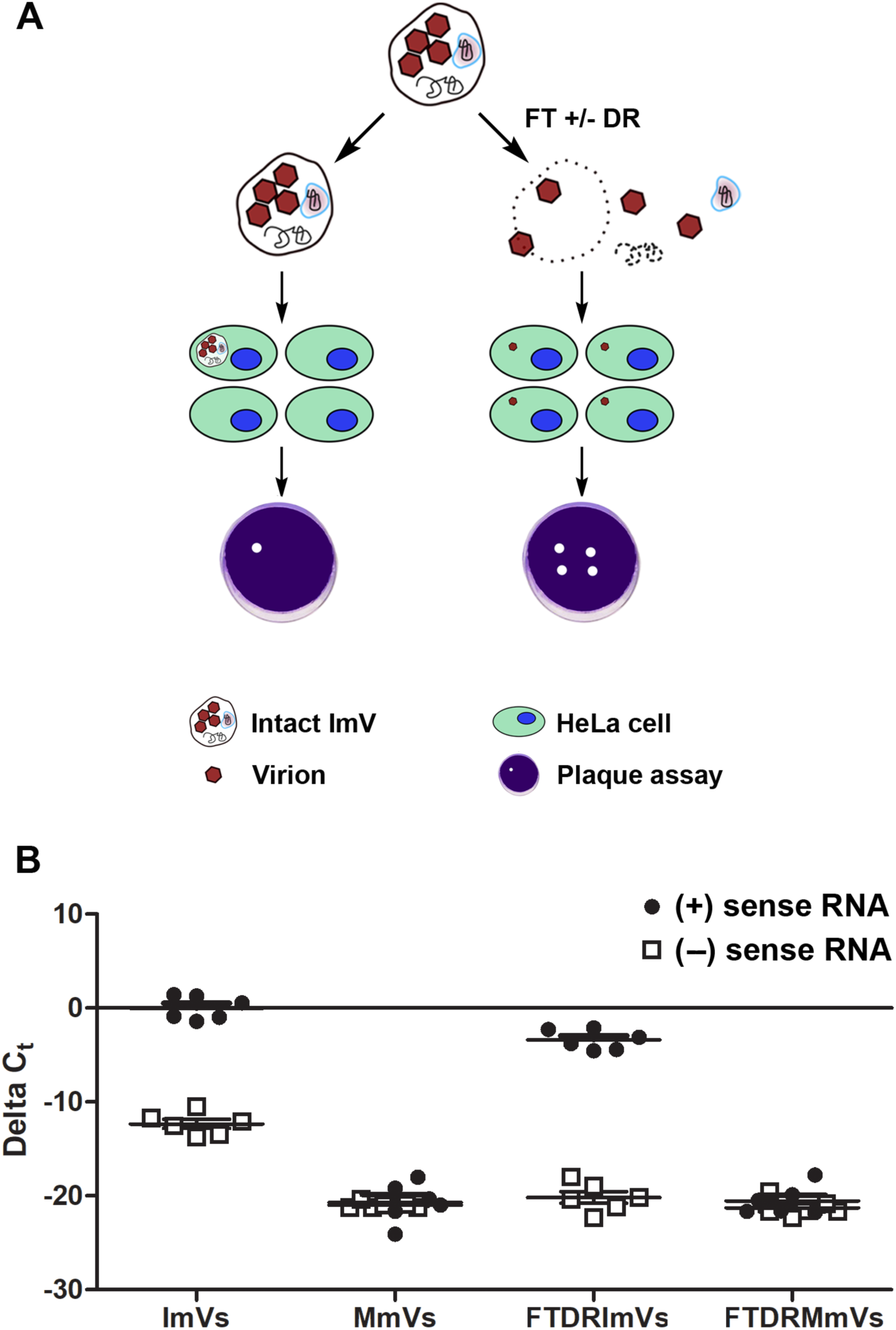
Infectious microvesicles (ImV) carry viral RNAs. (A) Schematic of experimental design for determining multiplicilty of infection of untreated infectious microvesicles (ImVs) and from ImVs treated by either freeze/thaw alone (FT) or FT, detergent, and RNase (DR). Plaque assays are used to determine the number of visible infectious sites (where cells have died) using serial dilutions of ImVs. (B) RT-qPCR showed the amounts of both (+) and (-) sense viral RNA (vRNA) from infectious microvesicles (ImV) and mock microvesicles (MmV) that were collected from supernatant at 8 hpi. vRNA was measured after extracellular vesicles underwent: 1) no treatment (ImV/MmV), or 2) freeze-thaw & detergent (1% sodium deoxycholate) & RNase treatment (DRImV/DRMmV) to break open vesicles and degrade unencapsidated RNAs. RT-qPCR relative quantification was calculated as ΔC_t_ where ΔC_t_ = (C_t_ of endogenous control gene (GAPDH)) – (Ct of gene of interest (vRNA)), using GAPDH of whole cells for normalization [].

The vRNA within secreted vesicles had been expected to be entirely virion-encapsidated vRNA (Altan-Bonnet, 2016; Chen et al., 2015). However, virion assembly is tightly regulated to encapsidate only (+) vRNA (Novak and Kirkegaard, 1991). We therefore tested whether the newly identified intravesicular (-) vRNA was “free”, or was packaged within virion capsids that were inside infectious microvesicles. We exposed microvesicles prepared as diagrammed in Figure 1A to a series of further treatments (Fig. 2B) that included membrane disruption to release the vesicle content by freeze-thaw and detergent (sodium deoxycholate), followed by RNase treatment to remove any unprotected (intravesicular yet unpackaged in capsids) RNA. Because membranes are disrupted, and capsids are resistant to and unaffected by freeze-thaw, detergent, or RNases (e.g. (Mandel, 1973; Fenwick and Cooper, 1962; Bishop and Koch, 1969)), the majority of (+) vRNA was still present in post-treated infectious microvesicles, as expected, whereas (-) vRNA was almost undetectable (C_t_ ≥ 37, a value determined based on the negative reverse transcription and non-template controls) (Fig. 2C). Interestingly, however, there was a small but significant decrease (*p* ≤ 0.05) in the abundance of (+) vRNA in post-treated infectious microvesicles. These data indicate that not all (+) vRNA was protected inside assembled virions, as capsids are well known to protect their internal RNA from RNase-mediated degradation (Bishop and Koch, 1969; Novak and Kirkegaard, 1991). The status of this RNA, whether single- or double-stranded, is not known, as under the conditions of the experiments both would be digested by RNaseA.

### Exosomes from PV-infected cells contained several classes of host cell RNAs

Previous studies demonstrated that extracellular vesicles secreted from uninfected human cells transport functional coding and non-coding RNA species (Ratajczak and Ratajczak, 2016). For an investigation of the RNA content of exosomes from infected cells, the highly purified CD9-positive exosome fraction was insufficient, as it represents only a small fraction of the whole exosome population. Therefore, exploratory RNA-Seq experiments were performed on size-fractionated exosomes secreted by K562 cells. These cells have two advantages: they are known for secreting high numbers of exosomes (Savina et al., 2002), and they can be persistently infected by PV (Lloyd and Bovee, 1993), allowing for long collection times without cell lysis, as documented in Figure 1B. Supernatants were collected from mock- and PV-infected K562 cells over a 24 h period, then subjected to size-exclusion fractionation as shown in Figure 1A. In addition to PV (+) and (-) vRNA shown in Figure 3, RNA-Seq identified over 60,000 identifiable short reads in the RNA extracted from the secreted exosomes. To interpret these data, we limited our analysis to RNA identified by at least 1000 short RNA-Seq reads, from exosomes secreted by PV-infected cells. Of the 400 host cell RNA sequences meeting these criteria, 198 were increased by at least 2-fold upon PV infection (Table 2, and Table S2 for primary data on the 400 identified RNAs). As shown in Table 2, PV infection greatly increased the relative amount of four types of RNA sequences: 1) piRNA (PIWI-interacting RNA), a class of small non-coding RNAs known for repressing transposons via translational and post-translational modifications; 2) small nucleolar RNAs, which are guide RNAs that direct methylation & pseudouridylation of mRNA; 3) ribosomal RNA (rRNA), as expected due to the abundant presence of ribosomal proteins identified in exosomes from PV-infected cells by mass spectrometry (Table S1C); and 4) tRNA, which could be present through their interactions with ribosomes.

**Table 2.**
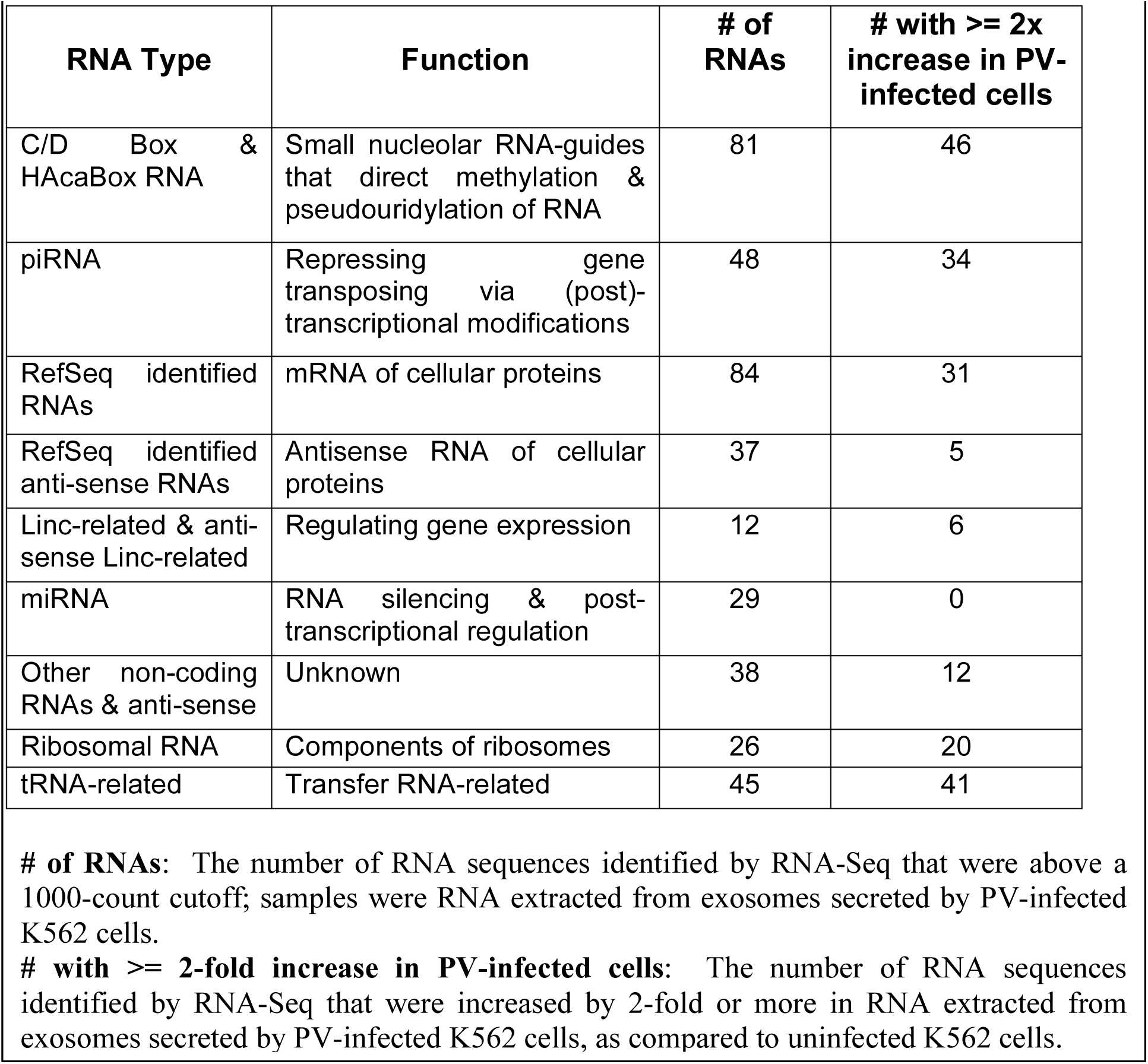
Summary of RNAs identified by RNA-Seq in exosomes from PV-infected cells

**Fig. 3.**
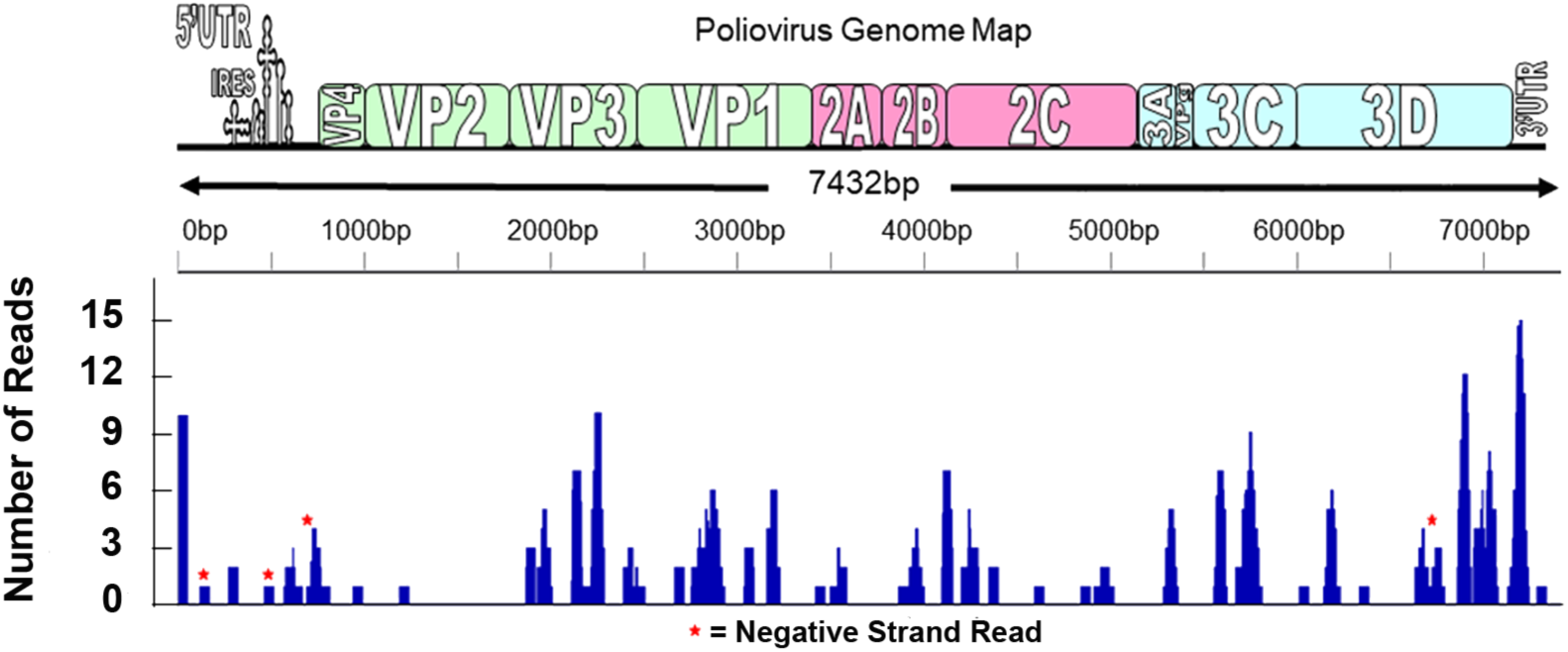
Viral RNA was identified in exploratory RNA-Seq experiments. Raw RNA-Seq data were compared to the PV Mahoney genome to determine the location and number of identified reads. Reads were spread throughout the genome, with clusters at the 5’ and 3’ ends and fewer reads at the 5’ end of the polyprotein-coding region. While the majority of reads correlated with the positive strand, four negative sense reads were identified (starred).

Taken together with mass spectrometry and western blot data, our findings demonstrate that PV-induced extracellular vesicles (both microvesicles and exosomes) carry multiple cargos, including viral replication machinery components, unencapsidated viral RNAs, cellular proteins and RNAs, including ribosomes, in addition to virions containing encapsidated genomic (+) vRNAs.

### Intact extracellular vesicles were infectious and accelerated initiation of vRNA replication

Previous studies demonstrated that microvesicles transport virions. Classic infectivity plaque assays showed that intact infectious microvesicles produced fewer infectious centers than intravesicular virions released from these vesicles by freeze/thaw (Chen et al., 2015). Confirming these previous results, our quantification of plaque assays revealed a 10-fold increase in plaque-forming units (pfu) after disruption of the infectious microvesicle membrane and release of intravesicular virions by freeze/thaw (FT) prior to infection (Fig. 4A, ImV *vs.* FT ImVs). We also saw a greater than 3-fold increase in pfu for infection with freeze/thawed-detergent/RNase-treated infectious microvesicles as compared to intact infectious microvesicles. We note that the number of infectious centers (pfu) was not significantly different when cells were infected with intravesicular PV virions released by either method: freeze/thaw or freeze/thaw/detergent/RNase (FTDR) (Fig. 4A, FT *vs.* FTDR). Analogous experiments were completed on CD9-positive exosomes purified from the media of PV-infected cells. Exosomes from PV-infected cells were shown to be infectious, exhibiting a three-fold increase in plaque formation after disruption of the vesicles’ membranous structures by freeze/thaw (Fig. 4B).

**Fig. 4.**
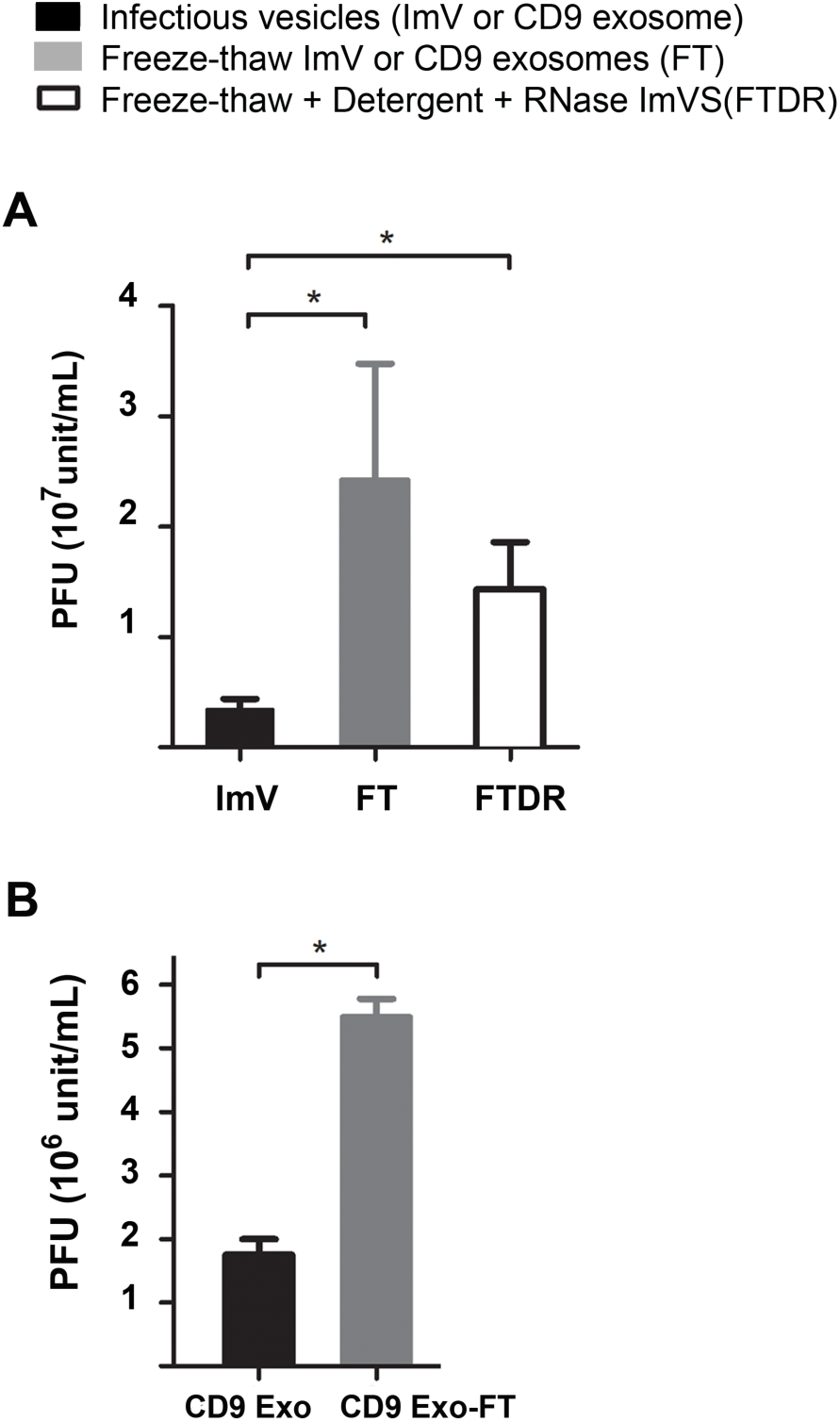
Extracellular vesicles from PV-infected cells are infectious. Infectivity of purified infectious microvesicles (ImVs; panel A) and CD9-positive exosomes (CD9 Exo; panel B) was quantified by standard plaque assay for virus titer expressed as plaque-forming unit (pfu) per mL; * P < 0.05. Samples were untreated, freeze/thawed (FT) or freeze/thawed and & detergent (1% sodium deoxycholate) & RNase treated (FTDR). Data are means ± standard deviations from at least three independent experiments. (C) Fold-change in viral RNA production from ImVs. RT-qPCR data were normalized against whole cell GAPDH, compared to a mock-infected group, and displayed as means ± standard deviations from at least three independent experiments. ** P < 0.001. Cells were infected with a constant number of virions, either transported in an intact extracellular vesicle, or as free virions after release from the vesicles, at an MOI of 1 pfu per cell.

### Ultrastructural analysis by cryo-electron tomography revealed multiple classes of infectious microvesicles

A remaining unanswered question was the 3-D structures of virus-induced extracellular vesicles that, as shown above, accommodate not only viral capsids, but also viral and cellular RNA, cellular proteins, and viral replication proteins. To visualize vesicle ultrastructure, PS-positive infectious microvesicles and purified CD9-positive exosomes secreted by PV-infected cells from 4 to 8 hpi were preserved by plunge-freezing, and then imaged by cryo-EM and cryo-ET. To enrich the infectious extracellular vesicle population for low throughput cryo-EM, we imaged PS-positive microvesicles and CD9-positive exosomes. PS-positive infectious microvesicles showed a wide size distribution, with diameters ranging from 70 nm to 820 nm. Approximately 90% of these microvesicles were 100 to 300 nm in diameter, with a median of 170 nm (n = 210 vesicles, Fig. 5A). The microvesicles displayed neither a strong uniformity in size, nor a strong correlation between size and number of included virions. 3-D reconstructions computed using cryo-ET tilt series revealed that 70% of the vesicles (n = 180 vesicles) contained virions. Quantification of virions per infectious microvesicle shows that 83% of infectious microvesicles carry 1 to 20 virions and, on average, each infectious microvesicle transports 10 virions (n = 150 vesicles from cryo-ET data, Fig. 5B).

**Fig. 5.**
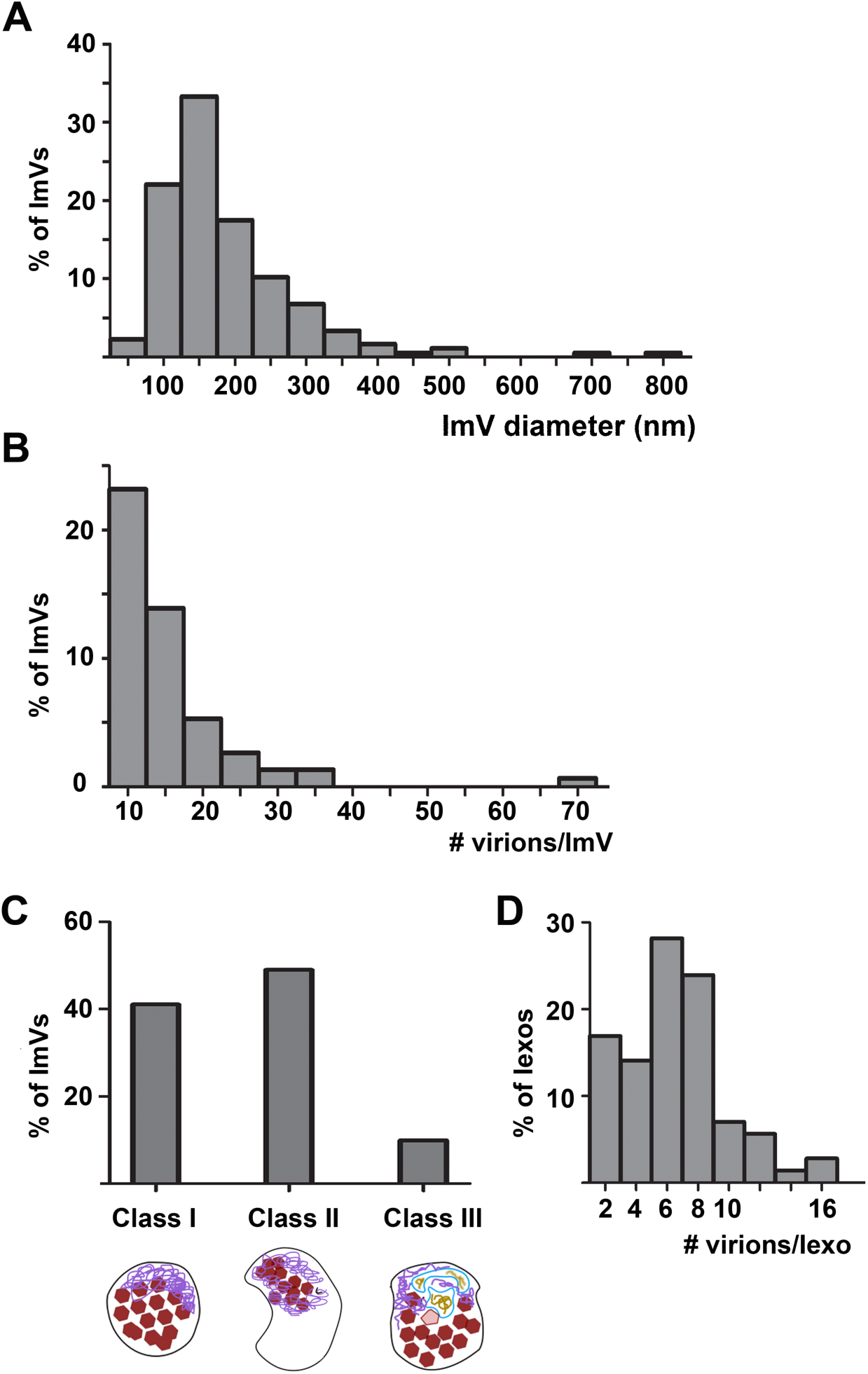
Statistics of vesicle size, virions per vesicles, and morphological classes of infectious extracellular vesicles from PV-infected cells. (A) Infectious microvesicle (ImV) diameter, (B) number of virions per ImV, (C) morphological class population, analyzed from cryoEM and cryo-tomographic data. As described in the text, and shown below panel D, Class I vesicles are close-packed with virions and additional electron-dense material; Class II vesicles are polar structures, with significant regions of low electron density in addition to virions and mat-like structures; and Class III vesicles contain the contents above as well as inner vesicular structures, and (D) number of virions per infectious exosome (Iexo).

Based on their internal morphology, we categorized infectious microvesicles into three classes (Fig. 5C). As shown in cryo-EM images in Figure 6, Class I contained densely packed virions (Fig. 6A-D’), Class II contained clustered virions and electron-lucent regions (Figs. 6E-F’), and Class III included internal vesicular structure(s) (Fig. 6G,G’). In addition to virions, all classes contained electron-dense material that resembles a mat of threads or noodles (modeled in purple in Figs. 6 and 7). As with size and number of virions per infectious microvesicle discussed above, there was no correlation between infectious microvesicle size and the packing arrangement of virions (i.e. infectious microvesicle class). For example, a relatively large infectious microvesicle with a diameter of 200 nm could have Class I morphology (Fig. 6C-D’) or Class II features (upper right vesicle in Fig. 6E,F).

**Fig. 6.**
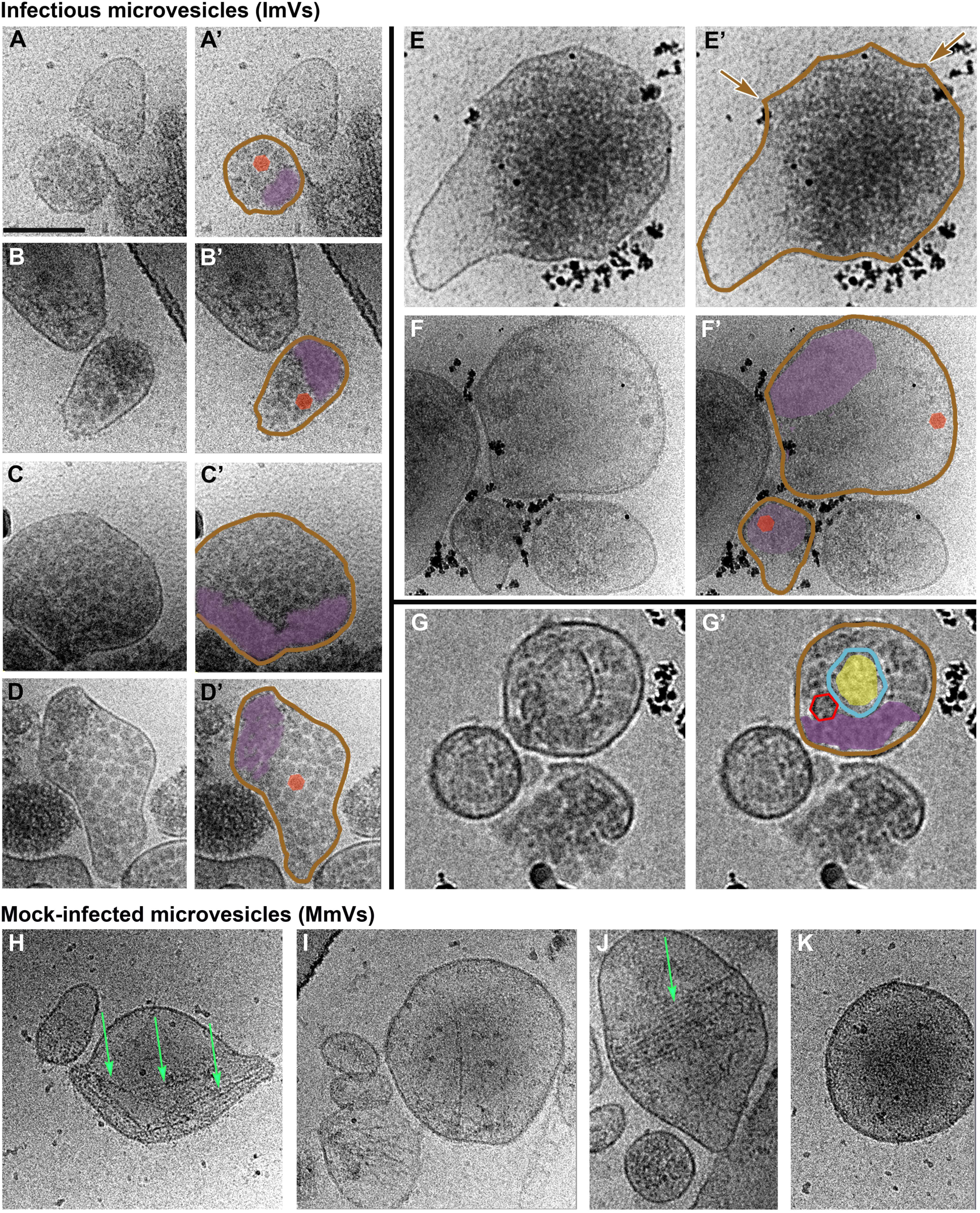
Virions and additional densities are visible in 2D cryoEM images of three morphological classes of infectious microvesicles secreted from PV-infected cells. (A-D) Class I vesicles are close-packed with virions and additional electron-dense material, termed “mat-like” structures. (E-F) Class II vesicles are polar structures, with significant regions of low electron density. (G) In addition to virions and mat-like structures, Class III vesicles contain inner vesicular structures. (H-K) Microvesicles from mock-infected cells have no virions, low internal density, and often include actin filaments (e.g., green arrows). Panels with primes (‘) highlight features including vesicle membranes (brown), virions (solid red), empty virions (traced in red, unfilled), mat-like density (purple), and inner vesicular structures (blue) with internal density (yellow). The outer membranes of infectious microvesicles often show a scalloped feature; e.g., region between brown arrows in E’. Scale bar shown in A represents 100 nm for all images. All images except G & G’ were taken at a tilt angle of 0 °. G & G’ were taken at a tilt angle of 15 °.

**Fig. 7.**
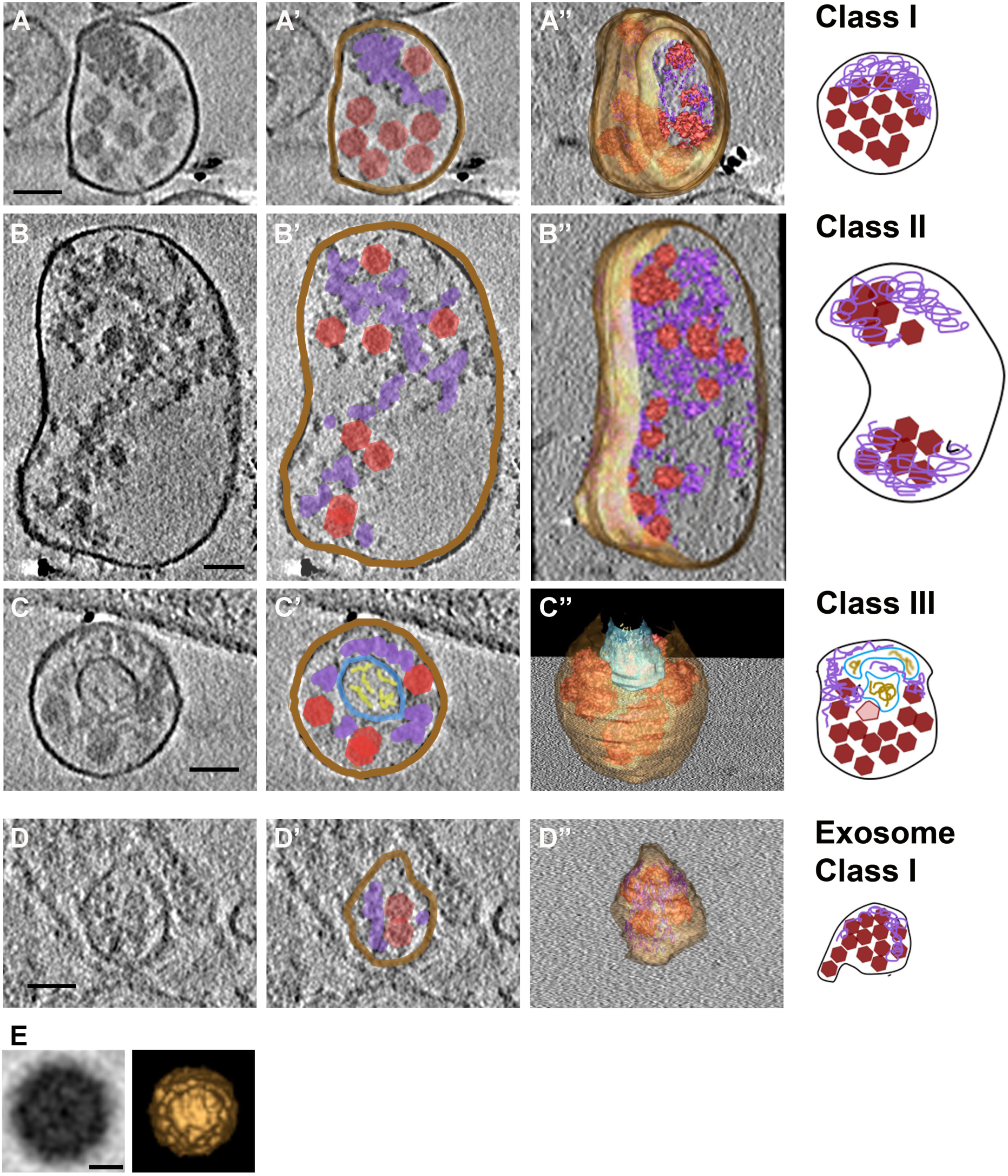
Three-dimensional data provide details and architectures of 3 classes of extracellular vesicles and virions from PV-infected cells. (A-D) Slices through tomograms computed from cryo-electron tomography (ET) tilt series of vesicles isolated at 8 hpi show details of the 3 classes from Fig. 5 and infectious exosomes. The corresponding cartoon models of the three classes are displayed on the right, and include highlighted virions (red polygons), mat-like structures (purple), and inner vesicular structures (blue/yellow). (E) Sub-volume average of virions identified inside infectious microvesicles and exosomes. (Left) A low-pass (cutoff spatial frequency of 0.028 Å) filtered tomographic slice of the sub-volume average of 116 virions inside extracellular vesicles. (Right) Isosurface rendering of the sub-volume average virion. Scale bars (A-D), 100 nm; (E) 10 nm

As expected, no virions were observed within any vesicles isolated from mock-infected (control) cells (Fig. 6H-K). In contrast to infectious microvesicles, the major recognizable structural components observed in control microvesicles were disordered or bundled actin filaments (Fig. 6H-J), which is consistent with the mass spectrometry data showing an abundance of actin and actin-binding proteins in microvesicles secreted from mock-infected cells (Table S1). In contrast, actin filaments were rarely observed within infectious microvesicles; in the few examples where infectious microvesicles contained actin filaments, multiple virions were dispersed within the bundled actin (data not shown).

Three-dimensional reconstructions were computed using cryo-ET data from PS-positive infectious microvesicles and purified CD9-positive exosomes secreted by PV-infected cells from 4 to 8 hpi. Virions and mat-like structures were observed in the lumen of the microvesicles and exosomes (Fig. 7 and Supplemental Movie). To confirm the presence of virions within the vesicles, we computed a subtomogram average of 115 structures from 1,022 particles we identified as virions, resulting in a reconstruction of the PV capsid at 6.9 nm resolution (Fig. 7E). Consistent with data from our 2-D cryo-EM images (Fig. 6), over 90% of the 3-D reconstructed infectious microvesicles showed either the Class I (Fig. 7A, deposited in the wwPDB under accession code EMD-7873) or Class II (Fig. 7B, wwPDB accession code EMD-7872) morphology, whereas Class III vesicles comprise only 10% of the population (Fig. 7C, wwPDB accession code EMD-7871, and Supplemental Movie). Infectious microvesicles were rarely spherical, and showed a single membrane enclosing an irregularly shaped structure, often with a scalloped outer contour, as seen in Figure 6E (between the brown arrows). Class II infectious microvesicles (Fig. 7B-B”), showed “empty” (electron-lucent) regions encompassing up to 90% of the infectious microvesicle volume, in addition to virions and mat-like structures. Class III infectious microvesicles contained inner vesicular structures as shown in Figures 7C-C” and Figure 8, each representing a single-membrane vesicle entrapped in the lumen of an infectious microvesicle. The average ratio of diameter of the inner vesicle to that of its infectious microvesicle carrier was 0.5 ± 0.06 (n = 17). Less dense features were observed within inner vesicles (e.g. yellow region in Fig. 7C,C’), as compared to the above-described mat-like structures of infectious microvesicles (Fig. 8, purple arrows). This suggests that structures inside the larger compartment of infectious microvesicles and those within inner vesicles may be distinct from each other. We also observed protein structures that comprise a globular ‘head’ on a stalk in the membrane of the inner vesicle, faced toward the microvesicle lumen (Fig. 8, cyan arrows).

**Fig. 8.**
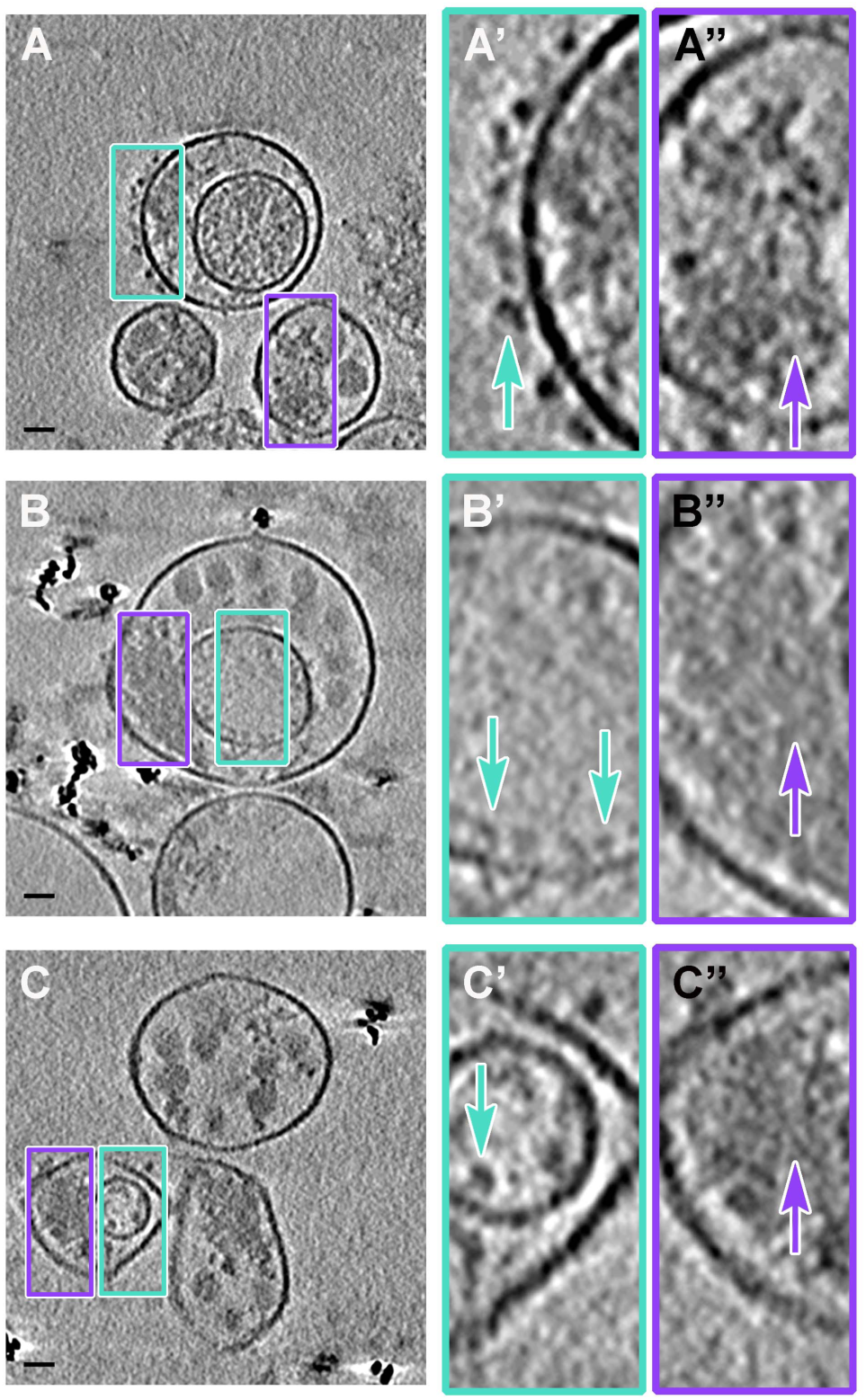
Mat-like structures are characteristics of extracellular vesicles from PV-infected cells. (A-C) Cryo-electron tomographic slices 8.2 nm thick illustrate details of infectious microvesicles; EM Databank IDs: EMD-7877, EMD-7878, EMD-7873, respectively. (A’-C’) are boxed regions of (A-C) at 3X magnification. Mat-like structures are present in all three classes of infectious microvesicles; purple arrows. Scale bars for original images, 100 nm; 33 nm for prime images (e.g. A’).

3-D structural analysis of CD9-positive exosomes secreted by PV-infected cells displayed a relatively uniform size, with an average diameter of 80 nm ± 27 nm and an average of 7 ± 3 virions per exosome (Fig. 5D; n = 69; data from nine reconstructed tomograms). The predominant morphology of exosomes from infected cells corresponded to features seen in Class I infectious microvesicles, with a densely packed interior volume (Fig. 7D-D”, and deposited in the wwPDB, accession code EMD-7879).

## Discussion

It has long been observed that non-enveloped viruses, such as those in the enterovirus genus, can propagate infection prior to lysis of host cells (Jackson et al., 2005; Lloyd and Bovee, 1993). However, the mechanism for exit (and entry) of these viruses without disruption of the cell’s plasma membrane is not well understood. Only recently has unconventional secretion of virion-containing extracellular vesicles been described (Bird et al., 2014; Chen et al., 2015; Feng et al., 2013; McKnight et al., 2017; Robinson et al., 2014). Delivery of such a ‘payload’ containing multiple virions has the potential to propagate infection that might otherwise be hindered by frequent detrimental mutations in the viral genome that are present due to the inherently error-prone replication of RNA (Biebricher and Eigen, 2005; Schulte et al., 2015). Such a diversity of virions can be achieved by either increasing the number of infecting free virions per cell (MOI) or by entry of a vesicle containing multiple virions into one cell.

Here, we have presented data showing that in addition to infectious virions, extracellular vesicles secreted from PV-infected cells contain a complex mixture of unencapsidated (+) vRNA as well as (-) vRNA, ready-made viral replication proteins, host proteins and host RNAs. In our new model, we speculate that vesicles comprising virions and macromolecules contribute to enhanced viral transmission and establishment of new infection, because in addition to virions, infected cells receive all the components necessary to begin infection *prior* to translation of the vRNA that is packaged into virions. A schematic of conventional cellular secretion (Fig. 9A) is compared to unconventional secretion, diagrammed by the interplay between viral replication and packaging of contents for non-lytic exit in Figure 9B. In this figure we designate structures in infected cells as “autophagosome-like” and “microvesicle-like”, because while markers for these vesicles have been identified in PV-infected cells, it has not been unambiguously determined that these are either precisely autophagosomes or microvesicles.

**Fig. 9.**
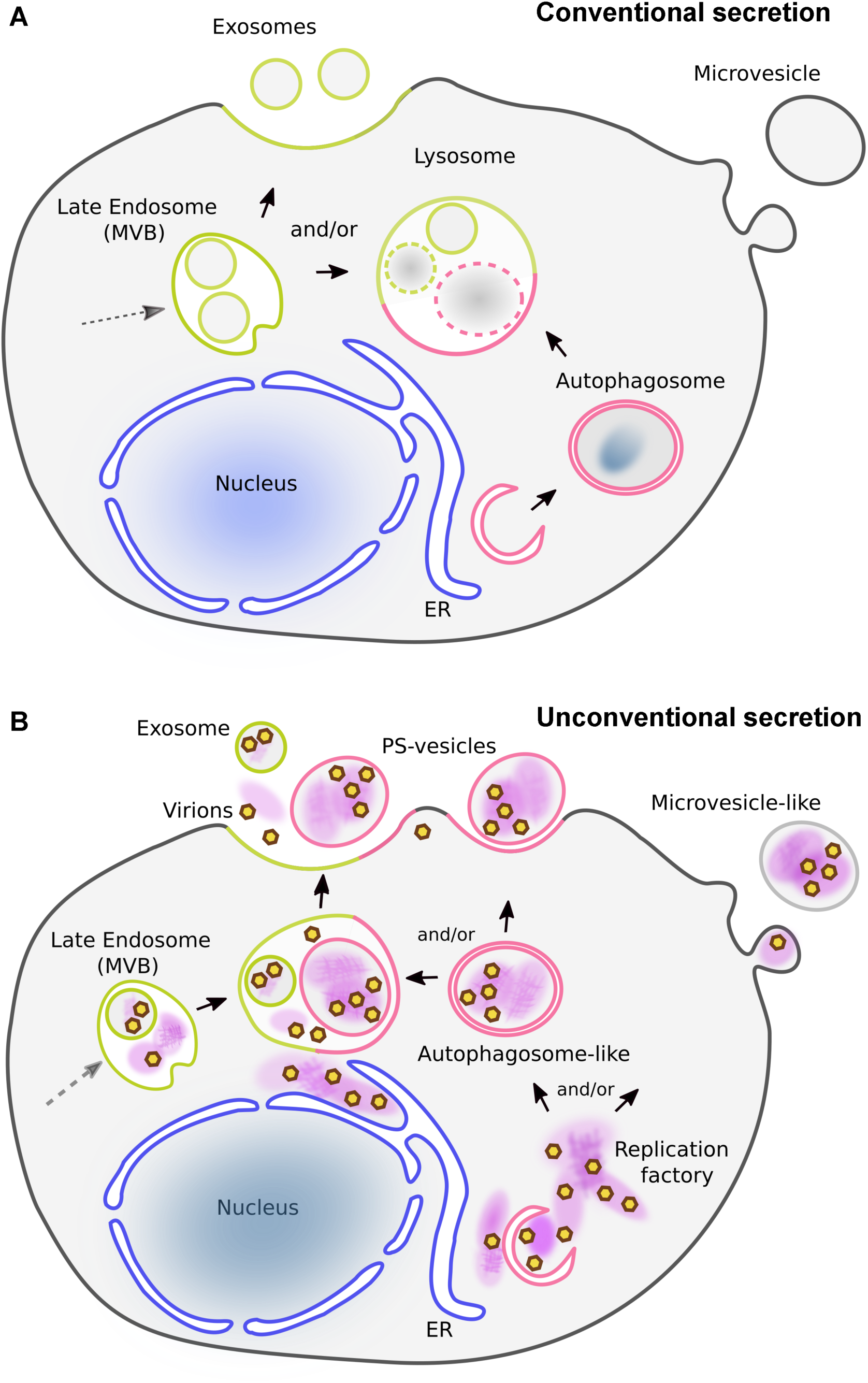
Models of cellular secretion from mock- and poliovirus-infected cells. (A) Endocytosis in uninfected cells provides a pathway for cell entry and distribution of internalized content to cell compartments. Cell sorting recycles selected components back to the plasma membrane while others become components of late endosomes, also called multivesicular bodies (MVBs). Components within the MVBs are then either exported as exosomes using (endosomal) ESCRT machinery, or fuse with lysosomes, where contents are degraded. Distinct from the endosomal pathway, autophagy is a mechanism to entrap and degrade specific cellular compartments and invading pathogens within double-membrane vesicles. Termed autophagosomes, these vesicles then fuse with lysosomes and the contents are degraded. In uninfected cells stimulated by, e.g., stress or starvation, lysosomes are the convergence point for the autophagic and endosomal pathways. (B) As PV infection progresses, membrane remodeling and lipid synthesis produce replication factories (purple patches) for vRNA synthesis. Prior to cell lysis, infectious exosomes and infectious microvesicles are formed within PV-infected cells. Infectious exosomes are present within late endosomes (also called multivesicular bodies, or MVBs; green). Exosomes are secreted either directly from MVBs, or after their fusion with autophagosome-like double membrane vesicles (pink). Infectious microvesicle-like structures can either bud directly from the cytoplasm, as in uninfected cells (see Figure S1), or they can be secreted after fusion of autophagosome-like vesicles with MVBs. Packaged inside exported vesicles are virions (brown/yellow hexagons), viral proteins, host proteins, host RNA, and both template and genomic viral RNA (together depicted as purple patches). These virion-containing extracellular vesicles get internalized into a neighboring cell. A rapid initiation of viral replication is achieved by this transport and internalization of all components needed for replication: virions, viral proteins, cellular proteins, ribosomes, viral RNA, and cellular RNA.

### Vesicle components 1: viral RNAs and proteins

As has been shown previously (Bird et al., 2014; Chen et al., 2015; Jackson et al., 2005), we provide direct evidence that secreted infectious microvesicles transport virions from cell to cell (Figs. 4A, 6-8). We further show that exosomes are also infectious and involved in PV nonlytic cell-to-cell transmission (Fig. 4B). It has been proposed that multiple capsid-enclosed viral genomes that are transferred by vesicles *en bloc* into a single cell are the sole contributor to replication kinetics in the new host (Chen et al., 2015). Indeed, this is the case for exosomes secreted from HAV-infected cells (McKnight et al., 2017). However, the delivery of (+) and (-) vRNA and viral replication proteins by extracellular vesicles into cells suggests that viral replication of PV may be facilitated by the delivery of (+) RNA, (-) RNA, and the viral proteins necessary for replication (e.g. the virus polymerase 3Dpol) together into new host cells. However, because there are data showing a dependence of viral replication on *cis*-translation of a central region of the PV genome (Novak and Kirkegaard, 1993), the question arises of roles for vesicle-transported non-structural proteins in *trans*. Perhaps the proximity of the proteins and RNA upon delivery overcomes the requirement of new translation, by eliminating the problem of diffusion-limited localization, such that the proteins are positioned as they would be, were they newly synthesized. We also note that inclusion of ribosomes in exosomes from PV-infected cells could facilitate passage of transported vRNA through these ribosomes. This could result in production of a specific quaternary structure of vRNA necessary for replication (Novak and Kirkegaard, 1993), providing a possible mechanism for subverting the *cis*-translation requirement for replication.

Virus-induced membrane remodeling occurs at approximately 3 hpi in PV-infected cells (e.g. (Belov et al., 2012; Rossignol et al., 2015)). With the transfer of membrane-remodeling proteins 2C, 2BC and 3AB that are ready for direct use, intact infectious extracellular vesicles could more quickly establish these membranous complexes for nascent viral replication. Support for this is the previous discovery that free virions from freeze-thawed microvesicles and intact infectious microvesicles induce similar (+) vRNA production when cells are pretreated with the RNA synthesis inhibitor guanidine hydrochloride. (Chen et al., 2015). We interpret this to mean that the increase in vRNA from vesicle-based infection is a result of newly synthesized (+) vRNA and not due simply to delivery of a larger quantity of vRNA.

### Vesicle components, 2: host proteins and host RNA

For infection propagation, viruses alter cell processes, facilitating the viral life cycle. This includes utilization of host cell proteins as well as ribosomes for translatation of viral proteins. Examples include 1) virus-induced reorganization of the actin cytoskeleton for viral entry (Baird et al., 2012; Coyne et al., 2007) and for transport of viral components within the cell (Vaughan et al., 2009), and 2) utilization of host proteins during PV infection to shut down cap-dependent protein translation, so that protein production continues almost exclusively for (cap-independent) viral proteins (Schmid and Wimmer, 1994). With less than 10% of the proteins identified by mass spectrometry in secreted infectious vesicles being virus proteins, in our model, transported host cell proteins and RNA have roles in advancing PV infection after cell-to-cell spread by vesicles.

Extracellular vesicles of different types have distinct roles in uninfected cells (Kanada et al., 2015), with studies suggesting selective packaging of RNAs and proteins through separate cellular pathways (Janas et al., 2015; Villarroya-Beltri et al., 2013). Therefore, differences in composition between microvesicles and exosomes from PV-infected cells suggest the possibility that some infection-related functions are specific for each vesicle type. For example, our mass spectrometry data identified a significant enrichment of proteins from the glycolytic pathway in infectious microvesicles (p<10^-9^; Table 1), including a strong and unique presence of pyruvate kinase PKM, the critical glycolysis rate-limiting enzyme. In addition, six other enzymes important for glycolysis were identified, and were absent from infectious exosomes. It is known that the presence of glucose during PV infection causes an average 170-fold increase in viral output (Eagle and Habel, 1956). Thus, a vesicle-infected host cell that has received PKM may be better prepared for viral replication than cells infected by free virions or infectious exosomes, due to the possibility of producing the large increase in glucose (or fructose) and glutamine known to be required for maximal PV replication (Darnell and Eagle, 1958; Eagle and Habel, 1956).

Compared to infectious microvesicles and vesicles from mock-infected cells, a more predominant presence of host cell ribosome proteins and rRNAs was observed in PV infectious exosomes. Similar to exosomes from uninfected cells (Tkach and Théry, 2016), we show that exosomes carry a repertoire of microRNAs and a large variety of small non-coding RNAs. However, distinct from uninfected cells, our RNA-Seq data show that rRNAs, and C/D Box snoRNAs that modify and stabilize rRNAs, were over-represented in infectious exosomes (Table 2), consistent with the significant fraction of ribosomal protein subunits identified by mass spectrometry (Fig. 2). Thus, much in the same way that enveloped viruses, such as arenavirus, contain cellular ribosomes (Rowe et al., 1970), these data suggest that transported ribosomal components from the host, with their proximity to vRNA, could help ensure a rapid and smooth production of viral proteins after vesicle entry in newly infected hosts, thereby facilitating infection.

Changes in immune responses and susceptibility to infection have also been observed in PV-infected cells (Yoshikawa et al., 2006). The chaperone protein HSP90, which is a facilitating factor in viral replication of hepatitis A virus (McKnight et al., 2017), hepatitis C virus (HCV) (Okamoto et al., 2006) and influenza A (Chase et al., 2008), was identified only in infectious vesicles, in both microvesicles and exosomes. Recent studies on Enterovirus 71, a member of the picornaviridae family, showed that virions transported in exosomes were able to replicate in new target cells, while the inflammatory type I interferon response was suppressed due to miRNAs co-delivered by these exosomes (Fu et al., 2017). Therefore, proteins involved in immune response seem to be excluded from, and negative regulators included in, infectious exosomes either because virus infection changed the expression levels of these cellular components, or because they are actively targeted to or excluded from secreted vesicles, respectively. Infectious vesicles are thereby proposed to be specifically promoting an immune-depressing and infection-promoting environment in the target cell.

### Structures of Infectious Vesicles

Structure and function are often linked. Therefore, it is not surprising that after we identified the large number of components in infectious microvesicles and exosomes using biochemical methods, we were also able to visualize significant structural complexity in infectious vesicles that clearly contained internal components in addition to virions. Extracellular vesicles from PV-infected cells included dense, mat-like structures (Figs. 6-8), often seen adjacent to membranes, and/or in close proximity to virions. This juxtaposition suggests a possible involvement of protein-lipid interactions that results in the observed spatial arrangement of the cargo. In our model, these mats are composed of the non-capsid proteins identified by mass spectrometry (Fig. 1B and Table S1) and western blots (Fig. 1C); vRNA identified by RT-qPCR (Fig. 4B); and cellular and viral RNA identified by RNA-Seq (Fig. 3, Tables 2 and S2). We note that unencapsidated (+) and (-) vRNA could be present as dsRNA, which would provide additional protection from the host immune system. Class III infectious vesicles, which are defined by the presence of internal membrane structures within the infectious microvesicle lumen, comprised only a minor population. It is currently unclear if Class III infectious vesicles are produced accidentally when a smaller vesicle happens to become entrapped, or if they originate from a cellular mechanism similar to the formation of mutivesicular bodies, where the inclusion of additional components advances virus replication.

### Cellular Origins of Infectious Vesicles

Cellular transport in uninfected cells, diagrammed in Figure 9A, is altered in response to intracellular stress, such as an invading pathogen (Baixauli et al., 2014). Autophagosomes can fuse with early or late endosomes/MVBs to form amphisomes (Eskelinen, 2005), and unconventional secretion can occur via secretory autophagy (Ponpuak et al., 2015) or lysosomal exocytosis (Settembre et al., 2013). Therefore, the endosomal, exosomal, lysosomal, and autophagic pathways are overlapping under some cellular conditions. Thus, it is not surprising that we identified both endocytic and exosomal marker proteins in infectious vesicles. We did not, however, identify autophagosome-associated lipidated LC3 (LC3-II). LC3-II was previously shown within cells, in autophagosome-like double membrane vesicles that contain virions; these autophagosomes are thought to be the precursor of secreted infectious vesicles (Bird et al., 2014; Chen et al., 2015; Feng et al., 2013; Jackson et al., 2005; Robinson et al., 2014). This omission in our results, and previously in exosomes from HAV-infected cells (McKnight et al., 2017), is likely due to technical limitations in detecting lipidated proteins by LC-MS (Karpievitch et al., 2010).

In our model (Fig. 9B), the endosomal and autophagosomal pathways both feed into and provide exits from viral replication factories that then utilize unconventional secretion for non-lytic spread of infectious vesicles, thereby accelerating viral replication in new cells.

### Summary

In summary, our results demonstrate that infectious vesicles derived from non-enveloped picornavirus-infected cells represent a new viral transmission model, in which multi-component transport of viral and host RNA and proteins facilitate cell-to-cell spread of viral infection by the non-enveloped virus PV.

## Materials & Methods

### Cell Culture and PV Infection

See below for K562 cell methods (RNA-Seq Sample Preparation and Data Analysis). HeLa cells (ATCC, Manassas, VA) were infected with Mahoney PV (gift from Dr. Karla Kirkegaard, Stanford University) using protocols from Burrill, Strings, and Andino (Burrill et al., 2013b). Briefly, HeLa cells were cultured in low-glucose DMEM medium supplemented with 5% fetal bovine serum (FBS) and 1% penicillin-streptomycin-glutamine (supplDMEM; Atlanta Biologicals, Cat.S11150; ThermoFisher, Cat.10378016, respectively) and grown at 37 °C (5% CO_2_) to 60 to 80 % confluence. After one wash with PBS+ (PBS supplemented with 0.01 mg/ml MgCl_2_ and 0.01 mg/ml CaCl_2_), cells were infected either with PBS+ or PBS+ and PV stock at a multiplicity of infection (MOI) of 30 virions per cell, titrated by classic plaque assay (see below). After 30 min at 37 °C, cells were washed in PBS + and grown in supplDMEM. For the infectious microvesicle-induced infection assay shown in Figure 3, infection was first synchronized by addition of infectious microvesicles on ice for 30 min to promote adherence, then washed with PBS+, as in (Kronenberger et al., 1992; Tsang et al., 2001) prior to growth at 37 °C in supplDMEM. Because PV replication and release of infectious extracellular vesicles decreased when cells were cultured in non-bovine serum media ((Burrill et al., 2013b) and data not shown), we first infected HeLa cells in FBS-containing media for 4 h, during which the RNA replication rate reaches its maximum. We then minimized contamination of the extracellular vesicles by components from bovine serum (FBS) by washing (3x) and then growing the cells in non-FBS media before collecting vesicles at 8 hpi, when cells and/or the supernatant were collected for subsequent analyses.

To test for cell viability at the time of vesicle collection (Fig. 1B), HeLa cells were infected with PV at an MOI of 30 for 8h, then stained in situ with a 0.2% Trypan Blue solution in PBS. Persistently PV-infected K562 cells were collected and stained with a 1:1 dilution of the cell suspension in 0.4% Trypan Blue-PBS. In each case, samples were double-blinded and two randomly selected fields of approximately 200 cells from each of three replicate wells were counted. Percentage viability was calculated as number of blue-staining cells divided by the total cells counted x 100.

### Collection and Purification of Microvesicles and Phosphatidylserine-containing Microvesicles

The non-FBS media supernatant of control (mock-infected) or PV-infected HeLa cells at 8 hpi was collected as described above, and infectious microvesicles were purified as in (Chen et al., 2015). Briefly, all at 4 °C, the supernatants were spun down at 150 x g for 15 min and then 1,010 x g for 20 min to remove cell debris and apoptosis bodies, respectively. The supernatant was then centrifuged at 10,000 x g, 60 min, to collect the extracellular vesicles with a size range of 100 nm to 1 µm in diameter, as in (Lane et al., 2017). Both the supernatant and pellet were saved. For all experiments except LC/MS, the annexin-V microbead kit (Miltenyi Biotec, Cat. 130-090-201) was used for further enrichment of the pelleted PS-containing vesicles, per manufacturer’s instructions. The eluted PS-enriched vesicles were spun down and resuspended in 1X PBS for subsequent assays.

### Collection and Purification of Exosomes and CD9-Positive Exosomes

To isolate CD9-positive exosomes, supernatants from the previously described 10,000 x g centrifugation were spun down at 100,000 x g for 1 h, the pellet was washed with 1X PBS, and spun again at 100,000 x g. The final pellet was resuspended in 1X PBS, and exosomes were collected using ExoQuick-TC (System Biosciences Inc.) as per manufacturer’s instructions. For all experiments except RNA-Seq, the collected exosomes were further purified with anti-CD9 antibody-coated magnetic beads (CD9 Exo-Flow Exosome Purification Kit, System Biosciences, Cat. EXOTC10A-1) as in (Rim and Kim, 2016; Smith and Daniel, 2016).

### Sample Preparation of Broken Extracellular Vesicles

Extracellular vesicles were freeze-thawed (3X) to break vesicle membranes. For experiments in which RNA was to be removed, freeze-thawed vesicles were incubated with 1% sodium deoxycholate (300 µg/ml) on ice for 45 minutes as in (Egger et al., 1996). The sample was then incubated with RNaseA (600 µg/ml, Sigma Cat. 10109142001) for 1 h at 37 °C, to remove exposed RNA, as in (Nugent et al., 1999); at the 150 mM salt concentration in these experiments, we expect both ssRNA and dsRNA to be removed (Ausubel et al., 2003).

### Immunoblotting

Purified extracellular vesicles were washed with 1X PBS, disrupted with chilled RIPA buffer (50 mM Tris/HCl pH 7.4, 75 mM NaCl, 1 mM EDTA, 1% NP-40, 0.25% sodium deoxycholate, 0.1% sodium dodecyl-sulfate) containing a protease inhibitor cocktail diluted 1:200 (Sigma, # P1860), and then incubated shaking for 30 min at 4 °C. Protein concentrations were determined by bicinchoninic assay (BCA, Thermo Fisher, # 23225). Four antibodies were used for immunoblotting: 1) anti-PV (type 1-3) goat IgG (Abcam, # ab22450); 2) anti-3Dpol rabbit IgG prepared by Cocalico Biologicals from purified 3Dpol protein, 1 mg/ml, diluted 1:1000; 3) anti-2C (CPSQEHQEILFN) rabbit IgG (Cocalico Biologicals, 1 mg/ml, diluted 1:1000); 4) anti-3A (KDLKIDIKTSPPPEC; (Richards et al., 2014)) rabbit IgG (Biomatik, 0.6 mg/ml, diluted 1:500). Protein derived from extracellular vesicles (20 µg per lane) was subjected to SDS-PAGE or Tricine-SDS gel electrophoresis and western blotting, as described previously (Schägger, 2006).

### RT-qPCR

RNA from whole cells or extracellular vesicles was extracted using the RNeasy Plus Mini Kit (Qiagen, # 74134), as in (Eldh et al., 2012). The full-length PV genome and its negative-sense template were amplified, in duplicate, through two-step RT-qPCR, as previously described (Burrill et al., 2013b). Briefly, cDNA was synthesized using the SuperScript III RT (Life Technology, # 18080-093) system; RT (Schulte et al., 2015), with Tag primer to increase binding specificity, full-length production, and efficiency. The qPCR was performed using a master mix (Fast SYBR Green master mix system; Life Technology, # 4385610). Detection of RNA was performed in a two-step RT-qPCR using the High Capacity cDNA Reverse Transcription Kit (Applied Biosystems, # 4374966) and the SYBR green master mix. In order to demonstrate the specificity of amplification, we conducted a series of controls including negative reverse transcription control and non-template controls, and melt curve analyses. As described in (Schmittgen and Livak, 2008), results were normalized to an endogenous control GAPDH of whole cells presented as ΔC_t_ (where ΔC_t_ = (C_t_ of endogenous control gene (GAPDH)) – (Ct of gene of interest)) (Livak & Schmittgen 2001; Schmittgen & Livak 2008). The mean of two technical replicates per cDNA sample was used to obtain raw C_t_ and ΔC_t_ for quantitative gene expression. The statistical analysis was obtained from six biological replicates. A non-paired one tailed Student’s t-test was used for the independent sample comparison, e.g. PV-infected (ImV) versus mock-infected (MmV) groups. A paired one tailed Student’s t-test was performed for comparison between the groups of untreated ImV/MmV and FTDR treated ImV/MmV. The Bonferroni method was used to calculate the *p*-value (cutoff = 0.05).

### Plaque Assay

Virus titrations were performed as described previously (Burrill et al., 2013b; Robinson et al., 2014). Briefly, HeLa cells were seeded to a concentration of 2.0 x 10^6^/60 mm diameter tissue culture dishes and grown overnight. Serial dilutions of extracellular vesicles were then added, in duplicate, onto the cells for 30 min at 37 °C, 5% CO_2_, then overlayed with 1% agar in supplDMEM, and incubated 48 h. After fixation with 2% formaldehyde, 0.25% crystal violet stain was added for 10 min and rinsed in water. Plaques were counted manually.

### Plunge Freezing for Cryo-electron Microscopy

Vesicle samples were freshly prepared for each experiment. C-Flat 4/2 holey carbon grids (Protochips, # CF-4/2-2C-50) were stabilized with an extra layer of carbon on the carbon surface. A 3 µl sample of extracellular vesicles was applied onto freshly glow-discharged grids. Samples for electron tomography included 0.5 µl of 5-nm fiducial gold (Ted Pella Inc. Cat. 82150-5) applied to the sample drop, and incubated for 1.5 minutes at 10 °C, 100% humidity. The grid was then blotted and plunge-frozen in a Vitrobot Mark III (FEI, Oregon), and stored in liquid nitrogen.

### Cryo-electron Tomography

Initial cryo-electron microscopy was performed on a Philips CM12 EM at 100 kV (TVIPS CCD camera, pixel size of 6.8 Å, electron dose ≤ 30 e^-^/Å^2^). To collect data for 3-D reconstructions, the majority of the single axis tilt series (typical tilt range −56° to + 56°, angle increment 2°, total electron dose ≤ 100 e^-^/Å^2^, defocus of −4 to −5 µm, were acquired on an FEI Tecnai F20 (TF20) EM at 160 kV (TVIPS CMOS camera; 2x binned pixel size of 8.3 Å) using SerialEM software (Mastronarde, 2005). Alternatively, some single axis tilt series were collected on a JEM JEOL 2200 FS equipped with an in-column Omega energy filter, at 200 kV on a Direct Electron Detector (DE20) using SerialEM software (Mastronarde, 2005), resulting in a pixel size of 4.01 Å. The typical tilts were recorded from −60° to + 60°, with an angle increment of 2°, total electron dose ≤ 100 e^-^/Å^2^, defocus of −4 µm. The tilt images collected on the DE20 were motion corrected using scripts provided by the manufacturer (Direct Electron, LP). Reconstructions of tomograms and 3-D surface model rendering were performed using the weighted back-projection method from the IMOD software package (Kremer et al., 1996). Volume averaging of virions inside vesicles was performed using PEET software (Cope et al., 2011; Nicastro et al., 2006) and the 3-D surface model of virions was rendered through averaged and low pass filtered sub-volumes from smoothed tomograms. Nonlinear anisotropic diffusion (NAD) filtering was used to generate 3-D surface models of inner mat-like structures in the infectious microvesicles, to provide better density continuity and reduced noise.

### Subtomogram averaging

Subvolumes of virus-like particles inside ImVs were manually selected (n = 1,022) from 32 smoothed tomograms (data recorded on an FEI TF20 EM) using the IMOD software package to determine the initial orientation of each particle. Alignment and averaging were performed using the PEET software (which is part of the IMOD package). Classification analyses using principal component analysis (PCA) were performed on a set of aligned sub-volumes (Cope et al., 2011) to group structurally homogeneous particles using clusterPca in PEET. The majority of virions (820 out of 1,022) were identified in class 1, 160 virions in class 2, 32 virions in class 3, and 10 virions in class 4. Class 1 virions with high cross-correlation coefficients (118 particles) were averaged, including imposing icosohedral symmetry; the process was then iterated using the average as the new reference. Based on the resolution estimation using the Fourier Shell Correlation (FSC at 0.5 cutoff), the final subtomogram average was low-pass filtered to a resolution of 6.9 nm and each virion in the tomogram is displayed using isosurface-rendering of the symmetrized averaged volume.

### Liquid Chromatography-Mass Spectrometry

Infectious microvesicles (without annexin-V purification) and CD9-positive exosomes from mock- and PV-infected cells at 8 hpi were collected as described above. Collected vesicles were sonicated in an ice-water bath for 4 min, then incubated with 2,2,2,-Trifluoroethanol (TFE) (Sigma, # T63002) (1:1, V:V), for 2 h at 60 °C with shaking. Ammonium bicarbonate (J.T. Baker) and dithiothreitol (DTT) (Sigma) were added at room temperature (RT), to final concentrations of 50 mM and 5 mM, respectively, and incubated at 60 °C for 30 min. Iodoacetamide (Bio-Rad) was added to a final concentration of 10 mM, incubated 30 min at RT in the dark and then quenched with DTT. After dilution to 5% TFE with a 3:1 mixture of water:50 mM ammonium bicarbonate, sequencing-grade trypsin (Promega Corp. Madison, WI) was added at 1:30 enzyme to substrate (V:V), incubated overnight at 37 °C, quenched with neat formic acid and cleaned using C18 PepClean spin columns (Pierce Biotechnology, Rockford, IL). Peptides from three technical replicates were analyzed by liquid chromatography–tandem mass spectrometry (LC-MS/MS) using a Q Exactive Hybrid Quadrupole-Orbitrap mass spectrometer (Thermo-Fisher) in positive mode, equipped with a nanoAcquity UPLC system, (nanoAcquity NPLC Symmetry C18 trap column and ACQUITY UPLC Peptide BEH C18 analytical column; Waters) and a Triversa Nanomate source (Advion, Ithaca, NY). Using mobile phase A (1% acetonitrile/0.1% formic acid) and mobile phase B (99% acetonitrile/0.1% formic acid), peptides were trapped for 4 min at 4 µL/min in A, and then separated using the gradient: 0–8 min: 2% B, 8–96 min: 2–40% B, 96–102 min: 40% B, 102–105 min: 40–70% B, 105–113 min: 70% B, 113– 114 min: 70–2% B, and 114–120 min: 2% B. MS spectra were acquired at 70,000 resolution at *m/z* 400, scan range *m/z* 370–1800, 1 microscan per spectrum, AGC target of 1e6, maximum injection time 100 ms. The 10 most abundant precursor ions per scan were fragmented at 17,500 resolution at *m/z* 400, AGC target 1e6, maximum injection time 100 ms, isolation window 10.0 *m/z*, isolation offset 0.4 *m/z*, normalized collision energy (NCE) 26, exclusion of unassigned charge states and charge states 1, and dynamic exclusion 8 s. Profile data were recorded for both MS and MS/MS scans.

### Proteomics Data Analysis

LC-MS/MS data were processed using Peaks Studio 8.5 (Bioinformatics Solutions, Waterloo, ON) (Zhang et al., 2012), using a database containing the PV proteome, either as one combined polypeptide or as individual polio proteins and stable intermediates, concatenated with human and bovine Uniprot proteomes (Magrane and Consortium, 2011). Technical replicates were searched together. We specified precursor ion (MS1) error tolerance of 10 ppm, and a fragment ion (MS/MS) error tolerance of 0.02 Da and a target-decoy false discovery threshold of 0.1%. Proteins identified as exclusively bovine were discarded, and we required protein identifications to contain at least two unique peptides when searching against the PV proteome as one polypeptide, and at least one unique peptide for searches against individual polio proteins and stable intermediates. Protein abundances were calculated by aggregating the MS1 peak areas for all the peptides identified for a specific protein, and normalized to an endogenous control GAPDH of whole cells.

To identify cellular pathways that were significantly associated with the identified protein matches, p-values of overlap were determined by IPA software (Qiagen) using Fisher’s exact test. Pathways shown to be enriched (protein matches were over-represented in our datasets as compared to a random dataset of the cellular proteome) were included when –log(p-value) >3.

### RNA-Seq Sample Preparation and Data Analysis

RNA-Seq: K562 cells (ATCC CRL-3343) were infected with PV and the infection allowed to reach a persistent steady-state over several days. 10^6^ infected or control cells were washed thoroughly with PBS, then the media was replaced. After 24 h, medium was collected by two rounds of spin-down of cells (1000 x g, 5 min). Exo-Quick precipitation was carried out according to manufacturer’s instructions (System Biosciences Inc.), with 3 ml ExoQuick added to approximately 15 ml tissue culture supernatant for each condition. Precipitation was carried out at 4 °C overnight. Precipitates were centrifuged at 1500 × g for 30 min. Phenol-free lysis buffer (SeraMir, System Biosciences) was added to pellets, vortexed, and samples were sent to the Institute for Genome Sciences (Univ. Maryland) for RNA column purification.

Next Gen Sequencing libraries were prepared by System Biosciences and sequenced on an Illumina Hi-Seq instrument with 50 bp single-end reads at a minimum depth of approximately 15 million reads per sample. In-depth data analysis on raw data against the reference human genome was provided by Maverix Biomics. Data analysis against viral sequences and assistance with the interpretation of RNA-Seq data was provided by the Institute for Genome Sciences.

### Schematic Diagrams

Schematic diagrams were drawn using Inkscape open source software (inkscape.org).

### Accession Numbers

Cryo-ET data have been deposited in the wwPDB, with EMDB accession codes: EMD-7877, EMD-7878, EMD-7879, EMD-7880, EMD-7881.

## Acknowledgments

We gratefully acknowledge the assistance of Dr. Wah Chiu and use of the CryoEM Data Collection Facility Consortium at NCMI.

## Supporting Information Legends

**Supplemental Movie. Cryotomogram of a microvesicle from poliovirus-infected cells**

Movie of an electron cryotomogram of a Class III microvesicle isolated from poliovirus-infected cells. Labeled, and later shown in isosurface rendered models in the tomogram, are virions, vesicle membranes, mat-like structures, and the inner vesicular structure. Additional labels indicate representative annexin-V beads that were used to isolate the sample, and fiducial gold beads that were added to aid alignment of tilt series data. Scale bar, 100 nm.

**Table S1. Proteomic profiles of microvesicles and exosomes from PV- and mock-infected cells at 8 hpi.**

The full proteomic profile of microvesicles from PV-infected cells (parts 1-4) and mock-infected cells by LC/MS shows 168 host protein matches in infectious microvesicles and 14 host protein matches in microvesicles from mock-infected cells (B). The full proteomic profile of CD9-positive infectious exosomes (C) and mock exosomes (D) shows 162 host proteins identified in infectious exosomes and 58 host proteins identified in exosomes from mock-infected cells. All proteomic analyses were conducted at a specified precursor ion (MS1) error tolerance of 10 ppm, a fragment ion (MS/MS) error tolerance of 0.02 Da. and a target-decoy false discovery threshold of 0.1%.

**Table S2. RNA-Seq profiles of exosomes from PV- and mock-infected K562 cells.**

In total, there are 14 types of RNAs identified in both infectious and mock exosomes. The number under each sample type represents the number of identifiable short reads in the corresponding sample. The table contains all RNAs with at least 1000 short RNA-Seq reads.

