## Supplemental Tables 1 and 2 for "Complexity and ultrastructure of infectious extracellular vesicles from cells infected by non-enveloped virus"

### Table S1A

| Human from infected extracellular microvesicles (ImVs) | | | | | | | | | | |
|---|---|---|---|---|---|---|---|---|---|---|
| | Accession # | Protein Abbreviation | -10lgP | Area from one run (IP3_030917_ 3reps) | Description | Present in both repeats? | Present in both ImV repeats AND in Iexo? | in ImV and Iexo? | In ImV and MmV? | In ImV and Mexo? |
| | P60709 | ACTB_HUMAN | 378.78 | 975000000 | Actin cytoplasmic 1 OS=Homo sapiens GN=ACTB PE=1 SV=1 | TRUE | TRUE | TRUE | TRUE | TRUE |
| | P63261 | ACTG_HUMAN | 378.78 | 975000000 | Actin cytoplasmic 2 OS=Homo sapiens GN=ACTG1 PE=1 SV=1 | TRUE | TRUE | TRUE | TRUE | TRUE |
| | P35908 | K22E_HUMAN | 330.12 | 517000000 | Keratin type II cytoskeletal 2 epidermal OS=Homo sapiens GN=KRT2 PE=1 SV=2 | TRUE | TRUE | TRUE | TRUE | TRUE |
| | P04264 | K2C1_HUMAN | 307.63 | 488000000 | Keratin type II cytoskeletal 1 OS=Homo sapiens GN=KRT1 PE=1 SV=6 | TRUE | TRUE | TRUE | TRUE | TRUE |
| | P13645 | K1C10_HUMAN | 266.4 | 416000000 | Keratin type I cytoskeletal 10 OS=Homo sapiens GN=KRT10 PE=1 SV=6 | TRUE | TRUE | TRUE | TRUE | TRUE |
| | P80723 | BASP1_HUMAN | 370.94 | 288000000 | Brain acid soluble protein 1 OS=Homo sapiens GN=BASP1 PE=1 SV=2 | TRUE | TRUE | TRUE | FALSE | FALSE |
| | P08174 | DAF_HUMAN | 190.48 | 216000000 | Complement decay-accelerating factor OS=Homo sapiens GN=CD55 PE=1 SV=4 | TRUE | TRUE | TRUE | FALSE | TRUE |
| | O75340 | PDCD6_HUMAN | 213.31 | 213000000 | Programmed cell death protein 6 OS=Homo sapiens GN=PDCD6 PE=1 SV=1 | TRUE | TRUE | TRUE | FALSE | FALSE |
| | P35527 | K1C9_HUMAN | 206.24 | 178000000 | Keratin type I cytoskeletal 9 OS=Homo sapiens GN=KRT9 PE=1 SV=3 | TRUE | TRUE | TRUE | TRUE | TRUE |
| | P50995 | ANX11_HUMAN | 269.92 | 148000000 | Annexin A11 OS=Homo sapiens GN=ANXA11 PE=1 SV=1 | TRUE | TRUE | TRUE | FALSE | FALSE |
| | P08195 | 4F2_HUMAN | 214.05 | 111000000 | 4F2 cell-surface antigen heavy chain OS=Homo sapiens GN=SLC3A2 PE=1 SV=3 | TRUE | TRUE | TRUE | TRUE | TRUE |
| | P04406 | G3P_HUMAN | 227.96 | 92700000 | Glyceraldehyde-3-phosphate dehydrogenase OS=Homo sapiens GN=GAPDH PE=1 SV=3 | TRUE | TRUE | TRUE | FALSE | TRUE |
| | P13987 | CD59_HUMAN | 190.45 | 61500000 | CD59 glycoprotein OS=Homo sapiens GN=CD59 PE=1 SV=1 | TRUE | TRUE | TRUE | FALSE | TRUE |
| | P0CG48 | UBC_HUMAN | 144.04 | 54000000 | Polyubiquitin-C OS=Homo sapiens GN=UBC PE=1 SV=3 | TRUE | FALSE | FALSE | TRUE | TRUE |
| | P62979 | RS27A_HUMAN | 144.04 | 54000000 | Ubiquitin-40S ribosomal protein S27a OS=Homo sapiens GN=RPS27A PE=1 SV=2 | TRUE | TRUE | TRUE | TRUE | TRUE |
| | P62987 | RL40_HUMAN | 144.04 | 54000000 | Ubiquitin-60S ribosomal protein L40 OS=Homo sapiens GN=UBA52 PE=1 SV=2 | TRUE | FALSE | FALSE | TRUE | TRUE |
| | P31949 | S10AB_HUMAN | 140.88 | 52900000 | Protein S100-A11 OS=Homo sapiens GN=S100A11 PE=1 SV=2 | TRUE | TRUE | TRUE | FALSE | FALSE |
| | P30626 | SORCN_HUMAN | 157.35 | 42900000 | Sorcin OS=Homo sapiens GN=SRI PE=1 SV=1 | TRUE | TRUE | TRUE | FALSE | FALSE |
| | P62937 | PPIA_HUMAN | 173.59 | 40700000 | Peptidyl-prolyl cis-trans isomerase A OS=Homo sapiens GN=PPIA PE=1 SV=2 | TRUE | TRUE | TRUE | FALSE | FALSE |
| | P04083 | ANXA1_HUMAN | 218.34 | 31700000 | Annexin A1 OS=Homo sapiens GN=ANXA1 PE=1 SV=2 | TRUE | TRUE | TRUE | FALSE | FALSE |
| | P02533 | K1C14_HUMAN | 237.36 | 30700000 | Keratin type I cytoskeletal 14 OS=Homo sapiens GN=KRT14 PE=1 SV=4 | TRUE | FALSE | FALSE | FALSE | TRUE |
| | P60174 | TPIS_HUMAN | 164.69 | 27000000 | Triosephosphate isomerase OS=Homo sapiens GN=TPI1 PE=1 SV=3 | TRUE | FALSE | FALSE | FALSE | FALSE |
| | Q9UBV8 | PEF1_HUMAN | 118.08 | 25400000 | Peflin OS=Homo sapiens GN=PEF1 PE=1 SV=1 | TRUE | FALSE | FALSE | FALSE | FALSE |
| | P05023 | AT1A1_HUMAN | 204.08 | 25300000 | Sodium/potassium-transporting ATPase subunit alpha-1 OS=Homo sapiens GN=ATP1A1 PE=1 SV=1 | TRUE | FALSE | FALSE | FALSE | FALSE |
| | P08758 | ANXA5_HUMAN | 142.1 | 24600000 | Annexin A5 OS=Homo sapiens GN=ANXA5 PE=1 SV=2 | TRUE | TRUE | FALSE | FALSE | FALSE |
| | P35613 | BASI_HUMAN | 253.39 | 24400000 | Basigin OS=Homo sapiens GN=BSG PE=1 SV=2 | TRUE | TRUE | TRUE | TRUE | TRUE |
| | P06733 | ENOA_HUMAN | 142.93 | 23900000 | Alpha-enolase OS=Homo sapiens GN=ENO1 PE=1 SV=2 | TRUE | FALSE | FALSE | FALSE | FALSE |
| | P26038 | MOES_HUMAN | 178.96 | 23500000 | Moesin OS=Homo sapiens GN=MSN PE=1 SV=3 | TRUE | TRUE | TRUE | FALSE | FALSE |
| | P09525 | ANXA4_HUMAN | 181.51 | 19800000 | Annexin A4 OS=Homo sapiens GN=ANXA4 PE=1 SV=4 | TRUE | FALSE | FALSE | FALSE | FALSE |
| | P68104 | EF1A1_HUMAN | 83.73 | 19500000 | Elongation factor 1-alpha 1 OS=Homo sapiens GN=EEF1A1 PE=1 SV=1 | TRUE | TRUE | TRUE | FALSE | FALSE |
| | Q05639 | EF1A2_HUMAN | 83.73 | 19500000 | Elongation factor 1-alpha 2 OS=Homo sapiens GN=EEF1A2 PE=1 SV=1 | TRUE | FALSE | FALSE | FALSE | FALSE |
| | Q5VTE0 | EF1A3_HUMAN | 83.73 | 19500000 | Putative elongation factor 1-alpha-like 3 OS=Homo sapiens GN=EEF1A1P5 PE=5 SV=1 | TRUE | TRUE | TRUE | FALSE | FALSE |
| | P23528 | COF1_HUMAN | 204.53 | 19300000 | Cofilin-1 OS=Homo sapiens GN=CFL1 PE=1 SV=3 | TRUE | TRUE | TRUE | FALSE | FALSE |
| | P14618 | KPYM_HUMAN | 138.78 | 17500000 | Pyruvate kinase PKM OS=Homo sapiens GN=PKM PE=1 SV=4 | TRUE | TRUE | TRUE | FALSE | FALSE |
| | P08238 | HS90B_HUMAN | 150.15 | 15800000 | Heat shock protein HSP 90-beta OS=Homo sapiens GN=HSP90AB1 PE=1 SV=4 | TRUE | TRUE | TRUE | FALSE | FALSE |
| | P11142 | HSP7C_HUMAN | 225.22 | 14600000 | Heat shock cognate 71 kDa protein OS=Homo sapiens GN=HSPA8 PE=1 SV=1 | TRUE | TRUE | TRUE | FALSE | FALSE |
| | P20073 | ANXA7_HUMAN | 135.42 | 13500000 | Annexin A7 OS=Homo sapiens GN=ANXA7 PE=1 SV=3 | TRUE | FALSE | FALSE | FALSE | FALSE |

| Human from infected extracellular microvesicles (ImVs) | | | | | | | | | | |
|---|---|---|---|---|---|---|---|---|---|---|
| | Accession # | Protein Abbreviation | -10lgP | Area from one run (IP3_030917_3reps) | Description | Present in both repeats? | Present in both ImV repeats AND in Iexo? | in ImV and Iexo? | In ImV and MmV? | In ImV and Mexo? |
| | P63104 | 1433Z_HUMAN | 147.65 | 12000000 | 14-3-3 protein zeta/delta OS=Homo sapiens GN=YWHAZ PE=1 SV=1 | TRUE | TRUE | TRUE | FALSE | FALSE |
| | Q06830 | PRDX1_HUMAN | 90.78 | 11500000 | Peroxiredoxin-1 OS=Homo sapiens GN=PRDX1 PE=1 SV=1 | TRUE | TRUE | TRUE | FALSE | FALSE |
| | P04075 | ALDOA_HUMAN | 140.39 | 10700000 | Fructose-bisphosphate aldolase A OS=Homo sapiens GN=ALDOA PE=1 SV=2 | TRUE | TRUE | TRUE | FALSE | FALSE |
| | O75131 | CPNE3_HUMAN | 123.29 | 10200000 | Copine-3 OS=Homo sapiens GN=CPNE3 PE=1 SV=1 | TRUE | FALSE | FALSE | FALSE | FALSE |
| | P05556 | ITB1_HUMAN | 120.67 | 9540000 | Integrin beta-1 OS=Homo sapiens GN=ITGB1 PE=1 SV=2 | TRUE | FALSE | FALSE | FALSE | FALSE |
| | P02786 | TFR1_HUMAN | 140.98 | 7890000 | Transferrin receptor protein 1 OS=Homo sapiens GN=TFRC PE=1 SV=2 | TRUE | FALSE | FALSE | FALSE | FALSE |
| | P00338 | LDHA_HUMAN | 121.52 | 7580000 | L-lactate dehydrogenase A chain OS=Homo sapiens GN=LDHA PE=1 SV=2 | TRUE | FALSE | FALSE | FALSE | FALSE |
| | Q09666 | AHNK_HUMAN | 150.48 | 7160000 | Neuroblast differentiation-associated protein AHNAK OS=Homo sapiens GN=AHNAK PE=1 SV=2 | TRUE | TRUE | TRUE | FALSE | FALSE |
| | Q8WUM4 | PDC6I_HUMAN | 138.08 | 6960000 | Programmed cell death 6-interacting protein OS=Homo sapiens GN=PDCD6IP PE=1 SV=1 | TRUE | FALSE | FALSE | FALSE | FALSE |
| | Q6YHK3 | CD109_HUMAN | 74.04 | 6860000 | CD109 antigen OS=Homo sapiens GN=CD109 PE=1 SV=2 | TRUE | FALSE | FALSE | FALSE | FALSE |
| | Q16658 | FSCN1_HUMAN | 112.1 | 6460000 | Fascin OS=Homo sapiens GN=FSCN1 PE=1 SV=3 | TRUE | FALSE | FALSE | FALSE | FALSE |
| | P61586 | RHOA_HUMAN | 152.91 | 6180000 | Transforming protein RhoA OS=Homo sapiens GN=RHOA PE=1 SV=1 | TRUE | FALSE | FALSE | FALSE | FALSE |
| | P16070 | CD44_HUMAN | 141.89 | 5710000 | CD44 antigen OS=Homo sapiens GN=CD44 PE=1 SV=3 | TRUE | FALSE | FALSE | FALSE | FALSE |
| | Q9H444 | CHM4B_HUMAN | 95.61 | 5080000 | Charged multivesicular body protein 4b OS=Homo sapiens GN=CHMP4B PE=1 SV=1 | TRUE | FALSE | FALSE | FALSE | FALSE |
| | P32004 | L1CAM_HUMAN | 134.29 | 5020000 | Neural cell adhesion molecule L1 OS=Homo sapiens GN=L1CAM PE=1 SV=2 | TRUE | TRUE | TRUE | FALSE | FALSE |
| | O95274 | LYPD3_HUMAN | 141.14 | 4970000 | Ly6/PLAUR domain-containing protein 3 OS=Homo sapiens GN=LYPD3 PE=1 SV=2 | TRUE | FALSE | FALSE | FALSE | FALSE |
| | P07437 | TBB5_HUMAN | 129.88 | 4670000 | Tubulin beta chain OS=Homo sapiens GN=TUBB PE=1 SV=2 | TRUE | FALSE | FALSE | FALSE | FALSE |
| | P15311 | EZRI_HUMAN | 129.76 | 4090000 | Ezrin OS=Homo sapiens GN=EZR PE=1 SV=4 | TRUE | TRUE | TRUE | FALSE | FALSE |
| | P08754 | GNAI3_HUMAN | 111.38 | 4060000 | Guanine nucleotide-binding protein G(k) subunit alpha OS=Homo sapiens GN=GNAI3 PE=1 SV=3 | TRUE | FALSE | FALSE | FALSE | FALSE |
| | P0DMV8 | HS71A_HUMAN | 118.29 | 3530000 | Heat shock 70 kDa protein 1A OS=Homo sapiens GN=HSPA1A PE=1 SV=1 | TRUE | TRUE | TRUE | FALSE | FALSE |
| | P0DMV9 | HS71B_HUMAN | 118.29 | 3530000 | Heat shock 70 kDa protein 1B OS=Homo sapiens GN=HSPA1B PE=1 SV=1 | TRUE | TRUE | TRUE | FALSE | FALSE |
| | P10809 | CH60_HUMAN | 114.89 | 3020000 | 60 kDa heat shock protein mitochondrial OS=Homo sapiens GN=HSPD1 PE=1 SV=2 | TRUE | FALSE | FALSE | FALSE | FALSE |
| | O00299 | CLIC1_HUMAN | 96.21 | 2990000 | Chloride intracellular channel protein 1 OS=Homo sapiens GN=CLIC1 PE=1 SV=4 | TRUE | FALSE | FALSE | FALSE | FALSE |
| | P15328 | FOLR1_HUMAN | 80.6 | 2540000 | Folate receptor alpha OS=Homo sapiens GN=FOLR1 PE=1 SV=3 | TRUE | FALSE | FALSE | FALSE | FALSE |
| | P29401 | TKT_HUMAN | 71.68 | 2340000 | Transketolase OS=Homo sapiens GN=TKT PE=1 SV=3 | TRUE | FALSE | FALSE | FALSE | FALSE |
| | O00159 | MYO1C_HUMAN | 85.9 | 1650000 | Unconventional myosin-Ic OS=Homo sapiens GN=MYO1C PE=1 SV=4 | TRUE | FALSE | FALSE | FALSE | FALSE |
| | P08133 | ANXA6_HUMAN | 91.15 | 1380000 | Annexin A6 OS=Homo sapiens GN=ANXA6 PE=1 SV=3 | TRUE | FALSE | FALSE | FALSE | FALSE |
| | P07195 | LDHB_HUMAN | 156.21 | 991000 | L-lactate dehydrogenase B chain OS=Homo sapiens GN=LDHB PE=1 SV=2 | TRUE | TRUE | TRUE | FALSE | FALSE |
| | P07355 | ANXA2_HUMAN | 297.56 | 371000000 | Annexin A2 OS=Homo sapiens GN=ANXA2 PE=1 SV=2 | FALSE | FALSE | TRUE | TRUE | TRUE |
| | Q01650 | LAT1_HUMAN | 198.82 | 54100000 | Large neutral amino acids transporter small subunit 1 OS=Homo sapiens GN=SLC7A5 PE=1 SV=2 | FALSE | FALSE | FALSE | FALSE | FALSE |
| | P13647 | K2C5_HUMAN | 215.06 | 51100000 | Keratin type II cytoskeletal 5 OS=Homo sapiens GN=KRT5 PE=1 SV=3 | FALSE | FALSE | TRUE | FALSE | TRUE |
| | P01891 | 1A68_HUMAN | 159.03 | 28100000 | HLA class I histocompatibility antigen A-68 alpha chain OS=Homo sapiens GN=HLA-A PE=1 SV=4 | FALSE | FALSE | FALSE | FALSE | FALSE |
| | A0A0U1RRH7 | A0A0U1RRH7_HUMAN | 58.84 | 23000000 | Histone H2A OS=Homo sapiens PE=3 SV=1 | FALSE | FALSE | TRUE | FALSE | FALSE |
| | P04908 | H2A1B_HUMAN | 58.84 | 23000000 | Histone H2A type 1-B/E OS=Homo sapiens GN=HIST1H2AB PE=1 SV=2 | FALSE | FALSE | TRUE | FALSE | FALSE |
| | P0C0S5 | H2AZ_HUMAN | 58.84 | 23000000 | Histone H2A.Z OS=Homo sapiens GN=H2AFZ PE=1 SV=2 | FALSE | FALSE | TRUE | FALSE | FALSE |
| | P0C0S8 | H2A1_HUMAN | 58.84 | 23000000 | Histone H2A type 1 OS=Homo sapiens GN=HIST1H2AG PE=1 SV=2 | FALSE | FALSE | TRUE | FALSE | FALSE |
| | P16104 | H2AX_HUMAN | 58.84 | 23000000 | Histone H2AX OS=Homo sapiens GN=H2AFX PE=1 SV=2 | FALSE | FALSE | TRUE | FALSE | FALSE |
| | P20671 | H2A1D_HUMAN | 58.84 | 23000000 | Histone H2A type 1-D OS=Homo sapiens GN=HIST1H2AD PE=1 SV=2 | FALSE | FALSE | TRUE | FALSE | FALSE |
| | Q16777 | H2A2C_HUMAN | 58.84 | 23000000 | Histone H2A type 2-C OS=Homo sapiens GN=HIST2H2AC PE=1 SV=4 | FALSE | FALSE | TRUE | FALSE | FALSE |
| | Q6FI13 | H2A2A_HUMAN | 58.84 | 23000000 | Histone H2A type 2-A OS=Homo sapiens GN=HIST2H2AA3 PE=1 SV=3 | FALSE | FALSE | TRUE | FALSE | FALSE |

| Human from infected extracellular microvesicles (ImVs) | | | | | | | | | | |
|---|---|---|---|---|---|---|---|---|---|---|
| | Accession # | Protein Abbreviation | -10lgP | Area from one run (IP3_030917_3reps) | Description | Present in both repeats? | Present in both ImV repeats AND in Iexo? | in ImV and Iexo? | In ImV and MmV? | In ImV and Mexo? |
| | Q71UI9 | H2AV_HUMAN | 58.84 | 23000000 | Histone H2A.V OS=Homo sapiens GN=H2AFV PE=1 SV=3 | FALSE | FALSE | TRUE | FALSE | FALSE |
| | Q7L7L0 | H2A3_HUMAN | 58.84 | 23000000 | Histone H2A type 3 OS=Homo sapiens GN=HIST3H2A PE=1 SV=3 | FALSE | FALSE | TRUE | FALSE | FALSE |
| | Q93077 | H2A1C_HUMAN | 58.84 | 23000000 | Histone H2A type 1-C OS=Homo sapiens GN=HIST1H2AC PE=1 SV=3 | FALSE | FALSE | TRUE | FALSE | FALSE |
| | Q96KK5 | H2A1H_HUMAN | 58.84 | 23000000 | Histone H2A type 1-H OS=Homo sapiens GN=HIST1H2AH PE=1 SV=3 | FALSE | FALSE | TRUE | FALSE | FALSE |
| | Q96QV6 | H2A1A_HUMAN | 58.84 | 23000000 | Histone H2A type 1-A OS=Homo sapiens GN=HIST1H2AA PE=1 SV=3 | FALSE | FALSE | TRUE | FALSE | FALSE |
| | Q99878 | H2A1J_HUMAN | 58.84 | 23000000 | Histone H2A type 1-J OS=Homo sapiens GN=HIST1H2AJ PE=1 SV=3 | FALSE | FALSE | TRUE | FALSE | FALSE |
| | Q9BTM1 | H2AJ_HUMAN | 58.84 | 23000000 | Histone H2A.J OS=Homo sapiens GN=H2AFJ PE=1 SV=1 | FALSE | FALSE | TRUE | FALSE | FALSE |
| | P60660 | MYL6_HUMAN | 125.92 | 20000000 | Myosin light polypeptide 6 OS=Homo sapiens GN=MYL6 PE=1 SV=2 | FALSE | FALSE | FALSE | FALSE | FALSE |
| | P62328 | TYB4_HUMAN | 110.67 | 16700000 | Thymosin beta-4 OS=Homo sapiens GN=TMSB4X PE=1 SV=2 | FALSE | FALSE | FALSE | FALSE | FALSE |
| | P37802 | TAGL2_HUMAN | 147.46 | 16400000 | Transgelin-2 OS=Homo sapiens GN=TAGLN2 PE=1 SV=3 | FALSE | FALSE | TRUE | FALSE | FALSE |
| | P0CG47 | UBB_HUMAN | 106.86 | 15600000 | Polyubiquitin-B OS=Homo sapiens GN=UBB PE=1 SV=1 | FALSE | FALSE | FALSE | FALSE | FALSE |
| | Q5T749 | KPRP_HUMAN | 97.43 | 14700000 | Keratinocyte proline-rich protein OS=Homo sapiens GN=KPRP PE=1 SV=1 | FALSE | FALSE | FALSE | FALSE | TRUE |
| | P60981 | DEST_HUMAN | 121.27 | 13100000 | Destrin OS=Homo sapiens GN=DSTN PE=1 SV=3 | FALSE | FALSE | FALSE | FALSE | FALSE |
|  | P08779 | K1C16_HUMAN | 223.24 | 12300000 | Keratin type I cytoskeletal 16 OS=Homo sapiens GN=KRT16 PE=1 SV=4 | FALSE | FALSE | FALSE | FALSE | TRUE |
|  | P13807 | GYS1_HUMAN | 142.78 | 11300000 | Glycogen [starch] synthase muscle OS=Homo sapiens GN=GYS1 PE=1 SV=2 | FALSE | FALSE | FALSE | FALSE | FALSE |
| | P15531 | NDKA_HUMAN | 94.23 | 10600000 | Nucleoside diphosphate kinase A OS=Homo sapiens GN=NME1 PE=1 SV=1 | FALSE | FALSE | FALSE | FALSE | FALSE |
| | P22392 | NDKB_HUMAN | 94.23 | 10600000 | Nucleoside diphosphate kinase B OS=Homo sapiens GN=NME2 PE=1 SV=1 | FALSE | FALSE | FALSE | FALSE | FALSE |
| | Q92734 | TFG_HUMAN | 129.65 | 10100000 | Protein TFG OS=Homo sapiens GN=TFG PE=1 SV=2 | FALSE | FALSE | FALSE | FALSE | FALSE |
| | P02656 | APOC3_HUMAN | 121.2 | 10100000 | Apolipoprotein C-III OS=Homo sapiens GN=APOC3 PE=1 SV=1 | FALSE | FALSE | FALSE | FALSE | FALSE |
| | P62805 | H4_HUMAN | 101.3 | 10100000 | Histone H4 OS=Homo sapiens GN=HIST1H4A PE=1 SV=2 | FALSE | FALSE | FALSE | FALSE | FALSE |
| | P31944 | CASPE_HUMAN | 97.43 | 9750000 | Caspase-14 OS=Homo sapiens GN=CASP14 PE=1 SV=2 | FALSE | FALSE | FALSE | FALSE | FALSE |
| | P07737 | PROF1_HUMAN | 134.06 | 9280000 | Profilin-1 OS=Homo sapiens GN=PFN1 PE=1 SV=2 | FALSE | FALSE | FALSE | FALSE | FALSE |
| | P63313 | TYB10_HUMAN | 102.53 | 7630000 | Thymosin beta-10 OS=Homo sapiens GN=TMSB10 PE=1 SV=2 | FALSE | FALSE | FALSE | FALSE | FALSE |
| | P06753 | TPM3_HUMAN | 118.93 | 7520000 | Tropomyosin alpha-3 chain OS=Homo sapiens GN=TPM3 PE=1 SV=2 | FALSE | FALSE | FALSE | FALSE | FALSE |
| | P05362 | ICAM1_HUMAN | 83.81 | 7490000 | Intercellular adhesion molecule 1 OS=Homo sapiens GN=ICAM1 PE=1 SV=2 | FALSE | FALSE | FALSE | FALSE | FALSE |
| | Q5T750 | XP32_HUMAN | 85.97 | 7140000 | Skin-specific protein 32 OS=Homo sapiens GN=XP32 PE=1 SV=1 | FALSE | FALSE | FALSE | FALSE | FALSE |
| | P21926 | CD9_HUMAN | 86.7 | 6750000 | CD9 antigen OS=Homo sapiens GN=CD9 PE=1 SV=4 | FALSE | FALSE | FALSE | FALSE | FALSE |
| | Q8NBI5 | S43A3_HUMAN | 87.96 | 6560000 | Solute carrier family 43 member 3 OS=Homo sapiens GN=SLC43A3 PE=1 SV=2 | FALSE | FALSE | FALSE | FALSE | FALSE |
| | P43121 | MUC18_HUMAN | 119.16 | 6460000 | Cell surface glycoprotein MUC18 OS=Homo sapiens GN=MCAM PE=1 SV=2 | FALSE | FALSE | FALSE | FALSE | FALSE |
| | P00558 | PGK1_HUMAN | 146.7 | 6240000 | Phosphoglycerate kinase 1 OS=Homo sapiens GN=PGK1 PE=1 SV=3 | FALSE | FALSE | FALSE | FALSE | FALSE |
| | P05026 | AT1B1_HUMAN | 89.38 | 6060000 | Sodium/potassium-transporting ATPase subunit beta-1 OS=Homo sapiens GN=ATP1B1 PE=1 SV=1 | FALSE | FALSE | FALSE | FALSE | FALSE |
| | P06703 | S10A6_HUMAN | 63.9 | 6060000 | Protein S100-A6 OS=Homo sapiens GN=S100A6 PE=1 SV=1 | FALSE | FALSE | FALSE | FALSE | FALSE |
| | P61981 | 1433G_HUMAN | 127.58 | 5850000 | 14-3-3 protein gamma OS=Homo sapiens GN=YWHAG PE=1 SV=2 | FALSE | FALSE | TRUE | FALSE | FALSE |
| | P08582 | TRFM_HUMAN | 154.17 | 5300000 | Melanotransferrin OS=Homo sapiens GN=MELTF PE=1 SV=2 | FALSE | FALSE | FALSE | FALSE | FALSE |
| | O15427 | MOT4_HUMAN | 84.26 | 4970000 | Monocarboxylate transporter 4 OS=Homo sapiens GN=SLC16A3 PE=1 SV=1 | FALSE | FALSE | FALSE | FALSE | FALSE |
| | P29966 | MARCS_HUMAN | 115.98 | 4240000 | Myristoylated alanine-rich C-kinase substrate OS=Homo sapiens GN=MARCKS PE=1 SV=4 | FALSE | FALSE | FALSE | FALSE | FALSE |
| | P14174 | MIF_HUMAN | 73.81 | 4220000 | Macrophage migration inhibitory factor OS=Homo sapiens GN=MIF PE=1 SV=4 | FALSE | FALSE | FALSE | FALSE | FALSE |
| | P40121 | CAPG_HUMAN | 89.22 | 4190000 | Macrophage-capping protein OS=Homo sapiens GN=CAPG PE=1 SV=2 | FALSE | FALSE | FALSE | FALSE | FALSE |
| | P05089 | ARGI1_HUMAN | 88.17 | 3700000 | Arginase-1 OS=Homo sapiens GN=ARG1 PE=1 SV=2 | FALSE | FALSE | FALSE | FALSE | FALSE |
| | P22626 | ROA2_HUMAN | 107.87 | 3600000 | Heterogeneous nuclear ribonucleoproteins A2/B1 OS=Homo sapiens GN=HNRNPA2B1 PE=1 SV=2 | FALSE | FALSE | TRUE | FALSE | FALSE |

| Human from infected extracellular microvesicles (ImVs) | | | | | | | | | | |
|---|---|---|---|---|---|---|---|---|---|---|
| | Accession # | Protein Abbreviation | -10lgP | Area from one run (IP3_030917_3reps) | Description | Present in both repeats? | Present in both ImV repeats AND in Iexo? | in ImV and Iexo? | In ImV and MmV? | In ImV and Mexo? |
| | P26447 | S10A4_HUMAN | 89.16 | 3560000 | Protein S100-A4 OS=Homo sapiens GN=S100A4 PE=1 SV=1 | FALSE | FALSE | TRUE | FALSE | FALSE |
| | P06241 | FYN_HUMAN | 75.07 | 3290000 | Tyrosine-protein kinase Fyn OS=Homo sapiens GN=FYN PE=1 SV=3 | FALSE | FALSE | FALSE | FALSE | FALSE |
| | P07947 | YES_HUMAN | 75.07 | 3290000 | Tyrosine-protein kinase Yes OS=Homo sapiens GN=YES1 PE=1 SV=3 | FALSE | FALSE | FALSE | FALSE | FALSE |
| | Q5JWF2 | GNAS1_HUMAN | 131.29 | 3140000 | Guanine nucleotide-binding protein G(s) subunit alpha isoforms XLas OS=Homo sapiens GN=GNAS PE=1 SV=2 | FALSE | FALSE | FALSE | FALSE | FALSE |
| | P30041 | PRDX6_HUMAN | 88.87 | 3130000 | Peroxiredoxin-6 OS=Homo sapiens GN=PRDX6 PE=1 SV=3 | FALSE | FALSE | TRUE | FALSE | FALSE |
| | P01009 | A1AT_HUMAN | 99.4 | 3090000 | Alpha-1-antitrypsin OS=Homo sapiens GN=SERPINA1 PE=1 SV=3 | FALSE | FALSE | FALSE | FALSE | FALSE |
| | P49006 | MRP_HUMAN | 101.97 | 2890000 | MARCKS-related protein OS=Homo sapiens GN=MARCKSL1 PE=1 SV=2 | FALSE | FALSE | FALSE | FALSE | FALSE |
| | P14923 | PLAK_HUMAN | 76.81 | 2850000 | Junction plakoglobin OS=Homo sapiens GN=JUP PE=1 SV=3 | FALSE | FALSE | FALSE | FALSE | TRUE |
| | P17931 | LEG3_HUMAN | 82.69 | 2630000 | Galectin-3 OS=Homo sapiens GN=LGALS3 PE=1 SV=5 | FALSE | FALSE | FALSE | FALSE | FALSE |
| | P21333 | FLNA_HUMAN | 96.83 | 2600000 | Filamin-A OS=Homo sapiens GN=FLNA PE=1 SV=4 | FALSE | FALSE | FALSE | FALSE | FALSE |
| | Q9UBI6 | GBG12_HUMAN | 72.29 | 2440000 | Guanine nucleotide-binding protein G(I)/G(S)/G(O) subunit gamma-12 OS=Homo sapiens GN=GNG12 PE=1 SV=3 | FALSE | FALSE | TRUE | FALSE | FALSE |
| | P10909 | CLUS_HUMAN | 91.3 | 1960000 | Clusterin OS=Homo sapiens GN=CLU PE=1 SV=1 | FALSE | FALSE | FALSE | FALSE | TRUE |
| | P61604 | CH10_HUMAN | 115.57 | 1940000 | 10 kDa heat shock protein mitochondrial OS=Homo sapiens GN=HSPE1 PE=1 SV=2 | FALSE | FALSE | FALSE | FALSE | FALSE |
| | P53990 | IST1_HUMAN | 74.95 | 1640000 | IST1 homolog OS=Homo sapiens GN=IST1 PE=1 SV=1 | FALSE | FALSE | FALSE | FALSE | FALSE |
| | O00560 | SDCB1_HUMAN | 64.61 | 1590000 | Syntenin-1 OS=Homo sapiens GN=SDCBP PE=1 SV=1 | FALSE | FALSE | FALSE | FALSE | FALSE |
| | P62258 | 1433E_HUMAN | 82.54 | 1260000 | 14-3-3 protein epsilon OS=Homo sapiens GN=YWHAE PE=1 SV=1 | FALSE | FALSE | TRUE | FALSE | FALSE |
| | P68431 | H31_HUMAN | 64.19 | 1050000 | Histone H3.1 OS=Homo sapiens GN=HIST1H3A PE=1 SV=2 | FALSE | FALSE | FALSE | FALSE | FALSE |
| | Q16695 | H31T_HUMAN | 64.19 | 1050000 | Histone H3.1t OS=Homo sapiens GN=HIST3H3 PE=1 SV=3 | FALSE | FALSE | FALSE | FALSE | FALSE |
| | Q71DI3 | H32_HUMAN | 64.19 | 1050000 | Histone H3.2 OS=Homo sapiens GN=HIST2H3A PE=1 SV=3 | FALSE | FALSE | FALSE | FALSE | FALSE |
| | Q92928 | RAB1C_HUMAN | 42.87 | 976000 | Putative Ras-related protein Rab-1C OS=Homo sapiens GN=RAB1C PE=5 SV=2 | FALSE | FALSE | FALSE | FALSE | FALSE |
| | Q9H0U4 | RAB1B_HUMAN | 42.87 | 976000 | Ras-related protein Rab-1B OS=Homo sapiens GN=RAB1B PE=1 SV=1 | FALSE | FALSE | FALSE | FALSE | FALSE |
| | P15924 | DESP_HUMAN | 68.18 | 689000 | Desmoplakin OS=Homo sapiens GN=DSP PE=1 SV=3 | FALSE | FALSE | FALSE | FALSE | TRUE |
| | P31946 | 1433B_HUMAN | 98.9 | 624000 | 14-3-3 protein beta/alpha OS=Homo sapiens GN=YWHAB PE=1 SV=3 | FALSE | FALSE | FALSE | FALSE | FALSE |
| | P62269 | RS18_HUMAN | 64.29 | 600000 | 40S ribosomal protein S18 OS=Homo sapiens GN=RPS18 PE=1 SV=3 | FALSE | FALSE | TRUE | FALSE | FALSE |
| | P05783 | K1C18_HUMAN | 71.69 | 577000 | Keratin type I cytoskeletal 18 OS=Homo sapiens GN=KRT18 PE=1 SV=2 | FALSE | FALSE | TRUE | FALSE | FALSE |
| | P62826 | RAN_HUMAN | 41.68 | 543000 | GTP-binding nuclear protein Ran OS=Homo sapiens GN=RAN PE=1 SV=3 | FALSE | FALSE | FALSE | FALSE | FALSE |
| | P02763 | A1AG1_HUMAN | 80.6 | 529000 | Alpha-1-acid glycoprotein 1 OS=Homo sapiens GN=ORM1 PE=1 SV=1 | FALSE | FALSE | FALSE | FALSE | FALSE |
| | P06744 | G6PI_HUMAN | 70.33 | 521000 | Glucose-6-phosphate isomerase OS=Homo sapiens GN=GPI PE=1 SV=4 | FALSE | FALSE | FALSE | FALSE | FALSE |
| | P36578 | RL4_HUMAN | 92.29 | 520000 | 60S ribosomal protein L4 OS=Homo sapiens GN=RPL4 PE=1 SV=5 | FALSE | FALSE | TRUE | FALSE | FALSE |
| | P12429 | ANXA3_HUMAN | 60.1 | 511000 | Annexin A3 OS=Homo sapiens GN=ANXA3 PE=1 SV=3 | FALSE | FALSE | FALSE | FALSE | FALSE |
| | P61769 | B2MG_HUMAN | 59.81 | 494000 | Beta-2-microglobulin OS=Homo sapiens GN=B2M PE=1 SV=1 | FALSE | FALSE | FALSE | FALSE | FALSE |
| | P60763 | RAC3_HUMAN | 69.72 | 429000 | Ras-related C3 botulinum toxin substrate 3 OS=Homo sapiens GN=RAC3 PE=1 SV=1 | FALSE | FALSE | FALSE | FALSE | FALSE |
| | P63000 | RAC1_HUMAN | 69.72 | 429000 | Ras-related C3 botulinum toxin substrate 1 OS=Homo sapiens GN=RAC1 PE=1 SV=1 | FALSE | FALSE | FALSE | FALSE | FALSE |
| | P62899 | RL31_HUMAN | 59 | 413000 | 60S ribosomal protein L31 OS=Homo sapiens GN=RPL31 PE=1 SV=1 | FALSE | FALSE | FALSE | FALSE | FALSE |
| | Q00839 | HNRPU_HUMAN | 63.56 | 393000 | Heterogeneous nuclear ribonucleoprotein U OS=Homo sapiens GN=HNRNPU PE=1 SV=6 | FALSE | FALSE | FALSE | FALSE | FALSE |
| | P06748 | NPM_HUMAN | 74.72 | 367000 | Nucleophosmin OS=Homo sapiens GN=NPM1 PE=1 SV=2 | FALSE | FALSE | TRUE | FALSE | FALSE |
| | Q02878 | RL6_HUMAN | 71.29 | 341000 | 60S ribosomal protein L6 OS=Homo sapiens GN=RPL6 PE=1 SV=3 | FALSE | FALSE | TRUE | FALSE | FALSE |
| | P31327 | CPSM_HUMAN | 82.49 | 337000 | Carbamoyl-phosphate synthase [ammonia] mitochondrial OS=Homo sapiens GN=CPS1 PE=1 SV=2 | FALSE | FALSE | FALSE | FALSE | FALSE |
| | P62906 | RL10A_HUMAN | 52.7 | 320000 | 60S ribosomal protein L10a OS=Homo sapiens GN=RPL10A PE=1 SV=2 | FALSE | FALSE | TRUE | FALSE | FALSE |

| Human from infected extracellular microvesicles (ImVs) | | | | | | | | | | |
|---|---|---|---|---|---|---|---|---|---|---|
| | Accession # | Protein Abbreviation | -10lgP | Area from one run (IP3_030917_ 3reps) | Description | Present in both repeats? | Present in both ImV repeats AND in Iexo? | in ImV and Iexo? | In ImV and MmV? | In ImV and Mexo? |
| | P11021 | GRP78_HUMAN | 38.77 | 319000 | 78 kDa glucose-regulated protein OS=Homo sapiens GN=HSPA5 PE=1 SV=2 | FALSE | FALSE | FALSE | FALSE | FALSE |
| | Q99829 | CPNE1_HUMAN | 64.35 | 312000 | Copine-1 OS=Homo sapiens GN=CPNE1 PE=1 SV=1 | FALSE | FALSE | FALSE | FALSE | FALSE |
| | P13639 | EF2_HUMAN | 77.29 | 237000 | Elongation factor 2 OS=Homo sapiens GN=EEF2 PE=1 SV=4 | FALSE | FALSE | TRUE | FALSE | FALSE |
| | O75347 | TBCA_HUMAN | 60.23 | 223000 | Tubulin-specific chaperone A OS=Homo sapiens GN=TBCA PE=1 SV=3 | FALSE | FALSE | FALSE | FALSE | FALSE |
| | P07954 | FUMH_HUMAN | 52.6 | 215000 | Fumarate hydratase mitochondrial OS=Homo sapiens GN=FH PE=1 SV=3 | FALSE | FALSE | FALSE | FALSE | FALSE |
| | P62701 | RS4X_HUMAN | 39.29 | 215000 | 40S ribosomal protein S4 X isoform OS=Homo sapiens GN=RPS4X PE=1 SV=2 | FALSE | FALSE | TRUE | FALSE | TRUE |
| | P08134 | RHOC_HUMAN | 53.55 | 189000 | Rho-related GTP-binding protein RhoC OS=Homo sapiens GN=RHOC PE=1 SV=1 | FALSE | FALSE | FALSE | FALSE | FALSE |
| | O75390 | CISY_HUMAN | 34.66 | 181000 | Citrate synthase mitochondrial OS=Homo sapiens GN=CS PE=1 SV=2 | FALSE | FALSE | FALSE | FALSE | FALSE |
| | P46940 | IQGA1_HUMAN | 62.74 | 176000 | Ras GTPase-activating-like protein IQGAP1 OS=Homo sapiens GN=IQGAP1 PE=1 SV=1 | FALSE | FALSE | FALSE | FALSE | FALSE |
| | Q07020 | RL18_HUMAN | 62.96 | 167000 | 60S ribosomal protein L18 OS=Homo sapiens GN=RPL18 PE=1 SV=2 | FALSE | FALSE | TRUE | FALSE | FALSE |
| | P25705 | ATPA_HUMAN | 44.65 | 153000 | ATP synthase subunit alpha mitochondrial OS=Homo sapiens GN=ATP5A1 PE=1 SV=1 | FALSE | FALSE | FALSE | FALSE | FALSE |
| | P32119 | PRDX2_HUMAN | 69.68 | 143000 | Peroxiredoxin-2 OS=Homo sapiens GN=PRDX2 PE=1 SV=5 | FALSE | FALSE | FALSE | FALSE | FALSE |
| | | | | | # True: | 65 | 36 | 68 | 12 | 23 |
| | | | | | | | just Iexo: | 100 | | |
| | | | | | 65 proteins in both ImV repeats, out of 168 total above the 0.999 threshold (39%) | | | | | |
| | | | | | 162 proteins in Iexo above the 0.999 threshold | | | | | |
| | | | | | 36 proteins in both ImV and in Iexo (36/65 = 55% of those in both repeats are in Iexos, or 36/168=21% of any IEVs are in Iexos) | | | | | |
| | | | | | 68 proteins in one ImV and in Iexo | | | | | |
| | | | | | 11 proteins in both ImV repeats and MmV (11% of both ImV expts), and 1 more in one ImV and MmV (7% of proteins in at least one ImV expt are also found in MmV) | | | | | |
| | | | | | 15 proteins in both ImVs and Mexo 23% of both IEV expts), and 8 in one ImV and Mexo (14% of proteins in at least one ImV expt are also found in Mexo) | | | | | |

### Table S1B

| Human from Mock extracellular microvesicles (MmV) | | | | | | |
|---|---|---|---|---|---|---|
| | Accession # | Protein Abbreviation | -10lgP | Area | Description | Present in both repeats? |
| | P60709 | ACTB_HUMAN | 139.43 | 25400000 | Actin cytoplasmic 1 OS=Homo sapiens GN=ACTB PE=1 SV=1 | TRUE |
| | P63261 | ACTG_HUMAN | 139.43 | 25400000 | Actin cytoplasmic 2 OS=Homo sapiens GN=ACTG1 PE=1 SV=1 | TRUE |
| | P04264 | K2C1_HUMAN | 150.49 | 19900000 | Keratin type II cytoskeletal 1 OS=Homo sapiens GN=KRT1 PE=1 SV=6 | TRUE |
| | P13645 | K1C10_HUMAN | 141.52 | 14000000 | Keratin type I cytoskeletal 10 OS=Homo sapiens GN=KRT10 PE=1 SV=6 | TRUE |
| | P35908 | K22E_HUMAN | 115.88 | 4230000 | Keratin type II cytoskeletal 2 epidermal OS=Homo sapiens GN=KRT2 PE=1 SV=2 | TRUE |
| | O43854 | EDIL3_HUMAN | 121.46 | 36200000 | EGF-like repeat and discoidin I-like domain-containing protein 3 OS=Homo sapiens GN=EDIL3 PE=1 SV=1 | FALSE |
| | P35613 | BASI_HUMAN | 171.18 | 21700000 | Basigin OS=Homo sapiens GN=BSG PE=1 SV=2 | FALSE |
| | A6NMY6 | AXA2L_HUMAN | 119.11 | 11600000 | Putative annexin A2-like protein OS=Homo sapiens GN=ANXA2P2 PE=5 SV=2 | FALSE |
| | P07355 | ANXA2_HUMAN | 119.11 | 11600000 | Annexin A2 OS=Homo sapiens GN=ANXA2 PE=1 SV=2 | FALSE |
| | P35527 | K1C9_HUMAN | 166.72 | 8340000 | Keratin type I cytoskeletal 9 OS=Homo sapiens GN=KRT9 PE=1 SV=3 | FALSE |
| | P0CG48 | UBC_HUMAN | 69.46 | 3030000 | Polyubiquitin-C OS=Homo sapiens GN=UBC PE=1 SV=3 | FALSE |
| | P62979 | RS27A_HUMAN | 69.46 | 3030000 | Ubiquitin-40S ribosomal protein S27a OS=Homo sapiens GN=RPS27A PE=1 SV=2 | FALSE |
| | P62987 | RL40_HUMAN | 69.46 | 3030000 | Ubiquitin-60S ribosomal protein L40 OS=Homo sapiens GN=UBA52 PE=1 SV=2 | FALSE |
| | P08195 | 4F2_HUMAN | 91.91 | 1820000 | 4F2 cell-surface antigen heavy chain OS=Homo sapiens GN=SLC3A2 PE=1 SV=3 | FALSE |
| | | | | | # True: | 5 |

### Table S1C

| Human from infected CD9 exos (Iexo) | | | | | | |
|---|---|---|---|---|---|---|
| | Accession # | Protein Abbreviation | -10lgP | Area CD9_iexo_3reps | Description | In Iexo and Mexo? |
| | P13645 | K1C10_HUMAN | 224 | 54000000 | Keratin type I cytoskeletal 10 OS=Homo sapiens GN=KRT10 PE=1 SV=6 | TRUE |
| | P04264 | K2C1_HUMAN | 256.62 | 51200000 | Keratin type II cytoskeletal 1 OS=Homo sapiens GN=KRT1 PE=1 SV=6 | TRUE |
| | P60709 | ACTB_HUMAN | 225.17 | 42700000 | Actin cytoplasmic 1 OS=Homo sapiens GN=ACTB PE=1 SV=1 | TRUE |
| | P63261 | ACTG_HUMAN | 225.17 | 42700000 | Actin cytoplasmic 2 OS=Homo sapiens GN=ACTG1 PE=1 SV=1 | TRUE |
| | P23396 | RS3_HUMAN | 189.35 | 28000000 | 40S ribosomal protein S3 OS=Homo sapiens GN=RPS3 PE=1 SV=2 | TRUE |
| | P35527 | K1C9_HUMAN | 187.86 | 23500000 | Keratin type I cytoskeletal 9 OS=Homo sapiens GN=KRT9 PE=1 SV=3 | FALSE |
| | P80723 | BASP1_HUMAN | 193.06 | 20700000 | Brain acid soluble protein 1 OS=Homo sapiens GN=BASP1 PE=1 SV=2 | FALSE |
| | P62263 | RS14_HUMAN | 166.7 | 20600000 | 40S ribosomal protein S14 OS=Homo sapiens GN=RPS14 PE=1 SV=3 | FALSE |
| | P04406 | G3P_HUMAN | 181.15 | 19000000 | Glyceraldehyde-3-phosphate dehydrogenase OS=Homo sapiens GN=GAPDH PE=1 SV=3 | TRUE |
| | P61247 | RS3A_HUMAN | 181.98 | 19000000 | 40S ribosomal protein S3a OS=Homo sapiens GN=RPS3A PE=1 SV=2 | FALSE |
| | P62269 | RS18_HUMAN | 161.3 | 18500000 | 40S ribosomal protein S18 OS=Homo sapiens GN=RPS18 PE=1 SV=3 | FALSE |
| | P35908 | K22E_HUMAN | 230.65 | 17900000 | Keratin type II cytoskeletal 2 epidermal OS=Homo sapiens GN=KRT2 PE=1 SV=2 | FALSE |
| | P08865 | RSSA_HUMAN | 127.2 | 17700000 | 40S ribosomal protein SA OS=Homo sapiens GN=RPSA PE=1 SV=4 | TRUE |
| | P60866 | RS20_HUMAN | 161.54 | 17600000 | 40S ribosomal protein S20 OS=Homo sapiens GN=RPS20 PE=1 SV=1 | FALSE |
| | P19338 | NUCL_HUMAN | 167.18 | 17600000 | Nucleolin OS=Homo sapiens GN=NCL PE=1 SV=3 | FALSE |
| | P22626 | ROA2_HUMAN | 192.68 | 17000000 | Heterogeneous nuclear ribonucleoproteins A2/B1 OS=Homo sapiens GN=HNRNPA2B1 PE=1 SV=2 | FALSE |
| | P62081 | RS7_HUMAN | 115.28 | 16900000 | 40S ribosomal protein S7 OS=Homo sapiens GN=RPS7 PE=1 SV=1 | FALSE |
| | P15880 | RS2_HUMAN | 137.76 | 14800000 | 40S ribosomal protein S2 OS=Homo sapiens GN=RPS2 PE=1 SV=2 | FALSE |
| | P07355 | ANXA2_HUMAN | 158.99 | 13700000 | Annexin A2 OS=Homo sapiens GN=ANXA2 PE=1 SV=2 | TRUE |
| | P62701 | RS4X_HUMAN | 188.98 | 12800000 | 40S ribosomal protein S4 X isoform OS=Homo sapiens GN=RPS4X PE=1 SV=2 | TRUE |
| | O75340 | PDCD6_HUMAN | 166.78 | 12700000 | Programmed cell death protein 6 OS=Homo sapiens GN=PDCD6 PE=1 SV=1 | FALSE |
| | P63220 | RS21_HUMAN | 117.66 | 11800000 | 40S ribosomal protein S21 OS=Homo sapiens GN=RPS21 PE=1 SV=1 | FALSE |
| | P35613 | BASI_HUMAN | 196.01 | 10900000 | Basigin OS=Homo sapiens GN=BSG PE=1 SV=2 | TRUE |
| | P62244 | RS15A_HUMAN | 122.67 | 10600000 | 40S ribosomal protein S15a OS=Homo sapiens GN=RPS15A PE=1 SV=2 | FALSE |
| | P42677 | RS27_HUMAN | 139.52 | 10400000 | 40S ribosomal protein S27 OS=Homo sapiens GN=RPS27 PE=1 SV=3 | FALSE |
| | P62277 | RS13_HUMAN | 118.35 | 10000000 | 40S ribosomal protein S13 OS=Homo sapiens GN=RPS13 PE=1 SV=2 | FALSE |
| | P62280 | RS11_HUMAN | 168.65 | 9970000 | 40S ribosomal protein S11 OS=Homo sapiens GN=RPS11 PE=1 SV=3 | FALSE |
| | P08174 | DAF_HUMAN | 145.52 | 9660000 | Complement decay-accelerating factor OS=Homo sapiens GN=CD55 PE=1 SV=4 | FALSE |
| | P07910 | HNRPC_HUMAN | 115.77 | 9570000 | Heterogeneous nuclear ribonucleoproteins C1/C2 OS=Homo sapiens GN=HNRNPC PE=1 SV=4 | FALSE |
| | P39019 | RS19_HUMAN | 130.68 | 8990000 | 40S ribosomal protein S19 OS=Homo sapiens GN=RPS19 PE=1 SV=2 | FALSE |
| | P68104 | EF1A1_HUMAN | 111.7 | 8910000 | Elongation factor 1-alpha 1 OS=Homo sapiens GN=EEF1A1 PE=1 SV=1 | FALSE |
| | Q5VTE0 | EF1A3_HUMAN | 111.7 | 8910000 | Putative elongation factor 1-alpha-like 3 OS=Homo sapiens GN=EEF1A1P5 PE=5 SV=1 | FALSE |
| | P62753 | RS6_HUMAN | 139.43 | 7500000 | 40S ribosomal protein S6 OS=Homo sapiens GN=RPS6 PE=1 SV=1 | TRUE |

| Human from infected CD9 exos (Iexo) | | | | | | |
|---|---|---|---|---|---|---|
| | Accession # | Protein Abbreviation | -10lgP | Area CD9_iexo_3reps | Description | In Iexo and Mexo? |
| | P25398 | RS12_HUMAN | 149.95 | 6750000 | 40S ribosomal protein S12 OS=Homo sapiens GN=RPS12 PE=1 SV=3 | TRUE |
| | P62241 | RS8_HUMAN | 144.82 | 6260000 | 40S ribosomal protein S8 OS=Homo sapiens GN=RPS8 PE=1 SV=2 | FALSE |
| | P62841 | RS15_HUMAN | 95 | 5660000 | 40S ribosomal protein S15 OS=Homo sapiens GN=RPS15 PE=1 SV=2 | FALSE |
| | P08621 | RU17_HUMAN | 150.53 | 5370000 | U1 small nuclear ribonucleoprotein 70 kDa OS=Homo sapiens GN=SNRNP70 PE=1 SV=2 | FALSE |
| | P09651 | ROA1_HUMAN | 145.16 | 4920000 | Heterogeneous nuclear ribonucleoprotein A1 OS=Homo sapiens GN=HNRNPA1 PE=1 SV=5 | FALSE |
| | P49368 | TCPG_HUMAN | 145.3 | 4880000 | T-complex protein 1 subunit gamma OS=Homo sapiens GN=CCT3 PE=1 SV=4 | FALSE |
| | P46782 | RS5_HUMAN | 140.66 | 4860000 | 40S ribosomal protein S5 OS=Homo sapiens GN=RPS5 PE=1 SV=4 | FALSE |
| | P62851 | RS25_HUMAN | 110.79 | 4550000 | 40S ribosomal protein S25 OS=Homo sapiens GN=RPS25 PE=1 SV=1 | FALSE |
| | P46777 | RL5_HUMAN | 132.6 | 4390000 | 60S ribosomal protein L5 OS=Homo sapiens GN=RPL5 PE=1 SV=3 | FALSE |
| | P13987 | CD59_HUMAN | 142.28 | 4240000 | CD59 glycoprotein OS=Homo sapiens GN=CD59 PE=1 SV=1 | TRUE |
| | P62249 | RS16_HUMAN | 115.59 | 4130000 | 40S ribosomal protein S16 OS=Homo sapiens GN=RPS16 PE=1 SV=2 | FALSE |
| | P02794 | FRIH_HUMAN | 134.6 | 4110000 | Ferritin heavy chain OS=Homo sapiens GN=FTH1 PE=1 SV=2 | TRUE |
| | Q02878 | RL6_HUMAN | 130.56 | 3800000 | 60S ribosomal protein L6 OS=Homo sapiens GN=RPL6 PE=1 SV=3 | FALSE |
| | P09012 | SNRPA_HUMAN | 88.36 | 3600000 | U1 small nuclear ribonucleoprotein A OS=Homo sapiens GN=SNRPA PE=1 SV=3 | FALSE |
| | P30050 | RL12_HUMAN | 107.74 | 3460000 | 60S ribosomal protein L12 OS=Homo sapiens GN=RPL12 PE=1 SV=1 | FALSE |
| | P62750 | RL23A_HUMAN | 131.73 | 3340000 | 60S ribosomal protein L23a OS=Homo sapiens GN=RPL23A PE=1 SV=1 | FALSE |
| | P40227 | TCPZ_HUMAN | 130.82 | 3200000 | T-complex protein 1 subunit zeta OS=Homo sapiens GN=CCT6A PE=1 SV=3 | FALSE |
| | P50995 | ANX11_HUMAN | 147.88 | 3200000 | Annexin A11 OS=Homo sapiens GN=ANXA11 PE=1 SV=1 | FALSE |
| | Q99832 | TCPH_HUMAN | 141.47 | 3120000 | T-complex protein 1 subunit eta OS=Homo sapiens GN=CCT7 PE=1 SV=2 | FALSE |
| | P62979 | RS27A_HUMAN | 91.67 | 3060000 | Ubiquitin-40S ribosomal protein S27a OS=Homo sapiens GN=RPS27A PE=1 SV=2 | FALSE |
| | P04083 | ANXA1_HUMAN | 133.71 | 2910000 | Annexin A1 OS=Homo sapiens GN=ANXA1 PE=1 SV=2 | FALSE |
| | P78371 | TCPB_HUMAN | 144.48 | 2900000 | T-complex protein 1 subunit beta OS=Homo sapiens GN=CCT2 PE=1 SV=4 | FALSE |
| | P62917 | RL8_HUMAN | 111.55 | 2840000 | 60S ribosomal protein L8 OS=Homo sapiens GN=RPL8 PE=1 SV=2 | FALSE |
| | P08238 | HS90B_HUMAN | 173.46 | 2800000 | Heat shock protein HSP 90-beta OS=Homo sapiens GN=HSP90AB1 PE=1 SV=4 | FALSE |
| | P47914 | RL29_HUMAN | 87.87 | 2690000 | 60S ribosomal protein L29 OS=Homo sapiens GN=RPL29 PE=1 SV=2 | FALSE |
| | Q06830 | PRDX1_HUMAN | 134.76 | 2630000 | Peroxiredoxin-1 OS=Homo sapiens GN=PRDX1 PE=1 SV=1 | FALSE |
| | P30626 | SORCN_HUMAN | 109.95 | 2500000 | Sorcin OS=Homo sapiens GN=SRI PE=1 SV=1 | FALSE |
| | P62306 | RUXF_HUMAN | 106.11 | 2490000 | Small nuclear ribonucleoprotein F OS=Homo sapiens GN=SNRPF PE=1 SV=1 | FALSE |
| | P62316 | SMD2_HUMAN | 136.83 | 2420000 | Small nuclear ribonucleoprotein Sm D2 OS=Homo sapiens GN=SNRPD2 PE=1 SV=1 | FALSE |
| | P46783 | RS10_HUMAN | 117.88 | 2400000 | 40S ribosomal protein S10 OS=Homo sapiens GN=RPS10 PE=1 SV=1 | FALSE |
| | Q9NQ39 | RS10L_HUMAN | 117.88 | 2400000 | Putative 40S ribosomal protein S10-like OS=Homo sapiens GN=RPS10P5 PE=5 SV=1 | FALSE |
| | P31949 | S10AB_HUMAN | 89.89 | 2370000 | Protein S100-A11 OS=Homo sapiens GN=S100A11 PE=1 SV=2 | FALSE |
| | P26373 | RL13_HUMAN | 122.62 | 2370000 | 60S ribosomal protein L13 OS=Homo sapiens GN=RPL13 PE=1 SV=4 | FALSE |
| | Q53EL6 | PDCD4_HUMAN | 148.05 | 2290000 | Programmed cell death protein 4 OS=Homo sapiens GN=PDCD4 PE=1 SV=2 | FALSE |
| | P14618 | KPYM_HUMAN | 118.32 | 2260000 | Pyruvate kinase PKM OS=Homo sapiens GN=PKM PE=1 SV=4 | FALSE |

| Human from infected CD9 exos (Iexo) | | | | | | |
|---|---|---|---|---|---|---|
| | Accession # | Protein Abbreviation | -10lgP | Area CD9_iexo_3reps | Description | In Iexo and Mexo? |
| | P48643 | TCPE_HUMAN | 118.31 | 2250000 | T-complex protein 1 subunit epsilon OS=Homo sapiens GN=CCT5 PE=1 SV=1 | FALSE |
| | P46781 | RS9_HUMAN | 82.75 | 2220000 | 40S ribosomal protein S9 OS=Homo sapiens GN=RPS9 PE=1 SV=3 | FALSE |
| | P11142 | HSP7C_HUMAN | 148.48 | 2210000 | Heat shock cognate 71 kDa protein OS=Homo sapiens GN=HSPA8 PE=1 SV=1 | FALSE |
| | P18124 | RL7_HUMAN | 155.86 | 2160000 | 60S ribosomal protein L7 OS=Homo sapiens GN=RPL7 PE=1 SV=1 | FALSE |
| | P50914 | RL14_HUMAN | 99.82 | 2040000 | 60S ribosomal protein L14 OS=Homo sapiens GN=RPL14 PE=1 SV=4 | FALSE |
| | P62273 | RS29_HUMAN | 93.73 | 1960000 | 40S ribosomal protein S29 OS=Homo sapiens GN=RPS29 PE=1 SV=2 | FALSE |
| | A0A0U1RRH7 | A0A0U1RRH7_HUMAN | 54.8 | 1850000 | Histone H2A OS=Homo sapiens PE=3 SV=1 | FALSE |
| | P04908 | H2A1B_HUMAN | 54.8 | 1850000 | Histone H2A type 1-B/E OS=Homo sapiens GN=HIST1H2AB PE=1 SV=2 | FALSE |
| | P0C0S5 | H2AZ_HUMAN | 54.8 | 1850000 | Histone H2A.Z OS=Homo sapiens GN=H2AFZ PE=1 SV=2 | FALSE |
| | P0C0S8 | H2A1_HUMAN | 54.8 | 1850000 | Histone H2A type 1 OS=Homo sapiens GN=HIST1H2AG PE=1 SV=2 | FALSE |
| | P16104 | H2AX_HUMAN | 54.8 | 1850000 | Histone H2AX OS=Homo sapiens GN=H2AFX PE=1 SV=2 | FALSE |
| | P20671 | H2A1D_HUMAN | 54.8 | 1850000 | Histone H2A type 1-D OS=Homo sapiens GN=HIST1H2AD PE=1 SV=2 | FALSE |
| | Q16777 | H2A2C_HUMAN | 54.8 | 1850000 | Histone H2A type 2-C OS=Homo sapiens GN=HIST2H2AC PE=1 SV=4 | FALSE |
| | Q6FI13 | H2A2A_HUMAN | 54.8 | 1850000 | Histone H2A type 2-A OS=Homo sapiens GN=HIST2H2AA3 PE=1 SV=3 | FALSE |
| | Q71UI9 | H2AV_HUMAN | 54.8 | 1850000 | Histone H2A.V OS=Homo sapiens GN=H2AFV PE=1 SV=3 | FALSE |
| | Q7L7L0 | H2A3_HUMAN | 54.8 | 1850000 | Histone H2A type 3 OS=Homo sapiens GN=HIST3H2A PE=1 SV=3 | FALSE |
| | Q93077 | H2A1C_HUMAN | 54.8 | 1850000 | Histone H2A type 1-C OS=Homo sapiens GN=HIST1H2AC PE=1 SV=3 | FALSE |
| | Q96KK5 | H2A1H_HUMAN | 54.8 | 1850000 | Histone H2A type 1-H OS=Homo sapiens GN=HIST1H2AH PE=1 SV=3 | FALSE |
| | Q96QV6 | H2A1A_HUMAN | 54.8 | 1850000 | Histone H2A type 1-A OS=Homo sapiens GN=HIST1H2AA PE=1 SV=3 | FALSE |
| | Q99878 | H2A1J_HUMAN | 54.8 | 1850000 | Histone H2A type 1-J OS=Homo sapiens GN=HIST1H2AJ PE=1 SV=3 | FALSE |
| | Q9BTM1 | H2AJ_HUMAN | 54.8 | 1850000 | Histone H2A.J OS=Homo sapiens GN=H2AFJ PE=1 SV=1 | FALSE |
| | P50991 | TCPD_HUMAN | 137.39 | 1850000 | T-complex protein 1 subunit delta OS=Homo sapiens GN=CCT4 PE=1 SV=4 | FALSE |
| | P08195 | 4F2_HUMAN | 126.29 | 1810000 | 4F2 cell-surface antigen heavy chain OS=Homo sapiens GN=SLC3A2 PE=1 SV=3 | FALSE |
| | P62266 | RS23_HUMAN | 108 | 1780000 | 40S ribosomal protein S23 OS=Homo sapiens GN=RPS23 PE=1 SV=3 | FALSE |
| | P62258 | 1433E_HUMAN | 96.63 | 1770000 | 14-3-3 protein epsilon OS=Homo sapiens GN=YWHAE PE=1 SV=1 | FALSE |
| | P63244 | RACK1_HUMAN | 97.95 | 1770000 | Receptor of activated protein C kinase 1 OS=Homo sapiens GN=RACK1 PE=1 SV=3 | FALSE |
| | P04075 | ALDOA_HUMAN | 103.04 | 1770000 | Fructose-bisphosphate aldolase A OS=Homo sapiens GN=ALDOA PE=1 SV=2 | FALSE |
| | P23528 | COF1_HUMAN | 96.93 | 1710000 | Cofilin-1 OS=Homo sapiens GN=CFL1 PE=1 SV=3 | FALSE |
| | P50990 | TCPQ_HUMAN | 131.72 | 1650000 | T-complex protein 1 subunit theta OS=Homo sapiens GN=CCT8 PE=1 SV=4 | FALSE |
| | P39023 | RL3_HUMAN | 130.45 | 1630000 | 60S ribosomal protein L3 OS=Homo sapiens GN=RPL3 PE=1 SV=2 | FALSE |
| | P26038 | MOES_HUMAN | 142.8 | 1490000 | Moesin OS=Homo sapiens GN=MSN PE=1 SV=3 | FALSE |
| | P36578 | RL4_HUMAN | 127.9 | 1440000 | 60S ribosomal protein L4 OS=Homo sapiens GN=RPL4 PE=1 SV=5 | FALSE |
| | P11940 | PABP1_HUMAN | 106.29 | 1420000 | Polyadenylate-binding protein 1 OS=Homo sapiens GN=PABPC1 PE=1 SV=2 | FALSE |
| | P63104 | 1433Z_HUMAN | 117.16 | 1360000 | 14-3-3 protein zeta/delta OS=Homo sapiens GN=YWHAZ PE=1 SV=1 | FALSE |
| | O15372 | EIF3H_HUMAN | 83.01 | 1290000 | Eukaryotic translation initiation factor 3 subunit H OS=Homo sapiens GN=EIF3H PE=1 SV=1 | FALSE |
| | P62424 | RL7A_HUMAN | 106.67 | 1290000 | 60S ribosomal protein L7a OS=Homo sapiens GN=RPL7A PE=1 SV=2 | FALSE |
| | P05387 | RLA2_HUMAN | 71.34 | 1270000 | 60S acidic ribosomal protein P2 OS=Homo sapiens GN=RPLP2 PE=1 SV=1 | FALSE |
| | P05388 | RLA0_HUMAN | 126.01 | 1180000 | 60S acidic ribosomal protein P0 OS=Homo sapiens GN=RPLP0 PE=1 SV=1 | FALSE |

| Human from infected CD9 exos (Iexo) | | | | | | |
|---|---|---|---|---|---|---|
| | Accession # | Protein Abbreviation | -10lgP | Area CD9_iexo_3reps | Description | In Iexo and Mexo? |
| | Q9UBI6 | GBG12_HUMAN | 101.29 | 1140000 | Guanine nucleotide-binding protein G(I)/G(S)/G(O) subunit gamma-12 OS=Homo sapiens GN=GNG12 PE=1 SV=3 | FALSE |
| | P68363 | TBA1B_HUMAN | 86.92 | 1070000 | Tubulin alpha-1B chain OS=Homo sapiens GN=TUBA1B PE=1 SV=1 | FALSE |
| | Q71U36 | TBA1A_HUMAN | 86.92 | 1070000 | Tubulin alpha-1A chain OS=Homo sapiens GN=TUBA1A PE=1 SV=1 | FALSE |
| | Q9BQE3 | TBA1C_HUMAN | 86.92 | 1070000 | Tubulin alpha-1C chain OS=Homo sapiens GN=TUBA1C PE=1 SV=1 | FALSE |
| | Q07020 | RL18_HUMAN | 105.93 | 1070000 | 60S ribosomal protein L18 OS=Homo sapiens GN=RPL18 PE=1 SV=2 | FALSE |
| | P61978 | HNRPK_HUMAN | 118.86 | 1070000 | Heterogeneous nuclear ribonucleoprotein K OS=Homo sapiens GN=HNRNPK PE=1 SV=1 | FALSE |
| | P37802 | TAGL2_HUMAN | 120.68 | 1030000 | Transgelin-2 OS=Homo sapiens GN=TAGLN2 PE=1 SV=3 | FALSE |
| | P55072 | TERA_HUMAN | 87.55 | 997000 | Transitional endoplasmic reticulum ATPase OS=Homo sapiens GN=VCP PE=1 SV=4 | FALSE |
| | P61313 | RL15_HUMAN | 62.7 | 951000 | 60S ribosomal protein L15 OS=Homo sapiens GN=RPL15 PE=1 SV=2 | FALSE |
| | P07195 | LDHB_HUMAN | 90.53 | 930000 | L-lactate dehydrogenase B chain OS=Homo sapiens GN=LDHB PE=1 SV=2 | FALSE |
| | B5ME19 | EIFCL_HUMAN | 104.7 | 908000 | Eukaryotic translation initiation factor 3 subunit C-like protein OS=Homo sapiens GN=EIF3CL PE=3 SV=1 | FALSE |
| | Q99613 | EIF3C_HUMAN | 104.7 | 908000 | Eukaryotic translation initiation factor 3 subunit C OS=Homo sapiens GN=EIF3C PE=1 SV=1 | FALSE |
| | P13639 | EF2_HUMAN | 97.43 | 885000 | Elongation factor 2 OS=Homo sapiens GN=EEF2 PE=1 SV=4 | FALSE |
| | P35268 | RL22_HUMAN | 68.49 | 869000 | 60S ribosomal protein L22 OS=Homo sapiens GN=RPL22 PE=1 SV=2 | FALSE |
| | J3QQQ9 | J3QQQ9_HUMAN | 105.27 | 825000 | Uncharacterized protein OS=Homo sapiens PE=4 SV=1 | FALSE |
| | P61254 | RL26_HUMAN | 105.27 | 825000 | 60S ribosomal protein L26 OS=Homo sapiens GN=RPL26 PE=1 SV=1 | FALSE |
| | Q9UNX3 | RL26L_HUMAN | 105.27 | 825000 | 60S ribosomal protein L26-like 1 OS=Homo sapiens GN=RPL26L1 PE=1 SV=1 | FALSE |
| | P62829 | RL23_HUMAN | 89.02 | 742000 | 60S ribosomal protein L23 OS=Homo sapiens GN=RPL23 PE=1 SV=1 | FALSE |
| | P35637 | FUS_HUMAN | 48.77 | 664000 | RNA-binding protein FUS OS=Homo sapiens GN=FUS PE=1 SV=1 | FALSE |
| | P49207 | RL34_HUMAN | 64.94 | 655000 | 60S ribosomal protein L34 OS=Homo sapiens GN=RPL34 PE=1 SV=3 | FALSE |
|  | P05783 | K1C18_HUMAN | 94.85 | 631000 | Keratin type I cytoskeletal 18 OS=Homo sapiens GN=KRT18 PE=1 SV=2 | FALSE |
| | Q13347 | EIF3I_HUMAN | 77.05 | 627000 | Eukaryotic translation initiation factor 3 subunit I OS=Homo sapiens GN=EIF3I PE=1 SV=1 | FALSE |
| | P30041 | PRDX6_HUMAN | 87.23 | 567000 | Peroxiredoxin-6 OS=Homo sapiens GN=PRDX6 PE=1 SV=3 | FALSE |
| | P61353 | RL27_HUMAN | 67.05 | 553000 | 60S ribosomal protein L27 OS=Homo sapiens GN=RPL27 PE=1 SV=2 | FALSE |
| | P62847 | RS24_HUMAN | 75.67 | 553000 | 40S ribosomal protein S24 OS=Homo sapiens GN=RPS24 PE=1 SV=1 | FALSE |
|  | P13647 | K2C5_HUMAN | 174.85 | 553000 | Keratin type II cytoskeletal 5 OS=Homo sapiens GN=KRT5 PE=1 SV=3 | FALSE |
| | P46776 | RL27A_HUMAN | 87.13 | 550000 | 60S ribosomal protein L27a OS=Homo sapiens GN=RPL27A PE=1 SV=2 | FALSE |
| | P08708 | RS17_HUMAN | 69.48 | 539000 | 40S ribosomal protein S17 OS=Homo sapiens GN=RPS17 PE=1 SV=2 | FALSE |
| | P62906 | RL10A_HUMAN | 51.17 | 536000 | 60S ribosomal protein L10a OS=Homo sapiens GN=RPL10A PE=1 SV=2 | FALSE |
| | P32004 | L1CAM_HUMAN | 94.01 | 497000 | Neural cell adhesion molecule L1 OS=Homo sapiens GN=L1CAM PE=1 SV=2 | FALSE |
| | P18077 | RL35A_HUMAN | 82.15 | 482000 | 60S ribosomal protein L35a OS=Homo sapiens GN=RPL35A PE=1 SV=2 | FALSE |
| | P49327 | FAS_HUMAN | 94.13 | 481000 | Fatty acid synthase OS=Homo sapiens GN=FASN PE=1 SV=3 | FALSE |
| | P00966 | ASSY_HUMAN | 94.52 | 447000 | Argininosuccinate synthase OS=Homo sapiens GN=ASS1 PE=1 SV=2 | FALSE |
| | P06748 | NPM_HUMAN | 80.13 | 419000 | Nucleophosmin OS=Homo sapiens GN=NPM1 PE=1 SV=2 | FALSE |

| Human from infected CD9 exos (Iexo) | | | | | | |
|---|---|---|---|---|---|---|
| | Accession # | Protein Abbreviation | -10lgP | Area CD9_iexo_3reps | Description | In Iexo and Mexo? |
| | Q8NC51 | PAIRB_HUMAN | 79.54 | 397000 | Plasminogen activator inhibitor 1 RNA-binding protein OS=Homo sapiens GN=SERBP1 PE=1 SV=2 | FALSE |
| | P62937 | PPIA_HUMAN | 49.29 | 363000 | Peptidyl-prolyl cis-trans isomerase A OS=Homo sapiens GN=PPIA PE=1 SV=2 | FALSE |
| | P08758 | ANXA5_HUMAN | 63.95 | 343000 | Annexin A5 OS=Homo sapiens GN=ANXA5 PE=1 SV=2 | FALSE |
| | Q13435 | SF3B2_HUMAN | 78.27 | 327000 | Splicing factor 3B subunit 2 OS=Homo sapiens GN=SF3B2 PE=1 SV=2 | FALSE |
| | P09382 | LEG1_HUMAN | 57.92 | 274000 | Galectin-1 OS=Homo sapiens GN=LGALS1 PE=1 SV=2 | FALSE |
| | O14744 | ANM5_HUMAN | 96.51 | 257000 | Protein arginine N-methyltransferase 5 OS=Homo sapiens GN=PRMT5 PE=1 SV=4 | FALSE |
| | Q14152 | EIF3A_HUMAN | 73.4 | 251000 | Eukaryotic translation initiation factor 3 subunit A OS=Homo sapiens GN=EIF3A PE=1 SV=1 | FALSE |
| | Q15459 | SF3A1_HUMAN | 76.7 | 246000 | Splicing factor 3A subunit 1 OS=Homo sapiens GN=SF3A1 PE=1 SV=1 | FALSE |
| | P61981 | 1433G_HUMAN | 91.32 | 213000 | 14-3-3 protein gamma OS=Homo sapiens GN=YWHAG PE=1 SV=2 | FALSE |
| | P15311 | EZRI_HUMAN | 127.71 | 202000 | Ezrin OS=Homo sapiens GN=EZR PE=1 SV=4 | FALSE |
| | Q09666 | AHNK_HUMAN | 99.79 | 194000 | Neuroblast differentiation-associated protein AHNAK OS=Homo sapiens GN=AHNAK PE=1 SV=2 | FALSE |
| | P07900 | HS90A_HUMAN | 115.66 | 187000 | Heat shock protein HSP 90-alpha OS=Homo sapiens GN=HSP90AA1 PE=1 SV=5 | FALSE |
| | Q8IZ83 | A16A1_HUMAN | 50.25 | 164000 | Aldehyde dehydrogenase family 16 member A1 OS=Homo sapiens GN=ALDH16A1 PE=1 SV=2 | FALSE |
| | P26447 | S10A4_HUMAN | 43.79 | 145000 | Protein S100-A4 OS=Homo sapiens GN=S100A4 PE=1 SV=1 | FALSE |
| | P08670 | VIME_HUMAN | 73.52 | 135000 | Vimentin OS=Homo sapiens GN=VIM PE=1 SV=4 | FALSE |
| | P0DMV8 | HS71A_HUMAN | 104.08 | 109000 | Heat shock 70 kDa protein 1A OS=Homo sapiens GN=HSPA1A PE=1 SV=1 | FALSE |
| | P0DMV9 | HS71B_HUMAN | 104.08 | 109000 | Heat shock 70 kDa protein 1B OS=Homo sapiens GN=HSPA1B PE=1 SV=1 | FALSE |
| | Q9NR30 | DDX21_HUMAN | 52.05 | 108000 | Nucleolar RNA helicase 2 OS=Homo sapiens GN=DDX21 PE=1 SV=5 | FALSE |
| | Q12904 | AIMP1_HUMAN | 52.08 | 107000 | Aminoacyl tRNA synthase complex-interacting multifunctional protein 1 OS=Homo sapiens GN=AIMP1 PE=1 SV=2 | FALSE |
| | P53396 | ACLY_HUMAN | 43.33 | 81100 | ATP-citrate synthase OS=Homo sapiens GN=ACLY PE=1 SV=3 | FALSE |
| | Q08211 | DHX9_HUMAN | 74.22 | 69500 | ATP-dependent RNA helicase A OS=Homo sapiens GN=DHX9 PE=1 SV=4 | FALSE |
| | P78527 | PRKDC_HUMAN | 68.21 | 66400 | DNA-dependent protein kinase catalytic subunit OS=Homo sapiens GN=PRKDC PE=1 SV=3 | FALSE |
| | | | | # ribosomal: | 55 | |
| | | | | % ribosomal | 0.339506173 | |

### Table S1D

| Human from Mock CD9 exo (Mexo) | | | | | |
|---|---|---|---|---|---|
| | Accession # | Protein Abbreviation | -10lgP | Area CD9_mexo_3 reps | Description |
| | P13645 | K1C10_HUMAN | 334.05 | 375000000 | Keratin type I cytoskeletal 10 OS=Homo sapiens GN=KRT10 PE=1 SV=6 |
| | P04264 | K2C1_HUMAN | 392.16 | 359000000 | Keratin type II cytoskeletal 1 OS=Homo sapiens GN=KRT1 PE=1 SV=6 |
| | P35527 | K1C9_HUMAN | 304.91 | 219000000 | Keratin type I cytoskeletal 9 OS=Homo sapiens GN=KRT9 PE=1 SV=3 |
| | P35908 | K22E_HUMAN | 344.56 | 134000000 | Keratin type II cytoskeletal 2 epidermal OS=Homo sapiens GN=KRT2 PE=1 SV=2 |
| | Q08380 | LG3BP_HUMAN | 279.63 | 114000000 | Galectin-3-binding protein OS=Homo sapiens GN=LGALS3BP PE=1 SV=1 |
| | P02533 | K1C14_HUMAN | 324.22 | 19800000 | Keratin type I cytoskeletal 14 OS=Homo sapiens GN=KRT14 PE=1 SV=4 |
| | P98160 | PGBM_HUMAN | 333.53 | 19700000 | Basement membrane-specific heparan sulfate proteoglycan core protein OS=Homo sapiens GN=HSPG2 PE=1 SV=4 |
| | Q99715 | COCA1_HUMAN | 256.03 | 14900000 | Collagen alpha-1(XII) chain OS=Homo sapiens GN=COL12A1 PE=1 SV=2 |
| | P08174 | DAF_HUMAN | 165.96 | 10700000 | Complement decay-accelerating factor OS=Homo sapiens GN=CD55 PE=1 SV=4 |
| | P13647 | K2C5_HUMAN | 284.98 | 9250000 | Keratin type II cytoskeletal 5 OS=Homo sapiens GN=KRT5 PE=1 SV=3 |
| | P08779 | K1C16_HUMAN | 275.24 | 9230000 | Keratin type I cytoskeletal 16 OS=Homo sapiens GN=KRT16 PE=1 SV=4 |
| | O00468 | AGRIN_HUMAN | 233.5 | 8150000 | Agrin OS=Homo sapiens GN=AGRN PE=1 SV=5 |
| | P11047 | LAMC1_HUMAN | 215.83 | 6780000 | Laminin subunit gamma-1 OS=Homo sapiens GN=LAMC1 PE=1 SV=3 |
| | O43854 | EDIL3_HUMAN | 174.97 | 6670000 | EGF-like repeat and discoidin I-like domain-containing protein 3 OS=Homo sapiens GN=EDIL3 PE=1 SV=1 |
| | P60709 | ACTB_HUMAN | 207.85 | 6170000 | Actin cytoplasmic 1 OS=Homo sapiens GN=ACTB PE=1 SV=1 |
| | P63261 | ACTG_HUMAN | 207.85 | 6170000 | Actin cytoplasmic 2 OS=Homo sapiens GN=ACTG1 PE=1 SV=1 |
| | Q5T749 | KPRP_HUMAN | 157.97 | 5510000 | Keratinocyte proline-rich protein OS=Homo sapiens GN=KPRP PE=1 SV=1 |
| | P04003 | C4BPA_HUMAN | 158.03 | 4110000 | C4b-binding protein alpha chain OS=Homo sapiens GN=C4BPA PE=1 SV=2 |
| | P08865 | RSSA_HUMAN | 137.22 | 3360000 | 40S ribosomal protein SA OS=Homo sapiens GN=RPSA PE=1 SV=4 |
| | P15924 | DESP_HUMAN | 172.64 | 3340000 | Desmoplakin OS=Homo sapiens GN=DSP PE=1 SV=3 |
| | P14923 | PLAK_HUMAN | 160.32 | 2870000 | Junction plakoglobin OS=Homo sapiens GN=JUP PE=1 SV=3 |
| | P04406 | G3P_HUMAN | 147.47 | 2660000 | Glyceraldehyde-3-phosphate dehydrogenase OS=Homo sapiens GN=GAPDH PE=1 SV=3 |
| | P25391 | LAMA1_HUMAN | 170.48 | 2520000 | Laminin subunit alpha-1 OS=Homo sapiens GN=LAMA1 PE=1 SV=2 |
| | P05109 | S10A8_HUMAN | 107.5 | 2340000 | Protein S100-A8 OS=Homo sapiens GN=S100A8 PE=1 SV=1 |
| | P06702 | S10A9_HUMAN | 109.44 | 2320000 | Protein S100-A9 OS=Homo sapiens GN=S100A9 PE=1 SV=1 |
| | P0CG48 | UBC_HUMAN | 91.65 | 2250000 | Polyubiquitin-C OS=Homo sapiens GN=UBC PE=1 SV=3 |
| | P62979 | RS27A_HUMAN | 91.65 | 2250000 | Ubiquitin-40S ribosomal protein S27a OS=Homo sapiens GN=RPS27A PE=1 SV=2 |
| | P62987 | RL40_HUMAN | 91.65 | 2250000 | Ubiquitin-60S ribosomal protein L40 OS=Homo sapiens GN=UBA52 PE=1 SV=2 |
| | P35613 | BASI_HUMAN | 228.87 | 2250000 | Basigin OS=Homo sapiens GN=BSG PE=1 SV=2 |
| | Q86YZ3 | HORN_HUMAN | 172.12 | 2100000 | Hornerin OS=Homo sapiens GN=HRNR PE=1 SV=2 |

| Human from Mock CD9 exo (Mexo) | | | | | |
|---|---|---|---|---|---|
| | Accession # | Protein Abbreviation | -10lgP | Area CD9_mexo_3 reps | Description |
| | P07996 | TSP1_HUMAN | 221.73 | 1940000 | Thrombospondin-1 OS=Homo sapiens GN=THBS1 PE=1 SV=2 |
| | P07942 | LAMB1_HUMAN | 175.43 | 1890000 | Laminin subunit beta-1 OS=Homo sapiens GN=LAMB1 PE=1 SV=2 |
| | P23396 | RS3_HUMAN | 119.25 | 1790000 | 40S ribosomal protein S3 OS=Homo sapiens GN=RPS3 PE=1 SV=2 |
| | P05387 | RLA2_HUMAN | 108.01 | 1780000 | 60S acidic ribosomal protein P2 OS=Homo sapiens GN=RPLP2 PE=1 SV=1 |
| | Q04695 | K1C17_HUMAN | 263.72 | 1530000 | Keratin type I cytoskeletal 17 OS=Homo sapiens GN=KRT17 PE=1 SV=2 |
| | P07355 | ANXA2_HUMAN | 119.72 | 1410000 | Annexin A2 OS=Homo sapiens GN=ANXA2 PE=1 SV=2 |
| | Q7Z794 | K2C1B_HUMAN | 184.39 | 1160000 | Keratin type II cytoskeletal 1b OS=Homo sapiens GN=KRT77 PE=2 SV=3 |
| | Q14525 | KT33B_HUMAN | 207.47 | 1150000 | Keratin type I cuticular Ha3-II OS=Homo sapiens GN=KRT33B PE=1 SV=3 |
| | P10909 | CLUS_HUMAN | 93.11 | 1140000 | Clusterin OS=Homo sapiens GN=CLU PE=1 SV=1 |
| | P0CW18 | PRS56_HUMAN | 140.4 | 1130000 | Serine protease 56 OS=Homo sapiens GN=PRSS56 PE=1 SV=1 |
| | O15230 | LAMA5_HUMAN | 152 | 1130000 | Laminin subunit alpha-5 OS=Homo sapiens GN=LAMA5 PE=1 SV=8 |
| | P13987 | CD59_HUMAN | 109.03 | 976000 | CD59 glycoprotein OS=Homo sapiens GN=CD59 PE=1 SV=1 |
| | A0A140TA62 | A0A140TA62_HUMAN | 191.81 | 944000 | Uncharacterized protein OS=Homo sapiens GN=LOC100653049 PE=1 SV=1 |
| | P08195 | 4F2_HUMAN | 79.54 | 828000 | 4F2 cell-surface antigen heavy chain OS=Homo sapiens GN=SLC3A2 PE=1 SV=3 |
| | P81605 | DCD_HUMAN | 94.83 | 806000 | Dermcidin OS=Homo sapiens GN=DCD PE=1 SV=2 |
| | Q9Y4K0 | LOXL2_HUMAN | 110.7 | 702000 | Lysyl oxidase homolog 2 OS=Homo sapiens GN=LOXL2 PE=1 SV=1 |
| | P02794 | FRIH_HUMAN | 125.59 | 700000 | Ferritin heavy chain OS=Homo sapiens GN=FTH1 PE=1 SV=2 |
| | Q8N1N4 | K2C78_HUMAN | 149.66 | 672000 | Keratin type II cytoskeletal 78 OS=Homo sapiens GN=KRT78 PE=2 SV=2 |
| | Q02413 | DSG1_HUMAN | 118.87 | 658000 | Desmoglein-1 OS=Homo sapiens GN=DSG1 PE=1 SV=2 |
| | Q15818 | NPTX1_HUMAN | 96.27 | 557000 | Neuronal pentraxin-1 OS=Homo sapiens GN=NPTX1 PE=2 SV=2 |
| | P62701 | RS4X_HUMAN | 91.33 | 545000 | 40S ribosomal protein S4 X isoform OS=Homo sapiens GN=RPS4X PE=1 SV=2 |
| | P20908 | CO5A1_HUMAN | 83.82 | 516000 | Collagen alpha-1(V) chain OS=Homo sapiens GN=COL5A1 PE=1 SV=3 |
| | P23490 | LORI_HUMAN | 73.19 | 352000 | Loricrin OS=Homo sapiens GN=LOR PE=1 SV=2 |
| | Q08554 | DSC1_HUMAN | 71.67 | 239000 | Desmocollin-1 OS=Homo sapiens GN=DSC1 PE=1 SV=2 |
| | P62753 | RS6_HUMAN | 69.64 | 214000 | 40S ribosomal protein S6 OS=Homo sapiens GN=RPS6 PE=1 SV=1 |
| | P78386 | KRT85_HUMAN | 173.14 | 180000 | Keratin type II cuticular Hb5 OS=Homo sapiens GN=KRT85 PE=1 SV=1 |
| | P25398 | RS12_HUMAN | 70.75 | 157000 | 40S ribosomal protein S12 OS=Homo sapiens GN=RPS12 PE=1 SV=3 |
| | P62857 | RS28_HUMAN | 57.84 | 223000 | 40S ribosomal protein S28 OS=Homo sapiens GN=RPS28 PE=1 SV=1 |
| | | | | # ribosomal: | 9 |
| | | | | % ribosomal: | 0.155172414 |

| Gene ID | # Reads, PV-infected | #Reads, Mock-infected | RNA Type |
|---|---|---|---|
| tRNA-Gly-GGY_copy5 | 25474 | 8628 | tRNA_like |
| tRNA-Gly-GGG_copy19 | 25663 | 9223 | tRNA_like |
| tRNA-Lys-AAA_copy15 | 2177 | 1059 | tRNA_like |
| tRNA-Lys-AAA_copy26 | 6512 | 3209 | tRNA_like |
| tRNA-Lys-AAA_copy13 | 6510 | 3209 | tRNA_like |
| tRNA-Lys-AAA_copy17 | 6510 | 3209 | tRNA_like |
| tRNA-Lys-AAA_copy18 | 6510 | 3209 | tRNA_like |
| tRNA-Lys-AAA_copy20 | 6510 | 3209 | tRNA_like |
| tRNA-Lys-AAA_copy22 | 6510 | 3209 | tRNA_like |
| tRNA-Glu-GAG__copy60 | 1255 | 636 | tRNA_like |
| tRNA-Glu-GAG__copy61 | 1255 | 636 | tRNA_like |
| tRNA-Glu-GAG__copy62 | 1255 | 636 | tRNA_like |
| tRNA-Glu-GAG__copy63 | 1255 | 636 | tRNA_like |
| tRNA-Glu-GAG__copy64 | 1255 | 636 | tRNA_like |
| tRNA-Glu-GAG__copy65 | 1255 | 636 | tRNA_like |
| tRNA-Gly-GGG_copy22 | 1095 | 1302 | tRNA_like |
| chr16.tRNA18-GlyGCC | 25500 | 8640 | tRNA |
| chr16.tRNA25-GlyGCC | 26204 | 8936 | tRNA |
| chr16.tRNA24-GlyGCC | 26210 | 8943 | tRNA |
| chr16.tRNA19-GlyGCC | 26207 | 8943 | tRNA |
| chr17.tRNA5-GlyGCC | 26227 | 8950 | tRNA |
| chr2.tRNA19-GlyGCC | 26281 | 8977 | tRNA |
| chr1.tRNA68-GlyGCC | 26313 | 8992 | tRNA |
| chr6.tRNA128-GlyGCC | 26278 | 8995 | tRNA |
| chr1.tRNA133-GlyCCC | 30884 | 11541 | tRNA |
| chr1.tRNA4-GlyCCC | 30883 | 11541 | tRNA |
| chr21.tRNA2-GlyGCC | 19728 | 7526 | tRNA |
| chr1.tRNA35-GlyGCC | 19731 | 7528 | tRNA |
| chr1.tRNA37-GlyGCC | 19731 | 7528 | tRNA |
| chr1.tRNA39-GlyGCC | 19731 | 7528 | tRNA |
| chr1.tRNA41-GlyGCC | 19731 | 7528 | tRNA |
| chr1.tRNA3-PseudoCAC | 1036 | 407 | tRNA |
| chr6.tRNA71-LysTTT | 6510 | 3209 | tRNA |
| chr1.tRNA59-GluCTC | 1252 | 634 | tRNA |
| chr6.tRNA77-GluCTC | 1255 | 636 | tRNA |
| chr1.tRNA116-GluCTC | 1254 | 636 | tRNA |
| chr6.tRNA87-GluCTC | 1252 | 635 | tRNA |
| chr1.tRNA71-GluCTC | 1249 | 635 | tRNA |
| chr1.tRNA74-GluCTC | 1249 | 635 | tRNA |
| chr1.tRNA77-GluCTC | 1249 | 635 | tRNA |
| chr1.tRNA80-GluCTC | 1249 | 635 | tRNA |
| chr11.tRNA14-LysTTT | 1478 | 904 | tRNA |
| chr1.tRNA99-ValCAC | 3178 | 2451 | tRNA |
| chr1.tRNA91-PseudoCCC | 3637 | 3231 | tRNA |
| LSU-rRNA_Hsa_copy219 | 6815 | 2228 | rRNA |
| LSU-rRNA_Hsa_copy224 | 6815 | 2228 | rRNA |
| 5S_copy43 | 1881 | 902 | rRNA |
| 5S_copy44 | 1881 | 902 | rRNA |
| 5S_copy45 | 1881 | 902 | rRNA |
| 5S_copy46 | 1881 | 902 | rRNA |
| 5S_copy47 | 1881 | 902 | rRNA |
| 5S_copy48 | 1881 | 902 | rRNA |
| 5S_copy49 | 1881 | 902 | rRNA |
| 5S_copy50 | 1881 | 902 | rRNA |
| 5S_copy52 | 1881 | 902 | rRNA |
| 5S_copy53 | 1881 | 902 | rRNA |
| 5S_copy54 | 1881 | 902 | rRNA |
| 5S_copy55 | 1881 | 902 | rRNA |
| 5S_copy56 | 1881 | 902 | rRNA |
| 5S_copy57 | 1881 | 902 | rRNA |
| 5S_copy58 | 1881 | 902 | rRNA |
| 5S_copy59 | 1881 | 902 | rRNA |
| LSU-rRNA_Hsa_copy162 | 7428 | 3714 | rRNA |
| 5S_copy198 | 1461 | 735 | rRNA |
| 5S_copy51 | 1196 | 641 | rRNA |
| LSU-rRNA_Hsa_copy221 | 14276 | 8640 | rRNA |
| LSU-rRNA_Hsa_copy52 | 1503 | 1337 | rRNA |
| LSU-rRNA_Hsa_copy112 | 1272 | 1163 | rRNA |
| SSU-rRNA_Hsa_copy34 | 1788 | 1643 | rRNA |

| Gene ID | # Reads, PV-infected | #Reads, Mock-infected | RNA Type |
|---|---|---|---|
| SSU-rRNA_Hsa_copy35 | 1788 | 1643 | rRNA |
| snoMBII-202:RF00324.2 | 1390 | 144 | rfam |
| Y_RNA:RF00019.401 | 15911 | 2906 | rfam |
| Y_RNA:RF00019.402 | 7172 | 1860 | rfam |
| snoZ40:RF00342.2 | 8003 | 2817 | rfam |
| snoU83B:RF00593.3 | 2241 | 826 | rfam |
| snoU105B:RF01173.1 | 5820 | 2815 | rfam |
| U2:RF00004.6 | 1182 | 576 | rfam |
| U2:RF00004.59 | 2261 | 1125 | rfam |
| U2:RF00004.51 | 1043 | 537 | rfam |
| U2:RF00004.75 | 1406 | 726 | rfam |
| Y_RNA:RF00019.404 | 3052 | 1633 | rfam |
| snoZ17:RF00266.1 | 2742 | 1502 | rfam |
| U2:RF00004.61 | 1287 | 750 | rfam |
| snoZ39:RF00341.1 | 1827 | 1129 | rfam |
| snosnR60_Z15:RF00309.1 | 109517 | 69531 | rfam |
| RNase_MRP:RF00030.1 | 1330 | 905 | rfam |
| snoZ17:RF00266.2 | 10712 | 8772 | rfam |
| VAC141 | 51729 | 17594 | RefSeq_antisense |
| COLQ1 | 25690 | 9240 | RefSeq_antisense |
| LOC100129636:copy41 | 46022 | 19915 | RefSeq_antisense |
| FAM69A1 | 8115 | 3687 | RefSeq_antisense |
| BTRC1 | 2458 | 1211 | RefSeq_antisense |
| LOC1002881421 | 3469 | 1794 | RefSeq_antisense |
| NBPF101 | 3043 | 1576 | RefSeq_antisense |
| NBPF24:copy21 | 1685 | 888 | RefSeq_antisense |
| ZNF5211 | 1305 | 830 | RefSeq_antisense |
| SAMD4A1 | 1348 | 916 | RefSeq_antisense |
| ZNF2341 | 10390 | 7824 | RefSeq_antisense |
| CNTNAP21 | 1500 | 1291 | RefSeq_antisense |
| BCAS31 | 1234 | 1163 | RefSeq_antisense |
| ZNF8601 | 11780 | 11148 | RefSeq_antisense |
| TTC281 | 1005 | 1060 | RefSeq_antisense |
| HPSE21 | 1593 | 1740 | RefSeq_antisense |
| SMARCC11 | 1552 | 1727 | RefSeq_antisense |
| ANK11 | 5849 | 6526 | RefSeq_antisense |
| CELSR11 | 1455 | 1681 | RefSeq_antisense |
| TSHZ21 | 1031 | 1194 | RefSeq_antisense |
| ZNF6951 | 1414 | 1669 | RefSeq_antisense |
| MGAT4C1 | 1424 | 1688 | RefSeq_antisense |
| ABCC61 | 1380 | 1660 | RefSeq_antisense |
| KIAA13241 | 1366 | 1666 | RefSeq_antisense |
| OSBPL61 | 1348 | 1645 | RefSeq_antisense |
| SLC2A131 | 1343 | 1655 | RefSeq_antisense |
| RGSL11 | 1331 | 1643 | RefSeq_antisense |
| PTP4A21 | 1400 | 1732 | RefSeq_antisense |
| OSBPL6:copy21 | 1291 | 1620 | RefSeq_antisense |
| ACAN1 | 1288 | 1623 | RefSeq_antisense |
| CHRNA31 | 1285 | 1621 | RefSeq_antisense |
| IGF11 | 1287 | 1624 | RefSeq_antisense |
| IGF1:copy21 | 1281 | 1620 | RefSeq_antisense |
| ASPM1 | 1279 | 1623 | RefSeq_antisense |
| SF3A21 | 1268 | 1614 | RefSeq_antisense |
| FOXO4 | 1266 | 1614 | RefSeq_antisense |
| FBXO111 | 1967 | 2540 | RefSeq_antisense |
| RPL13 | 1393 | 146 | RefSeq |
| MTRNR2L2 | 3134 | 334 | RefSeq |
| MTRNR2L8 | 3239 | 488 | RefSeq |
| DHFR | 4888 | 753 | RefSeq |
| SNRPB | 9720 | 2437 | RefSeq |
| GNB2L1 | 15882 | 4102 | RefSeq |
| EEF2 | 3719 | 1108 | RefSeq |
| C6orf48:copy5 | 10673 | 3630 | RefSeq |
| VAC14 | 52443 | 17895 | RefSeq |
| MTRNR2L1 | 1629 | 574 | RefSeq |
| PCBD2 | 1502 | 535 | RefSeq |
| C6orf48 | 12544 | 4470 | RefSeq |
| C6orf48:copy2 | 12544 | 4470 | RefSeq |
| C6orf48:copy3 | 12544 | 4470 | RefSeq |
| C6orf48:copy4 | 12544 | 4470 | RefSeq |

| Gene ID | # Reads, PV-infected | #Reads, Mock-infected | RNA Type |
|---|---|---|---|
| EIF5 | 1211 | 450 | RefSeq |
| RPL3 | 29372 | 11149 | RefSeq |
| RANBP1 | 5705 | 2211 | RefSeq |
| RANBP1:copy2 | 5705 | 2211 | RefSeq |
| RANBP1:copy3 | 5705 | 2211 | RefSeq |
| RPS3A | 2728 | 1203 | RefSeq |
| RPL13A | 10873 | 4892 | RefSeq |
| RPL5 | 8031 | 3629 | RefSeq |
| TCP1 | 4297 | 1965 | RefSeq |
| RPL7A | 5863 | 2698 | RefSeq |
| MED24 | 1395 | 652 | RefSeq |
| RBMX | 11941 | 5650 | RefSeq |
| PPAN | 6234 | 2963 | RefSeq |
| NBPF24:copy2 | 1281 | 616 | RefSeq |
| UBAP2 | 3829 | 1940 | RefSeq |
| GNL3 | 12121 | 6178 | RefSeq |
| RPS12 | 73495 | 37697 | RefSeq |
| PUM1 | 3162 | 1624 | RefSeq |
| CCNB1IP1 | 1789 | 919 | RefSeq |
| RPL4 | 6833 | 3567 | RefSeq |
| RPL17 | 73107 | 38424 | RefSeq |
| C18orf32 | 73109 | 38426 | RefSeq |
| DDX39B:copy7 | 1096 | 593 | RefSeq |
| DDX39B | 1100 | 597 | RefSeq |
| DDX39B:copy2 | 1100 | 597 | RefSeq |
| DDX39B:copy5 | 1100 | 597 | RefSeq |
| DDX39B:copy6 | 1100 | 597 | RefSeq |
| DDX39B:copy4 | 1099 | 597 | RefSeq |
| DDX39B:copy8 | 1099 | 597 | RefSeq |
| LOC100129636:copy4 | 32138 | 17588 | RefSeq |
| NOP58 | 2867 | 1595 | RefSeq |
| SF3B3 | 4268 | 2389 | RefSeq |
| PRPF39 | 8077 | 4560 | RefSeq |
| RPS8 | 31612 | 18822 | RefSeq |
| COX7C | 1832 | 1132 | RefSeq |
| EIF4A2 | 185278 | 115721 | RefSeq |
| NCL | 54231 | 34077 | RefSeq |
| WDR43 | 28171 | 17865 | RefSeq |
| RPL21 | 32459 | 20754 | RefSeq |
| RPL37 | 4082 | 2732 | RefSeq |
| ATP5B | 76290 | 51250 | RefSeq |
| TARDBP | 1463 | 1008 | RefSeq |
| RPL23A | 19972 | 13836 | RefSeq |
| STK39 | 10715 | 8010 | RefSeq |
| HSPA9 | 184891 | 143673 | RefSeq |
| KIAA0825:copy2 | 89892 | 70589 | RefSeq |
| KIAA0825 | 90099 | 70756 | RefSeq |
| MCM7 | 1020 | 811 | RefSeq |
| RC3H2 | 6072 | 4900 | RefSeq |
| LOC100288142 | 2200 | 1777 | RefSeq |
| TSR1 | 23192 | 18939 | RefSeq |
| SORCS2 | 1569 | 1368 | RefSeq |
| PANK2 | 1095 | 985 | RefSeq |
| PANK3 | 1030 | 947 | RefSeq |
| PPARGC1B | 11841 | 11161 | RefSeq |
| NBPF10 | 1601 | 1555 | RefSeq |
| ANK1 | 5708 | 6320 | RefSeq |
| PI4KA | 1480 | 1714 | RefSeq |
| SHISA9 | 1403 | 1675 | RefSeq |
| EBF1 | 1332 | 1641 | RefSeq |
| MFHAS1 | 1330 | 1640 | RefSeq |
| TKTL1 | 1318 | 1634 | RefSeq |
| JAM3 | 1356 | 1687 | RefSeq |
| PTPLB | 1317 | 1640 | RefSeq |
| AHSA1 | 1285 | 1624 | RefSeq |
| SPATA6 | 1236 | 1572 | RefSeq |
| PLEKHG4B | 1272 | 1618 | RefSeq |
| NADKD1 | 1283 | 1634 | RefSeq |
| MSTO1 | 1328 | 1720 | RefSeq |
| piR-36056 | 1293 | 126 | piRNA |

| Gene ID | # Reads, PV-infected | #Reads, Mock-infected | RNA Type |
|---|---|---|---|
| piR-33783 | 7106 | 699 | piRNA |
| piR-31703 | 1389 | 144 | piRNA |
| piR-44892 | 6896 | 1047 | piRNA |
| piR-31636 | 2345 | 394 | piRNA |
| piR-31986 | 2346 | 396 | piRNA |
| piR-31987 | 2346 | 396 | piRNA |
| piR-34871 | 8430 | 1478 | piRNA |
| piR-34342 | 2910 | 789 | piRNA |
| piR-58469 | 1150 | 329 | piRNA |
| piR-47305 | 1149 | 329 | piRNA |
| piR-30113 | 4239 | 1270 | piRNA |
| piR-32376 | 28000 | 9039 | piRNA |
| piR-32377 | 28000 | 9039 | piRNA |
| piR-33856 | 5643 | 1905 | piRNA |
| piR-30112 | 7998 | 2812 | piRNA |
| piR-31447 | 7902 | 2782 | piRNA |
| piR-34531 | 1179 | 422 | piRNA |
| piR-34532 | 1179 | 422 | piRNA |
| piR-34533 | 1179 | 422 | piRNA |
| piR-34534 | 1179 | 422 | piRNA |
| piR-34535 | 1179 | 422 | piRNA |
| piR-34536 | 1179 | 422 | piRNA |
| piR-35551 | 1419 | 514 | piRNA |
| piR-35550 | 1419 | 514 | piRNA |
| piR-35549 | 1419 | 514 | piRNA |
| piR-61135 | 1165 | 422 | piRNA |
| piR-52729 | 2259 | 827 | piRNA |
| piR-30799 | 25316 | 9735 | piRNA |
| piR-33879 | 20913 | 8497 | piRNA |
| piR-31531 | 2185 | 1096 | piRNA |
| piR-36711 | 2827 | 1446 | piRNA |
| piR-36712 | 2827 | 1446 | piRNA |
| piR-60238 | 2825 | 1446 | piRNA |
| piR-31623 | 16458 | 9006 | piRNA |
| piR-31970 | 2695 | 1477 | piRNA |
| piR-30810 | 5673 | 3159 | piRNA |
| piR-35059 | 10684 | 6425 | piRNA |
| piR-61298 | 46516 | 28008 | piRNA |
| piR-32648 | 1816 | 1126 | piRNA |
| piR-30652 | 52878 | 33107 | piRNA |
| piR-45371 | 23108 | 14672 | piRNA |
| piR-33748 | 23116 | 14678 | piRNA |
| piR-30840 | 183284 | 142537 | piRNA |
| piR-57125 | 88946 | 69953 | piRNA |
| piR-33685 | 21850 | 18131 | piRNA |
| piR-33686 | 21850 | 18131 | piRNA |
| piR-33687 | 21850 | 18131 | piRNA |
| RHOU:antisense | 1950 | 930 | other_ncRNA_antisense |
| LOC285692:antisense | 1543 | 1867 | other_ncRNA_antisense |
| LOC400654:antisense | 1363 | 1736 | other_ncRNA_antisense |
| SNHG5 | 3351 | 974 | other_ncRNA |
| RABGGTB | 36455 | 11121 | other_ncRNA |
| GAS5 | 390137 | 175449 | other_ncRNA |
| C17orf76-AS1 | 21881 | 11319 | other_ncRNA |
| SNHG12 | 16626 | 8893 | other_ncRNA |
| THAP9-AS1 | 1032 | 558 | other_ncRNA |
| SNHG1 | 1000263 | 552939 | other_ncRNA |
| NOP56 | 15782 | 8780 | other_ncRNA |
| RNA45S5 | 35336 | 20417 | other_ncRNA |
| RNA5-8S5 | 40157 | 23358 | other_ncRNA |
| RNA5-8S5:copy2 | 4805 | 2936 | other_ncRNA |
| RMRP | 1330 | 905 | other_ncRNA |
| ZFAS1 | 485527 | 341950 | other_ncRNA |
| LOC285441 | 1364 | 1666 | other_ncRNA |
| PI4KAP2 | 1326 | 1637 | other_ncRNA |
| LOC284294 | 1423 | 1775 | other_ncRNA |
| PI4KAP1 | 1297 | 1621 | other_ncRNA |
| MSTO2P | 1330 | 1720 | other_ncRNA |
| hsa-miR-3607-5p | 1820 | 1122 | miRNA |
| hsa-mir-3607 | 1827 | 1129 | miRNA |

| Gene ID | # Reads, PV-infected | #Reads, Mock-infected | RNA Type |
|---|---|---|---|
| hsa-miR-3607-3p | 1815 | 1124 | miRNA |
| hsa-miR-378c | 1146 | 726 | miRNA |
| hsa-mir-378c | 1146 | 726 | miRNA |
| hsa-miR-92a-3p | 1415 | 932 | miRNA |
| hsa-mir-92a-2 | 1415 | 932 | miRNA |
| hsa-mir-18a | 1089 | 725 | miRNA |
| hsa-mir-92a-1 | 2656 | 1882 | miRNA |
| hsa-mir-21 | 7389 | 5277 | miRNA |
| hsa-miR-21-5p | 7365 | 5267 | miRNA |
| hsa-mir-17 | 1687 | 1313 | miRNA |
| hsa-miR-106a-5p | 1437 | 1134 | miRNA |
| hsa-mir-106a | 1437 | 1134 | miRNA |
| hsa-miR-17-5p | 1513 | 1201 | miRNA |
| hsa-miR-20a-5p | 2232 | 1897 | miRNA |
| hsa-mir-20a | 2365 | 2011 | miRNA |
| hsa-mir-103a-2 | 1049 | 963 | miRNA |
| hsa-mir-103a-1 | 1016 | 944 | miRNA |
| hsa-miR-103a-3p | 1016 | 944 | miRNA |
| hsa-mir-378a | 11783 | 11145 | miRNA |
| hsa-miR-378a-3p | 11769 | 11136 | miRNA |
| hsa-mir-19b-1 | 3980 | 3946 | miRNA |
| hsa-miR-19b-3p | 2728 | 3002 | miRNA |
| hsa-mir-19b-2 | 2728 | 3002 | miRNA |
| hsa-mir-19a | 1944 | 2168 | miRNA |
| hsa-mir-486 | 5559 | 6237 | miRNA |
| hsa-miR-486-5p | 5544 | 6227 | miRNA |
| hsa-miR-19a-3p | 1897 | 2144 | miRNA |
| linc-EPHA6-1:copy3:antisense | 2168 | 241 | lincRNA_antisense |
| linc-EPHA6-1:copy2:antisense | 2356 | 266 | lincRNA_antisense |
| linc-CLLU1-7:antisense | 1302 | 1623 | lincRNA_antisense |
| linc-ANGEL2-2:antisense | 1278 | 1619 | lincRNA_antisense |
| linc-EHD3-1:antisense | 1219 | 1570 | lincRNA_antisense |
| linc-DHFRL1-4:copy4 | 2124 | 232 | lincRNA |
| linc-DHFRL1-4:copy2 | 1765 | 195 | lincRNA |
| linc-DHFRL1-4:copy3 | 2168 | 241 | lincRNA |
| linc-DHFRL1-4 | 2356 | 266 | lincRNA |
| XLOC_014515 | 35341 | 20419 | lincRNA |
| linc-CPXM2-2 | 1271 | 1620 | lincRNA |
| linc-C5orf38-3 | 1267 | 1616 | lincRNA |
| ACA16 | 1574 | 548 | HAcaBox |
| ACA28 | 1201 | 440 | HAcaBox |
| ACA20 | 4277 | 1949 | HAcaBox |
| HBII-202 | 1390 | 144 | CDBox |
| U81 | 80589 | 18460 | CDBox |
| U78 | 20865 | 4822 | CDBox |
| SNORD119 | 9718 | 2437 | CDBox |
| U96a | 15868 | 4095 | CDBox |
| U45A | 23616 | 6668 | CDBox |
| U50 | 3307 | 948 | CDBox |
| U37 | 3712 | 1106 | CDBox |
| mgh28S-2409 | 69314 | 23338 | CDBox |
| U58A | 5649 | 1906 | CDBox |
| U52 | 10634 | 3603 | CDBox |
| U29 | 191712 | 65587 | CDBox |
| U45C | 12386 | 4316 | CDBox |
| mgh28S-2411 | 8003 | 2817 | CDBox |
| U24 | 2263 | 832 | CDBox |
| U83A | 2241 | 826 | CDBox |
| U43 | 26438 | 10178 | CDBox |
| U44 | 95529 | 37364 | CDBox |
| U58B | 20917 | 8498 | CDBox |
| HBII-135 | 2998 | 1223 | CDBox |
| U47 | 2433 | 999 | CDBox |
| U26 | 9180 | 3777 | CDBox |
| U16 | 2477 | 1022 | CDBox |
| U73a | 2691 | 1185 | CDBox |
| SNORD121A | 1343 | 601 | CDBox |
| U21 | 7910 | 3581 | CDBox |
| U18A | 2074 | 940 | CDBox |
| U49B | 1907 | 866 | CDBox |

| Gene ID | # Reads, PV-infected | #Reads, Mock-infected | RNA Type |
|---|---|---|---|
| U56 | 4630 | 2115 | CDBox |
| U101 | 50118 | 22897 | CDBox |
| U25 | 257666 | 117849 | CDBox |
| U104 | 9961 | 4622 | CDBox |
| U32A | 9832 | 4593 | CDBox |
| SNORD124 | 1325 | 623 | CDBox |
| U38A | 4965 | 2343 | CDBox |
| U61 | 11907 | 5625 | CDBox |
| U36B | 2249 | 1076 | CDBox |
| U105B | 5820 | 2815 | CDBox |
| U75 | 1598 | 791 | CDBox |
| U60 | 3474 | 1725 | CDBox |
| U38B | 2210 | 1109 | CDBox |
| U76 | 6507 | 3300 | CDBox |
| HBII-251 | 2852 | 1461 | CDBox |
| HBII-210 | 5541 | 2870 | CDBox |
| SNORD126 | 1698 | 883 | CDBox |
| HBII-234 | 1418 | 757 | CDBox |
| U77 | 58987 | 31884 | CDBox |
| SNORD121B | 2331 | 1265 | CDBox |
| U49A | 16948 | 9213 | CDBox |
| mgh18S-121 | 2742 | 1502 | CDBox |
| HBII-82 | 3819 | 2095 | CDBox |
| HBII-108B | 1112 | 612 | CDBox |
| U42B | 5693 | 3169 | CDBox |
| HBII-420 | 14577 | 8193 | CDBox |
| SNORD127 | 8058 | 4549 | CDBox |
| HBII-296B | 1108 | 628 | CDBox |
| U79 | 13207 | 7908 | CDBox |
| HBII-55 | 10803 | 6497 | CDBox |
| U58C | 46516 | 28008 | CDBox |
| U106 | 22860 | 14047 | CDBox |
| U59A | 21143 | 13141 | CDBox |
| snR39B | 184585 | 115378 | CDBox |
| U82 | 53154 | 33278 | CDBox |
| U46 | 24413 | 15364 | CDBox |
| U80 | 109517 | 69531 | CDBox |
| HBII-429 | 23154 | 14701 | CDBox |
| U102 | 32453 | 20749 | CDBox |
| U27 | 21914 | 14244 | CDBox |
| HBII-240 | 4082 | 2732 | CDBox |
| U30 | 518520 | 350943 | CDBox |
| U59B | 55126 | 38095 | CDBox |
| HBII-99 | 28498 | 19982 | CDBox |
| U18C | 1856 | 1304 | CDBox |
| HBII-99B | 434163 | 307916 | CDBox |
| U63 | 184414 | 143325 | CDBox |
| HBII-295 | 6005 | 4874 | CDBox |
| Z17B | 10712 | 8772 | CDBox |
| HBII-296A | 22064 | 18306 | CDBox |
